# Self-Organizing Neural Networks in Novel Moving Bodies: Anatomical, Behavioral, and Transcriptional Characterization of a Living Construct with a Nervous System

**DOI:** 10.1101/2025.04.14.648732

**Authors:** Haleh Fotowat, Laurie O’Neill, Léo Pio-Lopez, Megan Sperry, Patrick Erickson, Tiffany Lin, Michael Levin

## Abstract

A great deal is known about the formation and architecture of biological neural networks in animal models, which have arrived at their current structure-function relationship through evolution by natural selection. Little is known about the development of such structure-function relationships in a scenario where neurons are allowed to grow within evolutionarily-novel, motile bodies. Previous work showed that when a piece of ectodermal tissue is excised from *Xenopus* embryos and allowed to develop *ex vivo*, it will develop into a three-dimensional (3D) mucociliary organoid, and exhibits behaviors different from those observed in tadpoles of the same age. These ‘biological robots’ or ‘biobots’ are autonomous, self-powered, and able to move through aqueous environments. Here we report a novel type of biobot that is composed of ciliated epidermis and additionally incorporates neural tissue (neurobots). We show that neural precursor cells implanted within the *Xenopus* skin constructs develop into mature neurons and extend processes towards the outer surface of the bot as well as among each other. These self-organized neurobots show distinct external morphology, generate more complex patterns of spontaneous movements, and are differentially affected by neuroactive drugs compared to their non-neuronal counterparts. Calcium imaging experiments show that neurons within neurobots are indeed active. Transcriptomics analysis of the neurobots reveals increased variability of transcript profiles, expression of a plethora of genes relating to nervous system development and function, a shift toward more ancient genes, and up-regulation of neuronal genes implicated in visual perception.

## Introduction

Sensing cues from the environment and translating them into appropriate responses is the fundamental function of the nervous system in all animals. Critically, nervous systems endow animals with the ability to generate context- and experience-dependent changes in their behavior. Nervous systems are also known to be plastic and adapt both structurally and functionally, on a much shorter timescale, to changes in sensory and/or motor effectors that might occur in the lifetime of an organism, e.g. as a result of injury, amputation, or sensory deprivation, albeit in an age dependent manner^1–4^. The degree to which the nervous system of an animal can adapt to a new body plan is especially impressive in cases where changes in sensory-motor architecture is particularly drastic. For example, ectopically induced eyes in the tail of *Xenopus* tadpoles were shown to confer light sensitivity to the eyeless hosts ^5^.

What are the limits of neuroplasticity of nervous systems? How would wild-type neurons establish pattern and function in a new embodiment? Creating truly novel configurations of biological material allows us to probe the plasticity of evolutionary hardware to adapt on developmental (not evolutionary) timescales to truly *novel* circumstances and has applications for regenerative medicine, human augmentation, and biological engineering. We sought to establish a model system in which we could study the morphology and function of neural networks in novel scenarios, better understand the evolutionary developmental biology of the nervous system, and improve the design of future innervated constructs.

In the past, neuromuscular assembloids have been used to model cortico-spinal motor functions^6^, and neural cultures have been interfaced with computers or robots using micro electrode arrays to learn and perform various tasks ^7–10^. These hybrid robots^11,12^ are not built exclusively using biological tissue, are not fully embodied, and are not self-motile. Motile biohybrid soft robots have been built mostly through engineering patterned muscle tissues and propelled through external controls, e.g. electrical or optical stimulation ^13,14^. Biohybrid bots that move through neuromuscular actuation have also been built by embedding neural and muscle cells in silicone polymer scaffolds^15,16^. Similarly, such bots do not self-assemble, require external stimulation for propulsion, can only move at very slow speeds, have a very limited behavioral repertoire, and still require synthetic scaffolds.

When ectodermal tissue is excised from the animal pole of a late blastula stage *Xenopus* embryo, and allowed to develop *ex vivo*, it will develop into a self-motile, self-powered 3D mucociliary organoid^17^. These biobots express the four cell types normally present in a tadpole skin. These include multiciliated cells (MCCs), mucus secreting goblet cells, ionocytes that regulate ionic homeostasis of epidermis, and small secretory cells (SSCs) ^18,19^. MCCs act as motor effectors in these organoids via flow arising from their polarized beating, generating a suite of stereotyped movement trajectories and velocities. Interestingly, SSCs were shown to secrete serotonin, which stimulates an increase in ciliary beating frequency through serotonergic receptors expressed on MCCs ^20^. These self-motile mucociliary organoids, which we will refer to herein as biobots, are capable of navigating aqueous environments, generating a suite of stereotyped movement trajectories and velocities.

In this study we asked what would happen if we provided these biobots with the ‘‘raw materials’’ for building a nervous system? That is, if we used neural precursor cells in addition to embryonic ectodermal cells, would such neural precursor cells indeed differentiate into functional neurons within the biobot? What genes would they express? How would the behavior and morphology of these ‘‘neurobots’’ compare to their non-neuronal counterparts?

We show that neural precursor cells harvested from *Xenopus* embryos and implanted in biobots made from *Xenopus* ectodermal cells indeed differentiate into functional neurons and extend their processes within and toward the neurobot’s outer surface. We show that neurobots exhibit significant differences in behavior and anatomy compared to their non-neuronal counterparts. Transcriptomic characterization of neurobots reveals significant upregulation of genes involved in nervous system development, compared to non-neuronal biobots ^21–24^, including those important for processing light stimuli.

## Results

### Neurobots can be constructed by implanting exogenous neural precursors into Xenopus ectodermal explants

To characterize the structure and function of a nervous system that self-assembles in a novel embodiment, we established an experimental procedure for implanting biobots with neural precursor cells during the first few minutes of their formation. As shown previously, biobots can be constructed by excising tissue from the animal hemisphere of a Nieuwkoop and Faber stage 9 *Xenopus laevis* embryo (animal cap). In the course of 30 minutes, the excised tissue will gradually heal, initially forming a ‘‘bowl’’ shape before forming a closed spherical shape ^17^. We used the time window prior to the closure of the tissue to insert neuronal precursor cells inside the healing tissue (Fig. 1ai, 1aii-top left panel).

**Figure 1.**
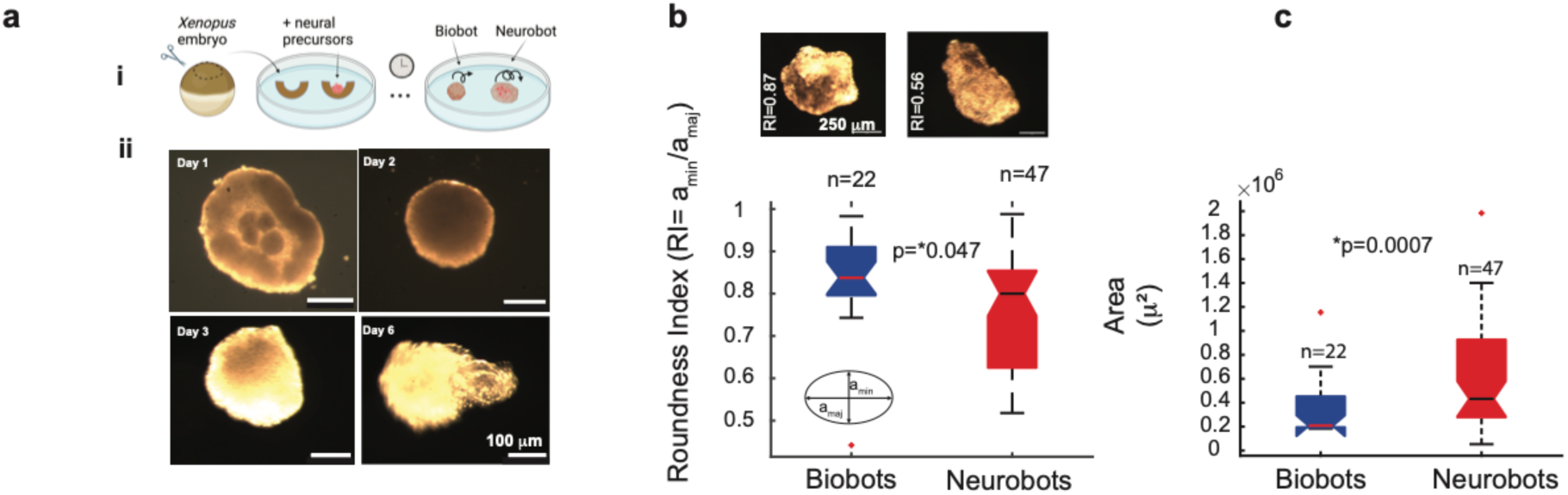
Construction and development of a neurobot. **a**. Neural precursor clumps were placed in the center of an animal cap ‘cup’, excised from the animal pole of a *Xenopus laevis* embryo before it fully closes up as it heals. The composite forms gradually into first a sphere and then a more elongated shape which is mobile by day 3. **b.** Examples of two neurobots, one more round than the other. **c.** Roundness Index (RI) was calculated by fitting an ellipse on the image of the bot and calculating the ratio between the minor and major axes. Neurobots tended to be less round than biobots. Non-parametric Kruskal-Wallis test was used to calculate statistical significance.

To obtain neural cells, we took advantage of the fact that if the animal cap is excised and dissociated at the late blastula to early gastrula stage, and the dissociated cells are allowed to remain separated for 3 hours or more, they will assume a neural fate^25,26^. To obtain aggregates of neural precursors, we dissociated animal caps from approximately 50 embryos, allowed them to remain separated for 3 hours, reaggregated the cells, then placed clumps of reaggregated cells inside a freshly excised animal cap before it fully healed into a spherical shape (Fig. 1ai, 1aii, top left panel, see Methods). Within 30 minutes the formed composite assumes a spherical shape, and by the second day, it is fully healed (Fig. 1aii, top right panel). As with non-neuronal biobots, by the third day, multiciliated cells start appearing on their outer surface and the bots start moving around in the dish. Similar to biobots, neurobots have a lifespan of about 9-10 days without being fed, and survive by consuming maternal yolk platelet present in all early *Xenopus* embryonic tissue^17^. Interestingly, by day 6 neurobots tend to have a more elongated shape than biobots, and they become significantly larger (Fig. 1b,c).

To investigate whether the difference in size and elongation is simply due to implanting the animal caps with additional cells, we generated a third type of bot (sham neurobots) in a manner similar to neurobots, except that the implanted cells were not allowed to remain separated for 3 hours. Instead, they were reaggregated shortly after dissociation (within 30 minutes) to prevent the induction of neural fate. We found that the sham neurobots were not elongated and did not show a significant size difference compared to biobots (Supp. Fig. 1a,b). These results suggest that the elongation and increase in size may be due to neuronal growth within neurobots.

### Implanted neural precursor cells differentiate into neurons, making projections within the neurobot as well as toward the cells lining the outer surface

To determine whether the implanted cells had indeed differentiated into neurons, we fixed and stained the neurobots with an antibody that specifically binds to acetylated alpha tubulin, which is abundantly present in neurons and multiciliated cells (see Methods; neurons and multiciliated cells are readily distinguishable from each other due to their distinctive morphology). We found that the implanted neural precursor cells indeed differentiate into neurons (Fig. 2). The neurons extend their processes not only within the neurobot, but also toward the outer surface (see arrows in Fig. 2c,d). Such projections toward the cells lining the surface of the bot suggest the possibility of neurons modulating the activity of surface effectors including multiciliated cells and/or the activity of those that modulate the ciliary beating frequency, e.g. small secretory cells.

**Figure 2.**
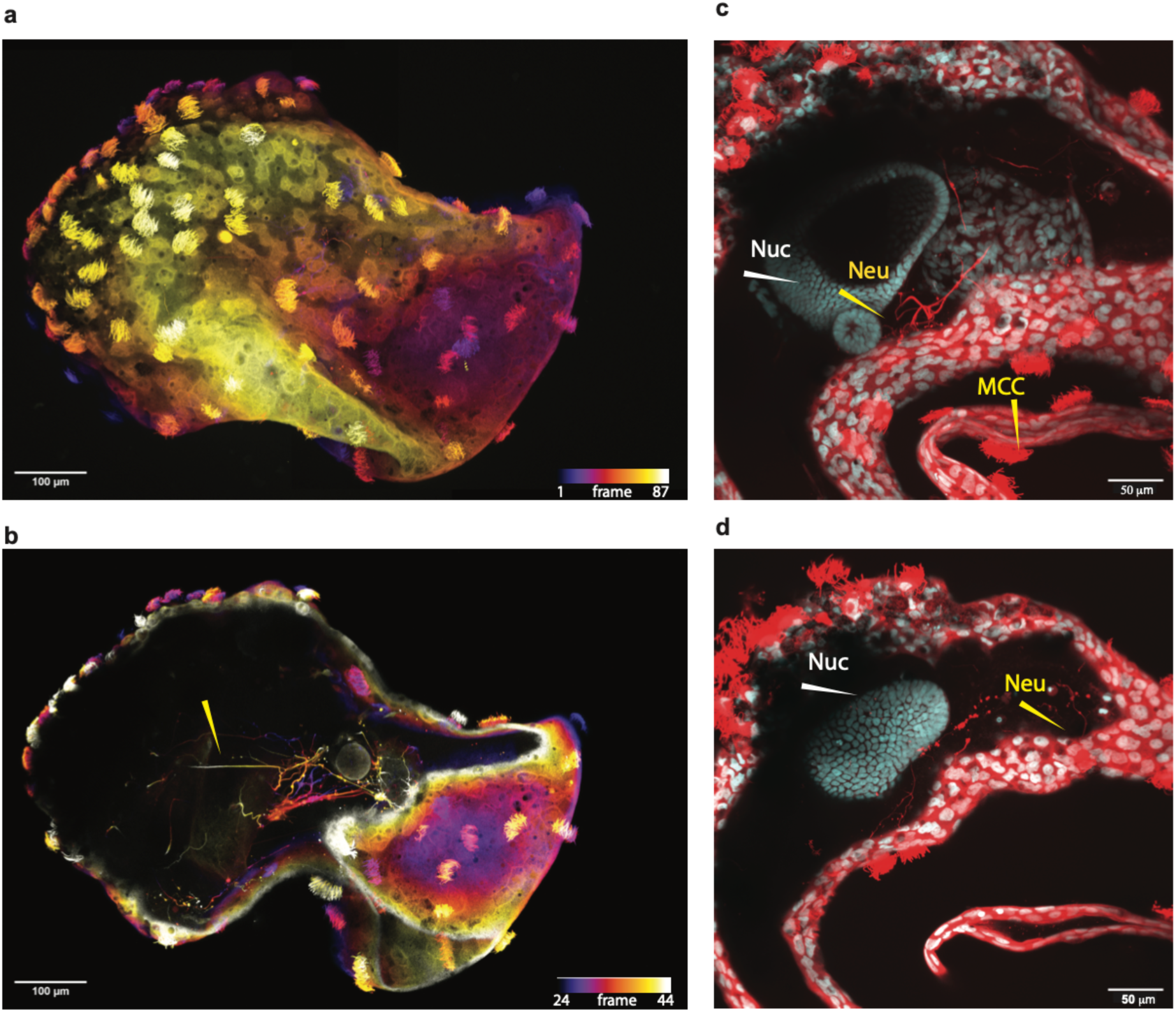
Implanted neural precursors develop into neurons and extend their processes throughout the neurobot. **a-b.** Z-projection of confocal image stack of a neurobot labeled with acetylated alpha tubulin which stains neurons and cilia of the multiciliated cells. **b** is the staining of the same neurobot as shown in **a** with fewer projected planes, rendering neural processes inside the bot visible. Color code corresponds to depth within the bot (confocal plane number). **c-d** Subregions of same neurobot, showing neural processes projecting toward surface cells. Red shows the acetylated alpha tubulin stain (AAT) and cyan depicts a nuclear (Hoechst) co-label (NUC). Yellow arrows point to neural processes (Neu) or multiciliated cells whose cilia are stained (MCC). White arrows point to nuclear staining (Nuc).

There was a large degree of variability in the structure of sprouting in different neurobots. No two neurobots showed identical neural architecture (Supp. Fig. 2). This is not surprising given the variability of initial conditions resulting from their manual construction and the inevitable variability of the amount of implanted tissue. This variability, however, allowed us to investigate correlations between various physical and behavioral characteristics of neurobots, which we will discuss in more detail in the following sections. Despite these variabilities, we found that neural processes in most neurobots tended to emanate from one or more nuclear regions (labeled using a nuclear stain), presumably corresponding to the implanted clumps of cells (Fig. 1a Fig. 2c,d, Supp. Fig. 2). Interestingly, these regions were often surrounded by regions with seemingly no nuclear staining (Fig. 2b-d, Supp. Fig. 2). The presence of these seemingly empty spaces is intriguing, and we hypothesize that these spaces may be composed of neural support structures, such as the extracellular matrix (ECM). Support for this hypothesis includes cases in which we observed neurites traversing long distances in this empty space along a straight line (Fig. 2b, yellow arrow). Additional support for this hypothesis comes from the significant upregulation in genes encoding various proteins associated with the ECM, which we will present later in this manuscript. Future experiments are needed to fully characterize the nature of this internal space.

### Neurobot neurons are functional

To assess whether neurons within neurobots are functional, we built neurobots using neural cells extracted from embryos with genetically encoded calcium indicators (GCaMP6s, see Methods); this tool is commonly used to study neural activity ^27^. Figure 3 shows an example of calcium signals measured in a freely moving neurobot, which was circling around in the dish (Supp. Video 1). Motion corrected videos (see Methods, Supp. Video 2) were then analyzed using an open-source software (suite2p) to extract the fluorescent activity. We found that the implanted cells did show calcium activity in all recorded neurobots. We occasionally observed synchronized activity in nearby or distant regions of interest, suggesting presence of connectivity among implanted neurons (see e.g. arrowheads in Fig. 3c). Measuring neural signals in freely moving neurobots however, proved to be challenging as it was not possible to keep individual regions of interest in focus over long periods of time. Moreover, since every neurobot showed a different pattern of neural expression (Supp. Fig. 2), findings in one neurobot could not be reproduced and confirmed in others. Developing methods for creating consistent neural expression and testing neurobots in experimental setups where a change in course of movement can be observed repeatedly (e.g. a maze) are critical for assessing the presence of correlations between neural activity and behavior.

**Figure 3.**
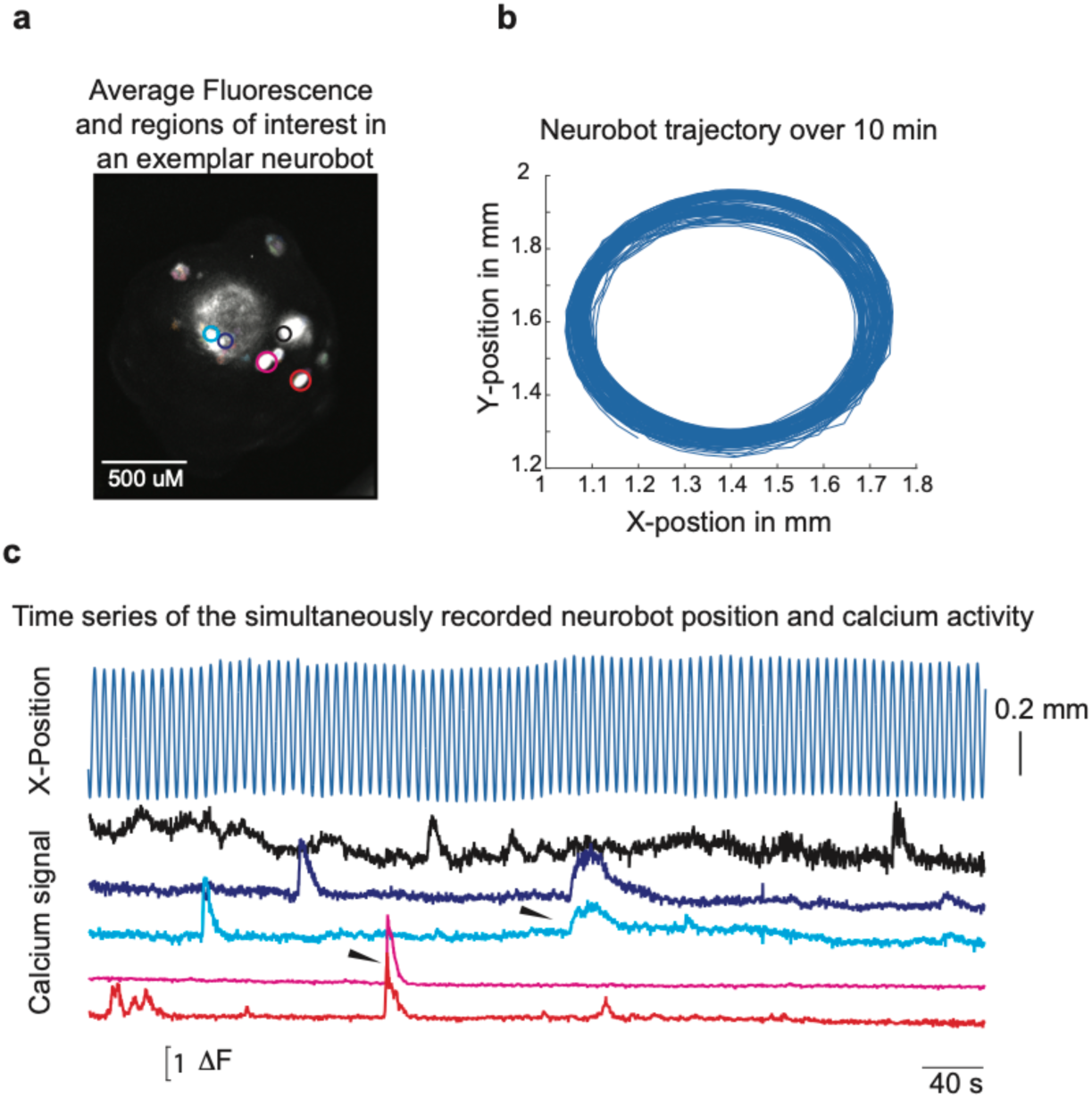
Calcium imaging in freely moving neurobots show that the implanted cells are indeed active. **a.** Average fluorescence of a freely moving neurobot *containing neurons expressing GCaMP6s* after motion correction (10 minutes of movement imaged at 5 frames per second). Colored circles correspond to regions of interest identified by suite2p software, which could be single or multiple units. **b.** Movement trajectory of the same neurobot. **c**. Top curve shows X-position of the neurobot over time. The 5 bottom curves show baseline-subtracted fluorescence activity of units labeled in panel **a**. Arrowheads point to synchronized activity in some nearby and distant ROIs.

### Neurobots tend to be more active, and show an increase in their movement complexity, compared to non-neural biobots

To assess behavioral phenotypes in neurobots and investigate potential differences with non-neuronal biobots, we video recorded spontaneous movements of neurobots and biobots in small 8-well plates (n=46 neurobots and 48 biobots, Figure 4a, Supp. Video 3: single bot moving, Supp. Video 4: bots moving in 8-well plates). We used an automatic tracking software ^28^ to measure the position of each bot at each video frame. Figure 4b shows the details of the 2D trajectories of the 8 neurobots depicted in panel a. There was a large degree of variability in these trajectories, with some bots moving in circular/oval trajectories with relatively constant diameter (Fig. 4b, bots #3, #8); bots that followed circular trajectories varying in diameter over time (Fig. 4b, bot #4); ones that made more complex, sometimes spirograph-like patterns (Fig. 4b, bots #1, #5, #6); those that were seemingly following the dish’s boundaries (Fig. 4b, bot #7); bots that circled over very small areas (Fig. 4b, bot #2); and those that did not move at all (data not shown). Interestingly, all moving bots tended to exhibit repeating behavioral motifs.

**Figure 4.**
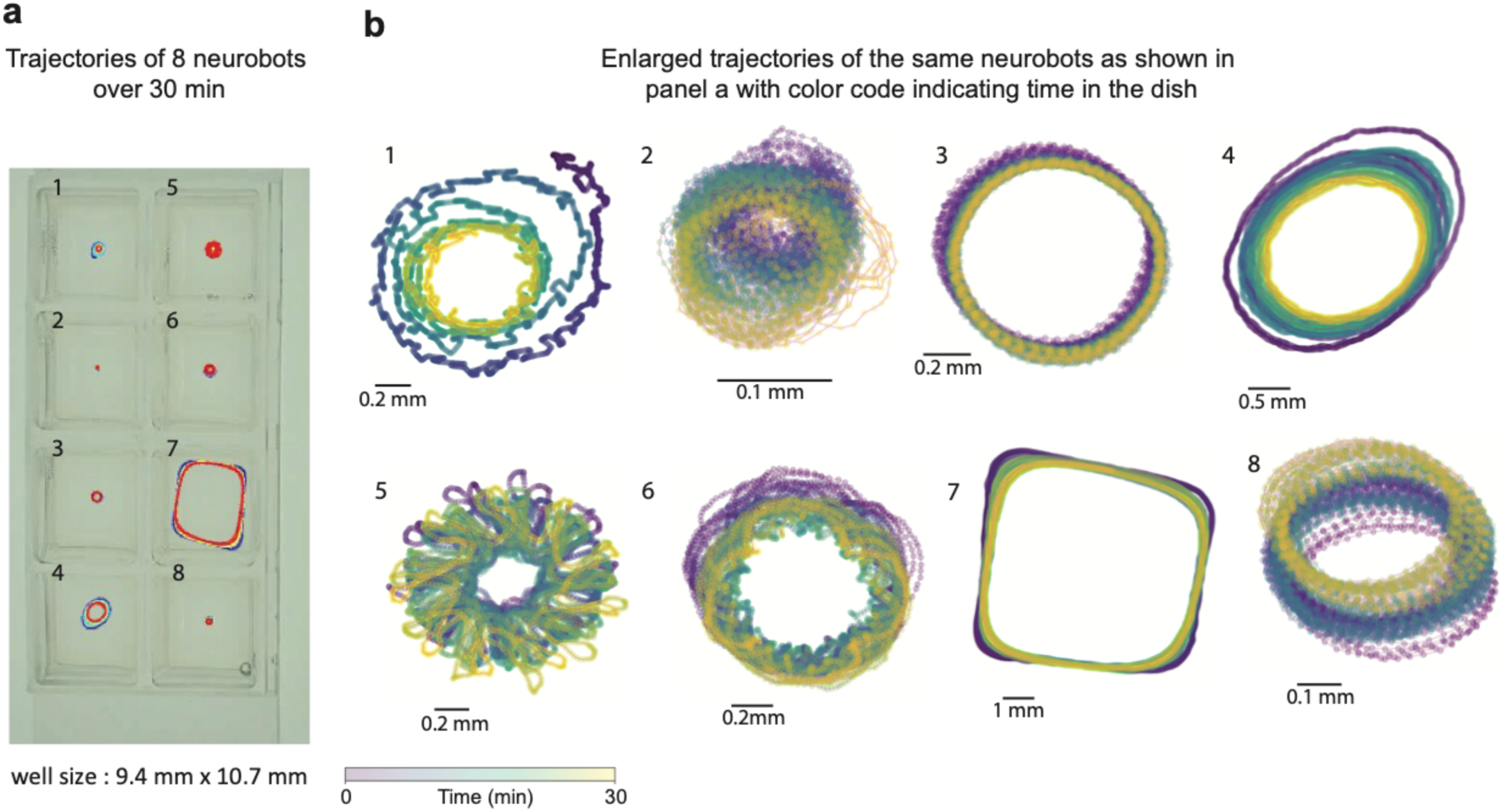
Neurobots show diverse and periodic patterns of spontaneous movement. **a**. Exemplar trajectories of neurobots moving in an 8-well plate over a 30 min trial. **b**. Details of the trajectories of the same bots as shown in panel **b**. Color gradient indicates time during the trial.

With the positional information obtained from tracking, we used custom Python code to extract 4 kinematic parameters from the bots to compare neurobots with biobots. These were total distance travelled in 30 minutes, speed, acceleration, and percentage of the well that was traversed (see Methods). We found no significant difference in the total distance traveled, the percentage of the well covered, and average speed and acceleration (Supp. Fig. 3). However, we found that the minimum movement speed of neurobots was significantly higher than that of biobots, indicating that neurobots tended to move more than biobots, remaining idle less often.

To further investigate potential differences in the complexity of movement trajectories, we used a spectral analysis, calculating the power spectral density (PSD) of the trajectory time series along the X and Y coordinates (Figure 5a-b, two exemplar trajectories and their corresponding time series on the X coordinate, see Methods). We then detected significant peaks in the X and Y PSD, summed the number of unique peaks in X and Y PSDs, and defined this number as the Complexity Index (Fig. 5c). In this analysis a complexity index of one indicates a circular trajectory with constant diameter (where both X and Y PSD have one peak at the same location), and the number increases as the trajectory becomes more complex. A complexity index of zero signifies non-moving bots.

**Figure 5.**
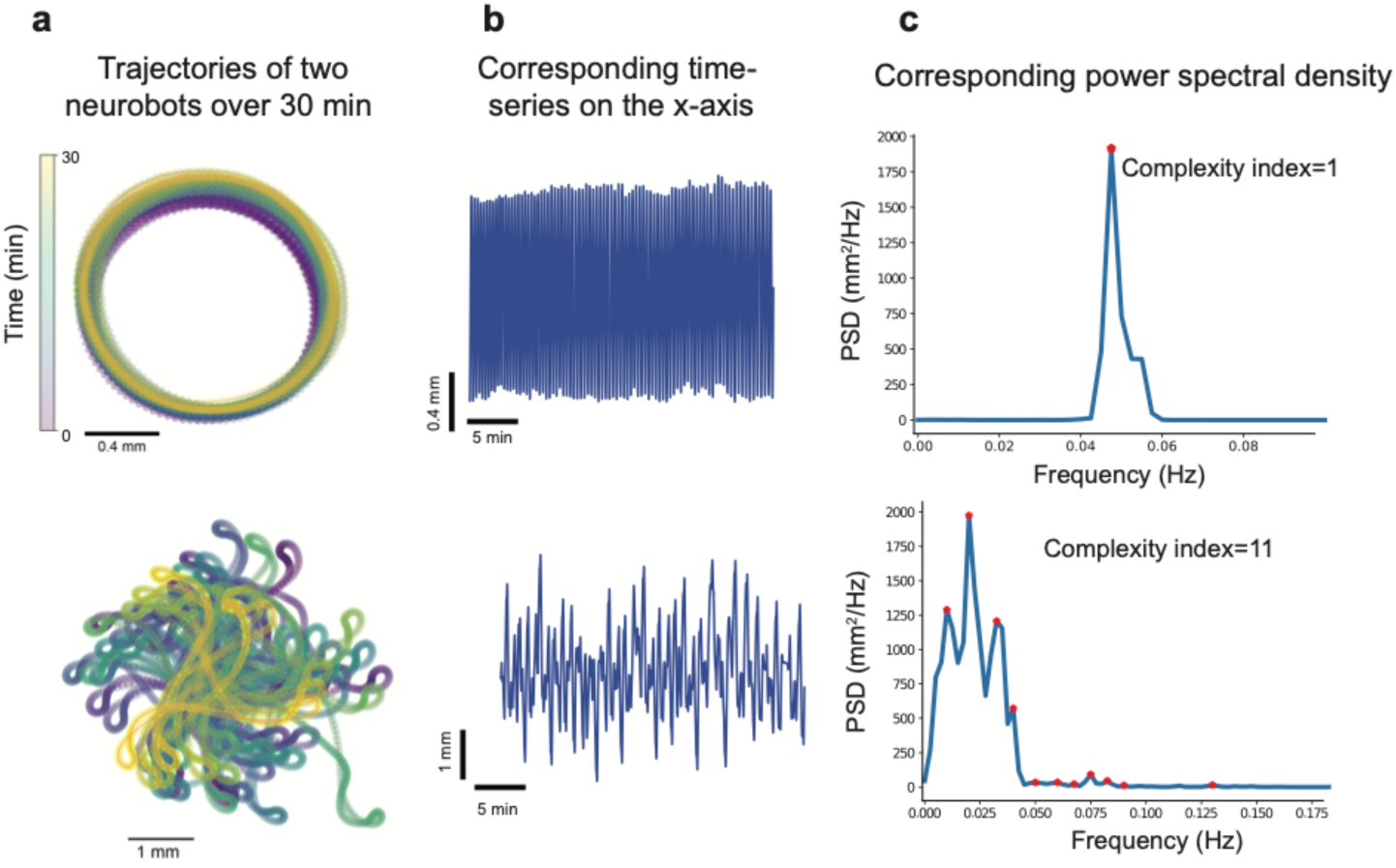
The number of peaks in the power spectral density were used to quantify the complexity of movement trajectories. **a.** Examples of simple (top panel) and complex (bottom) trajectories. **b**. Time series of the movement amplitudes projected on the X axis. **c**. Power spectral densities corresponding to time series in panel **b**. Red stars mark the location of significant peaks.

Interestingly, we found that neurobots showed a significantly higher degree of trajectory complexity compared to biobots (Fig. 6a). This increased complexity could not be explained by the roundness or size of the biobots and neurobots as these variables were not significantly correlated (Fig. 6b,c, correlation coefficient (CI, Area)=-0.03, p=0.76 and correlation coefficient (CI, RI)=-0.15, p=0.2). The increased complexity of biobots could be a result of increased variability in the beating frequency of cilia in MCCs, changes in their spatial distribution, changes in the 3D structure of the bot, or changes in the activity or distribution of other cell types (including neurons) that may modulate the ciliary beating frequency, or for other reasons. When we measured this index in sham neurobots, however, we also found an increase, albeit not significant, relative to biobots (Supp. Fig1-c), indicating that the increased complexity we observe in neurobots is at least partially due to factors other than neural signaling. Like biobots, sham neurobots were more likely to show longer periods of immobility, and their minimum speeds were not significantly different from that of biobots (Supp. Fig1-d).

**Figure 6.**
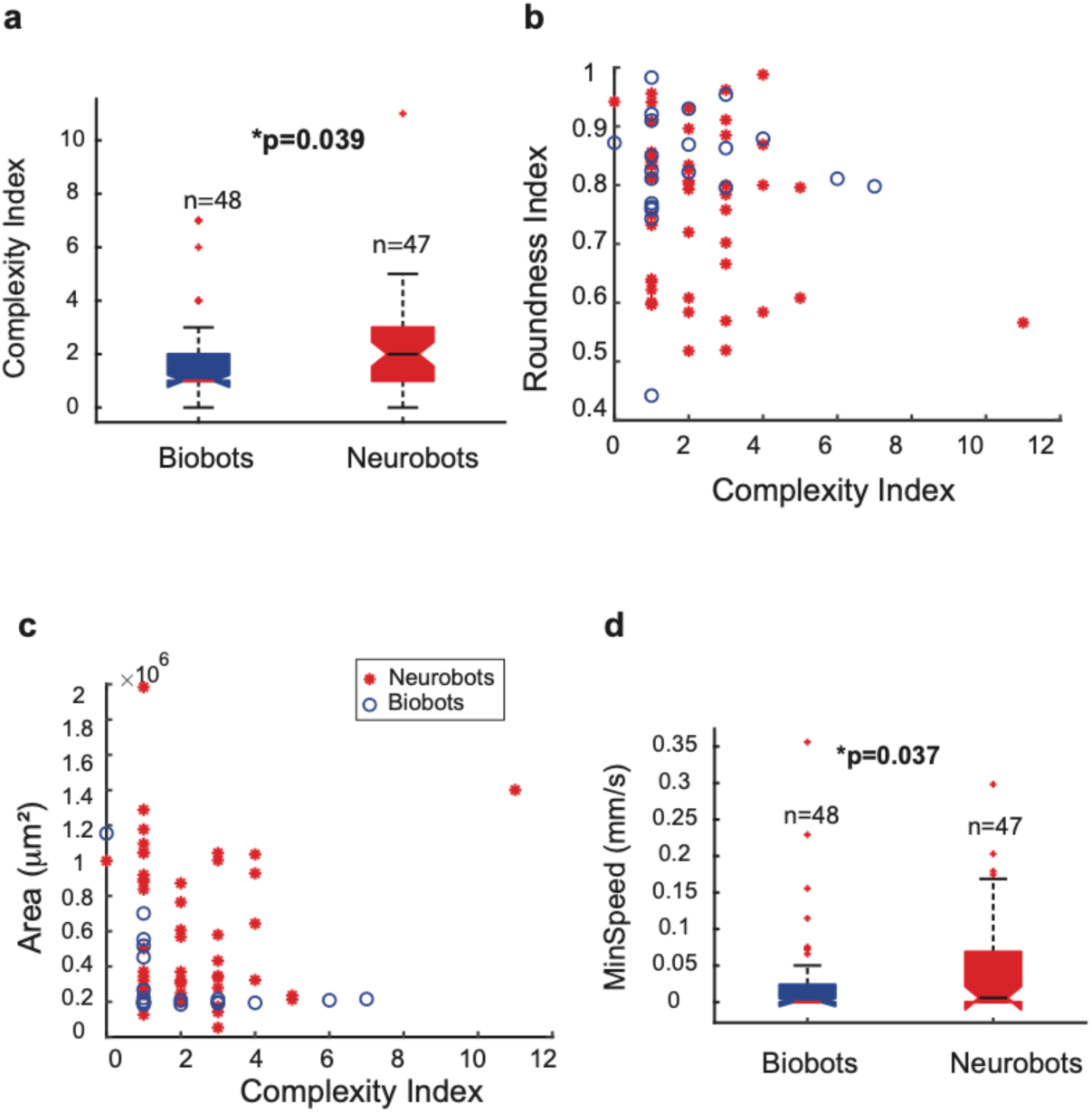
Neurobots show changes in movement patterns compared to biobots. **a.** Neurobots have more complex trajectories than biobots, and this complexity was not correlated with their size or roundness index **(b).** Complexity index was not correlated with the area of the bot **(c)**. **d**. Neurobots were more likely to be active than biobots. Non-parametric Kruskal-Wallis test was used to calculate statistical significance.

Finally, consistent with the finding that minimum speed was significantly higher in neurobots (Fig. 6d), we found that most inactive bots (Npeaks=0) were in the biobot category, with 6 out of 48 (12.5%) biobots inactive, whereas only one out of 47 (2.1%) neurobots were inactive.

### A seizure-inducing drug differentially affects behavior of neurobots and biobots

To further investigate whether neural activity could play a role in modulating trajectory complexity, we performed pharmacological experiments, treating groups of neurobots and biobots with pentylentetrazol (PTZ), a GABA_A_ receptor antagonist used for its seizure-inducing effects in animal studies^29^. Although we do not know the identity of neuronal constituents of neurobots, we reasoned that a positive result, e.g. an increase in the complexity index after PTZ treatment, could be an indirect indication of presence of GABAergic control of movement. To test this, we performed experiments in which we video recorded the movement of neurobots and biobots in regular media and compared the complexity indices after they were transferred to dishes containing 15 mM PTZ. To get an estimate on the baseline complexity index and account for potential variability due the transferring, we followed the experimental protocol depicted in Figure 7a.

**Figure 7.**
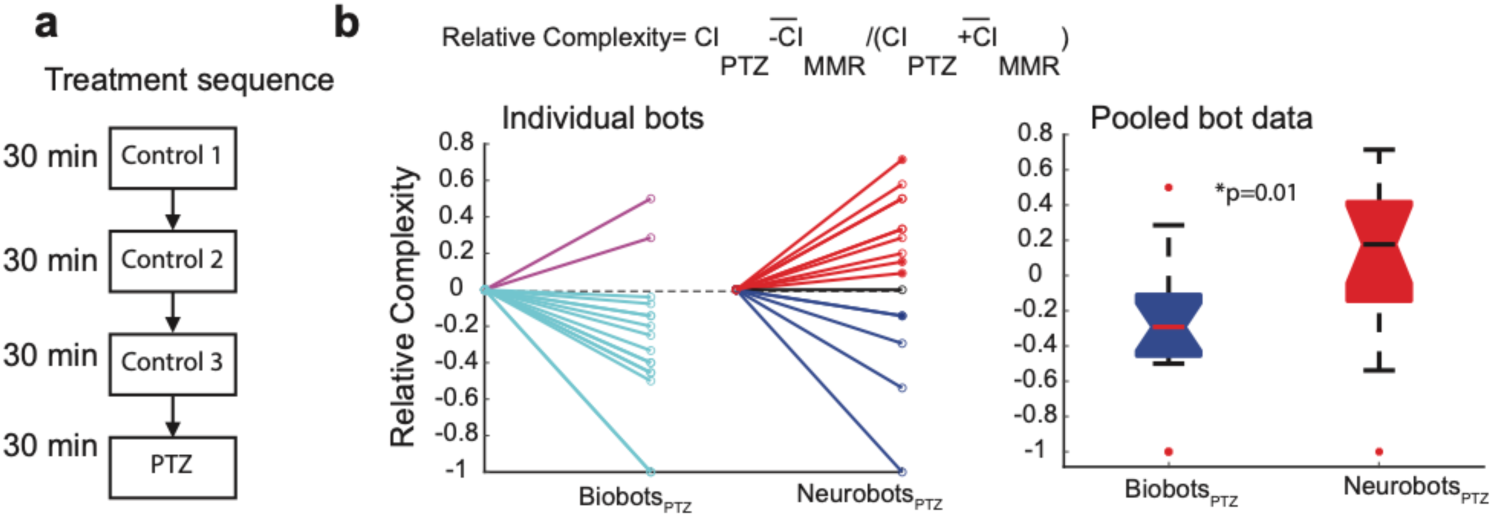
PTZ differentially impacted the movement of neurobots and biobots. **a**. Experimental protocol for testing the effect of PTZ on the movement of neurobots and biobots. Movement of bots was measured in control media, across three transfers for 30 min. The bots were then transferred to a dish containing 15 mM PTZ solution, and their behavior was measured for another 30 minutes. **b**. The effect of PTZ treatment on trajectory complexity was defined as the complexity index measured while in PTZ, relative to the average complexity measured for three consecutive MMR trials. Although most biobots reduced their complexity index relative to control, the relative complexity index for neurobots was equally likely to increase or decrease. Relative complexity of zero corresponds to the average CI for the three controls. Filled circles indicates neurobots cultured in zolmitriptan prior to testing (See Supp. Fig. 5).

Behavior of bots (16 neurobots and 16 biobots) was video recorded for 30 minutes in 8-well plates filled with regular media (control 1), after which the bots were transferred to the second and third sets of dishes containing regular media (control 2, control 3). The bots were then transferred to dishes containing PTZ (Fig. 7a). For each bot, we calculated the CIs for the three control conditions and used the average to calculate relative complexity measures for PTZ and the wash. We found that all except two biobots showed a relative decline in their movement complexity while in PTZ (Fig. 7b), thereby significantly reducing the CI relative to control (T test, p=0.009). On the other hand, the majority of neurobots showed increased complexity relative to control, although a few did show a decline (Fig. 7b). As a result of this dichotomy in the impact of PTZ, the average CI in neurobots was not significantly different relative to control (T test, p= 0.39). When we compared the relative complexity after PTZ treatment, however, we found a significant difference between neurobots and biobots, with neurobots showing significantly higher values for relative complexity (Fig. 7c). The variability of the impact of PTZ on neurobots is in fact not surprising given the potential variability in the identity and the degree of expression of neurons (Supp. Fig. 2). What is very interesting is the significant differential impact on the biobots and neurobots. Because neurobots are biobots + neurons, the fact that most of them showed an increase in their CI suggests a role for neural activity acting against the default inhibitory effect of PTZ on the movement of biobots. Alternatively, neural expression could indirectly contribute to these effects through its impact on the expression of MCCs or other cell types present on the outer surface of the bots.

### Neurobots show significant differences in the distribution of MCCs, and their roundness is anti-correlated with the degree of neural expression

To quantify the overall amount of neural expression and its relationship with neurobot morphology and behavior, we used the confocal images of neurobots immunostained with acetylated alpha tubulin antibody, which labels neurons and cilia in the MCCs. Using this data, we traced the neural processes, and quantified the position of the MCCs using Imaris software (Fig. 8a,b, see Methods). In this analysis, we did not distinguish between axons or dendrites and could not determine whether these processes belonged to individual neurons. We could, however, obtain rough estimates of the total amount of neural tissue and the degree of branching by calculating the total length of neurites and the number of terminals, which were defined as the number of nerve endings, agnostic of their identity (i.e. axonal or dendritic). Figure 8a,b shows two examples of such traces in cases of neurobots with very few and many neurites. We calculated the correlation between the degree of sprouting, expression of MCCs, bot size and shape, as well as the CI of the trajectories across all neurobots for which we had both behavioral and structural data (Fig.8c). We also calculated the correlation of all these parameters with the relative amount of neural tissue that was implanted on Day 1 as the neurobots were constructed. This ratio was calculated by dividing the area of the implanted clumps by that of the external shell (Fig. 8c: Neu/Ect, Fig. 1aii, see Methods).

**Figure 8.**
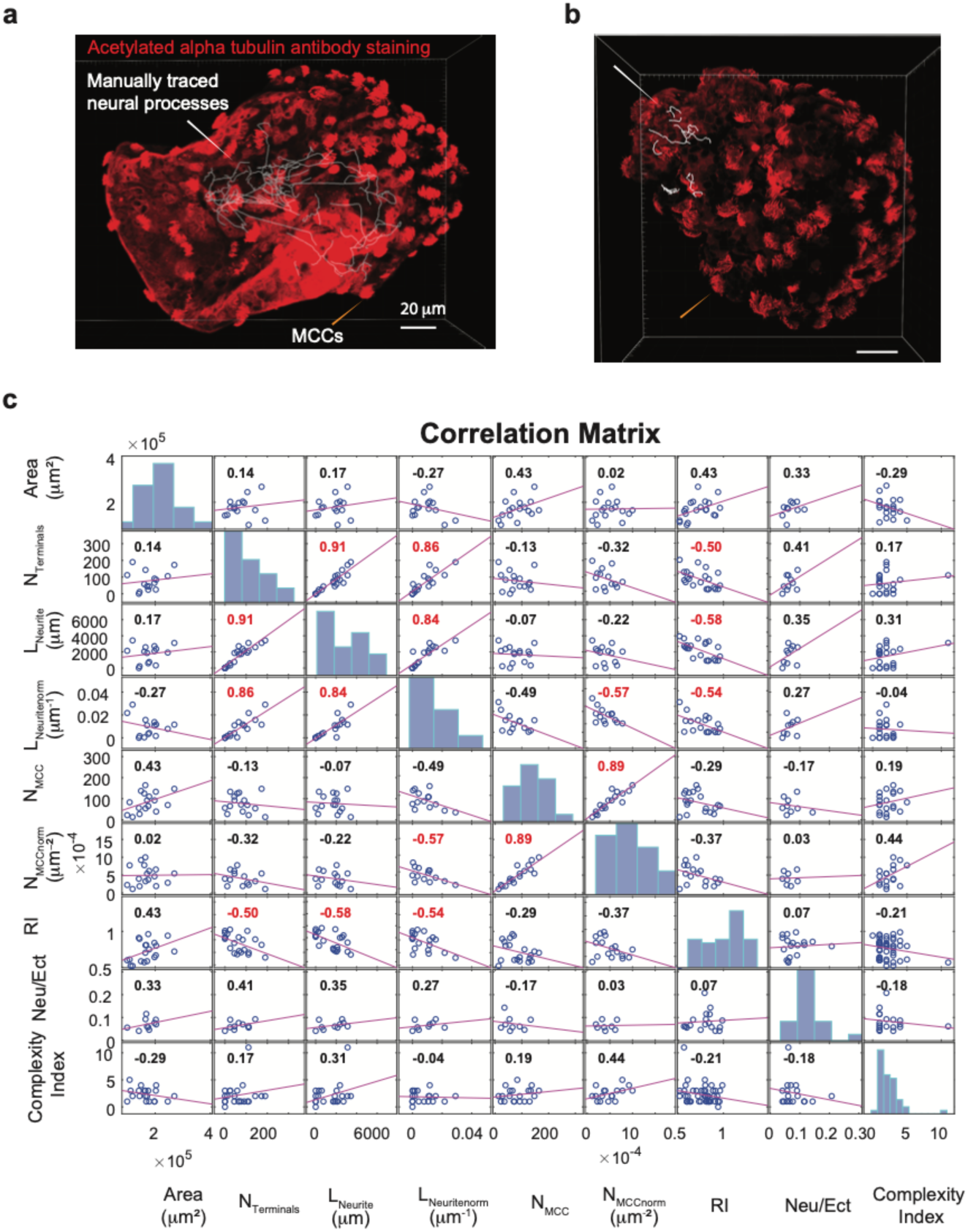
Pair-wise correlations between size, shape, neural expression, and movement complexity in neurobots. **a,b.** Examples of neurobots stained with acetylated alpha tubulin which stains multiciliated cells and neurons. Overlaid white curves show the neural processes traced using Imaris software. **a.** Example of a neurobot with small degree of innervation N_terminals_=40, L_Neurite_=753.8 mm. **b.** Example of a neurobot with a high degree of innervation N_terminals_=327, L_Neurite_=7635.9 mm. **c**. Pairwise correlation between structural parameters of all stained neurobots and Complexity Index, N_terminals_ = total number of endings, L_Neurite_=total length of neurites, L_Neuritenorm_=total length of neurites normalized to area, N_MCC_=total number of multiciliated cells on the top surface, N_MCCnorm_= N_MCC_ normalized to area, RI= Roundness Index, Neu/Ect= ratio of the areas of neural implant to ectoderm shell. White and yellow arrows point to the position of manually traced neural processes and multiciliated cells (MCCs).

Based on this analysis we found that the total number of neuron terminals was highly correlated with both absolute and area-normalized neural length (neurite density). Interestingly, we found a significant negative correlation between neurite density and MCC expression density; neurobots with higher neurite density tended to have lower overall density of MCCs. Consistent with this finding, we found that biobots (which do not have neurons) have a significantly higher density of MCCs compared to neurobots (Supp. Fig 4). Additionally, we found a significant negative correlation between the Roundness Index (RI) and neurites’ absolute length, neurites’ normalized length, and total number of terminals. That is, the more elongated the bot, the more neural expression, suggesting that the elongation could be a result of neural processes growing within the neurobot. This hypothesis is consistent with the finding that sham neurobots were not different from biobots in their roundness index (Supp. Fig. 1a). We did not however, find a significant correlation between CI and neural expression although the neurobot with highest complexity index also had the largest number of terminals and neurite length (see the outlier data point in the panels). Similarly, we found only a small correlation (non-significant) between the relative amount of implanted tissue and the degree of neural expression.

Previous studies showed that treatment with zolmitriptan, which is a selective 5-hydroxytryptamine (5-HT) 1B/1D receptor agonist, increased the degree of ectopic (but not native) neural sprouting in *Xenopus* embryos ^30^. We investigated whether this treatment would have an impact on the degree of neural sprouting in neurobots, where the neurons are all ectopic. Interestingly, we found that this treatment increased the degree of neural expression in neurobots as well, although it did not have a significant effect on any of the behavioral measurements including trajectory complexity (Supp. Fig. 5). In this group we found a tight correlation between the ratio of neural implant to ectoderm shell and total number of terminals, as well as total neural length (Supp. Fig. 5). These results indicate that neurons in a neurobot behave as ectopic, not native, cells.

Notably, 4 of the 16 neurobots in the PTZ study presented in the previous section were cultured in zolmitriptan (filled circles Fig. 7b, Supp. Fig. 5), three of which showed an increase in relative complexity. This result points to the possibility that this treatment may bias neural expression towards those that respond to PTZ i.e. GABAergic neurons. Further experiments are required to characterize the impact of zolmitriptan treatment on the neural expression patterns within neurobots.

In summary, neural growth in neurobots significantly impacts their shape and the distribution of MCCs, and this growth could be potentially increased by modulating serotonergic signaling. The degree to which neural expression contributes to trajectory complexity remains elusive. The lack of observed correlation between neurite growth and complexity index points to the potential heterogeneities in cell type expression and connectivity profiles of neurites across neurobots.

### The three types of bots exhibit significantly different patterns of gene expression

Like CNS structure and function, transcriptomes are usually thought of as being shaped by a long history of selection. In standard organisms, they are also shaped by neural inputs ^31,32^. What would the transcriptome of a novel construct with a nervous system look like? With this question in mind, we next asked what changes to default biobot transcriptomes, if any, would be induced by the presence of neural tissue. To characterize the transcriptome of neurobots and compare it to its non-neuronal counterparts (i.e. biobots and sham neurobots), we performed bulk RNA sequencing of their tissue. For each bot type four biological samples were included (neurobots: NB1-4, biobots: BB1-4, sham neurobots: SH1-4). Due to the small size of the bots, and therefore, small amount of RNA, each sample comprised tissue from multiple bots (see Methods).

We found a high degree of correlation between normalized gene expression levels among all samples within each group (Fragments Per Kilobase of transcript sequence per Millions base pairs sequenced, FPKM^33^), indicating reliability and repeatability of the results (Fig. 9a). Moreover, we found that gene expression levels in biobots and sham neurobots were much more correlated to one another than to neurobots (Fig. 9a, see Methods). Similarly, neurobots could clearly be separated from shams and biobots based on the principal component analysis on the normalized gene expression value (FPKM) of all samples (Fig. 9b, see Methods).

**Figure 9.**
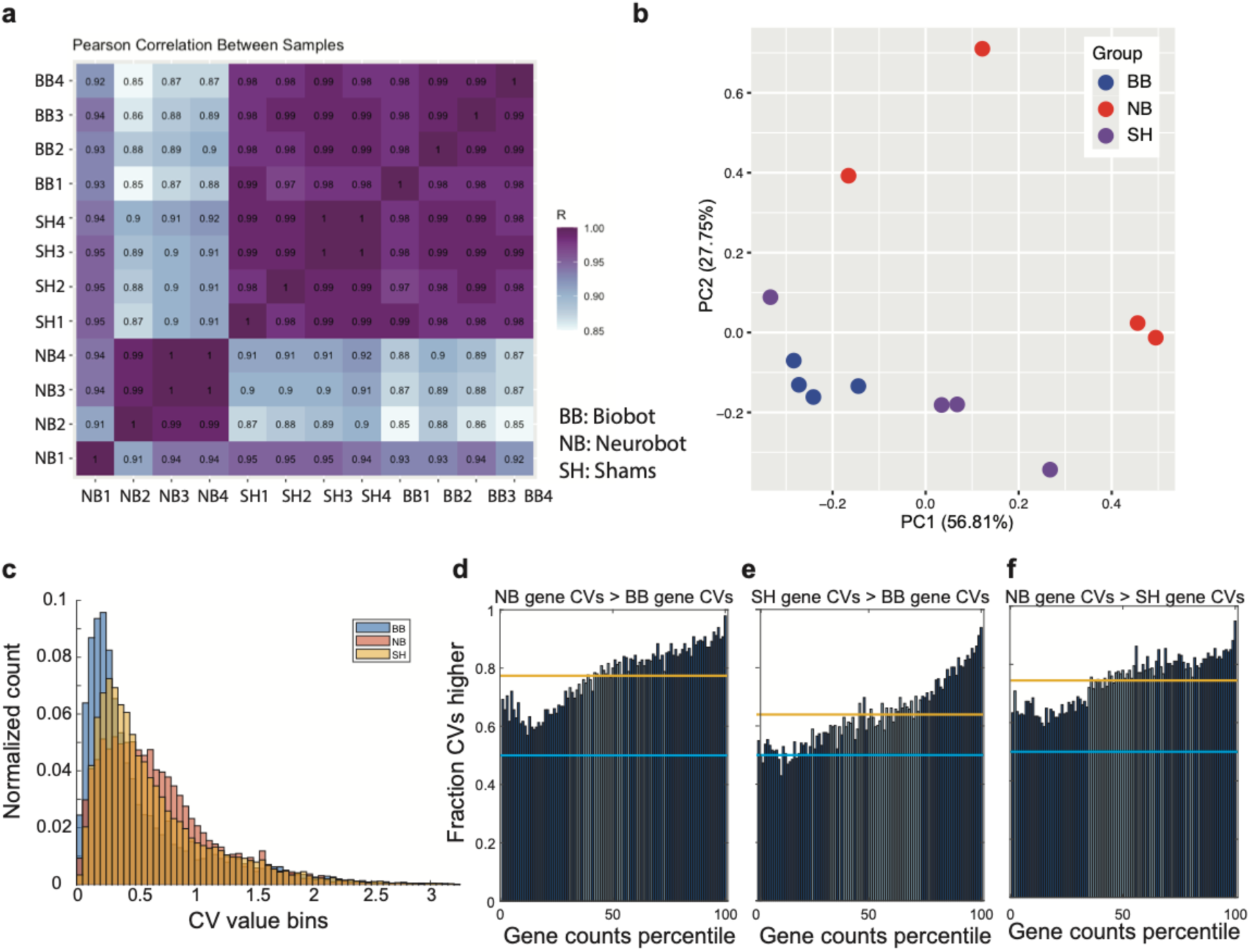
Comparison of gene expression and its variability between neurobots, biobots, and sham neurobots. **a.** Pearson correlation of gene expression within and across groups of neurobots (NB), sham neurobots (SH), and biobots (BB). The closer the value to 1, the more similar the expression patterns. **b.** Principal component analysis of gene expression values. Each dot corresponds to one sample, which contained multiple bots of one kind. **c.** Histograms of coefficients of variation (CV) in gene counts (FPKM) in neurobots (red), biobots (blue), and sham neurobots (yellow). Neurobots showed a significantly higher variability in their gene counts compared to both biobots and shams, and shams showed a higher variability compared to biobots. Genes with higher counts showed higher degree of variability in their expression when comparing neurobots with biobots and shams (**d,e**). Dark blue bars mark the bins where the difference in CV was significantly different from what is expected if the ranking of genes were randomly shuffled (see Methods). Similarly, genes with low levels of expression showed lower coefficient of variation than expected by chance. **f.** Same as **d** and **e** but comparing neurobots with shams. Bin values above 0.5 (light blue line) indicate that more than half CVs in that bin had higher values in that comparison. Orange horizontal line indicates average of all fractions regardless of the ordering. Significance was assessed relative to this value using Kruskal Wallis non-parametric statistics (see Methods).

We next compared the gene count variability across the samples of biobots, neurobots, and sham neurobots on a gene-by-gene basis (Fig. 9c-f). We found that neurobots showed a significantly higher variability, quantified by coefficient of variation (CV, see Methods) in their normalized gene counts (FPKM) across neurobot samples, compared to both biobots and shams, and samples of sham neurobots showed a higher variability compared to samples of biobots (Fig. 9c). We then compared pairs of groups (e.g., NB and BB) to determine the fraction of genes that had a greater CV in normalized gene count in one group than the same gene in the other group. For the chosen pair of groups, genes were ranked by the mean count value across all pools of both groups, and the CV of each gene’s counts across the pools of each group was calculated (Supp. Fig. 9, see Methods). The CV list was split into 100 bins (percentiles) containing equal number of genes, and the fraction of genes in the bin for which the CV of the first group was greater than that of the second group was found and plotted.

We found that in all bins, more than half of the neurobot genes showed higher CV compared to biobot and sham group (Fig. 9d,f, blue line is at 0.5). For the sham neurobot group, genes in most, but not all bins showed higher CV compared to biobots (Fig. 9c). In addition, we found a trend for genes with higher normalized counts showing a higher degree of variability in their expression when comparing neurobots with biobots and shams (Fig. 9d,f). This pattern was significantly different from what is expected from chance in most bins (dark blue bins), i.e. relative to the average CV calculated across all bins regardless of the order (orange line, see Methods). The overall higher variability seen in neurobots and shams compared to biobots could be due to several factors. First, in both neurobots and shams, the implanted cells are harvested from ∼50 embryos, whereas a biobot is made out of a single embryo. Moreover, higher variability in the implanted bots is expected due to the high variability in the size of the implants in both neurobots and shams. However, these are likely not the only factors involved, as neurobots showed significant differences in their gene count variability compared to shams. Neural differentiation, therefore, likely plays an important role in the increased variability in gene counts seen in neurobots.

Additionally, we found that neurobots included a significantly larger number of genes that were differentially expressed relative to biobots and sham neurobots (Fig. 10 a,b), whereas biobots exhibited a smaller subset of differentially expressed genes relative to sham neurobots (Fig. 10c). Moreover, the number of significantly upregulated genes (p<0.05, red dots with positive fold change) in neurobots compared to biobots and shams (6774 and 6859 genes, respectively), were much higher than those that were significantly downregulated (red dots with negative log fold change, 3578 in neurobots vs biobots and 4010 in neurobots vs shams), resulting in highly asymmetric volcano plots (Fig. 10 a,b). This was not the case when comparing shams with biobots (Fig. 10c, 1733 upregulated and 1429 down regulated genes). These results are consistent with a gain of function as a result of neural growth in neurobots.

**Figure 10.**
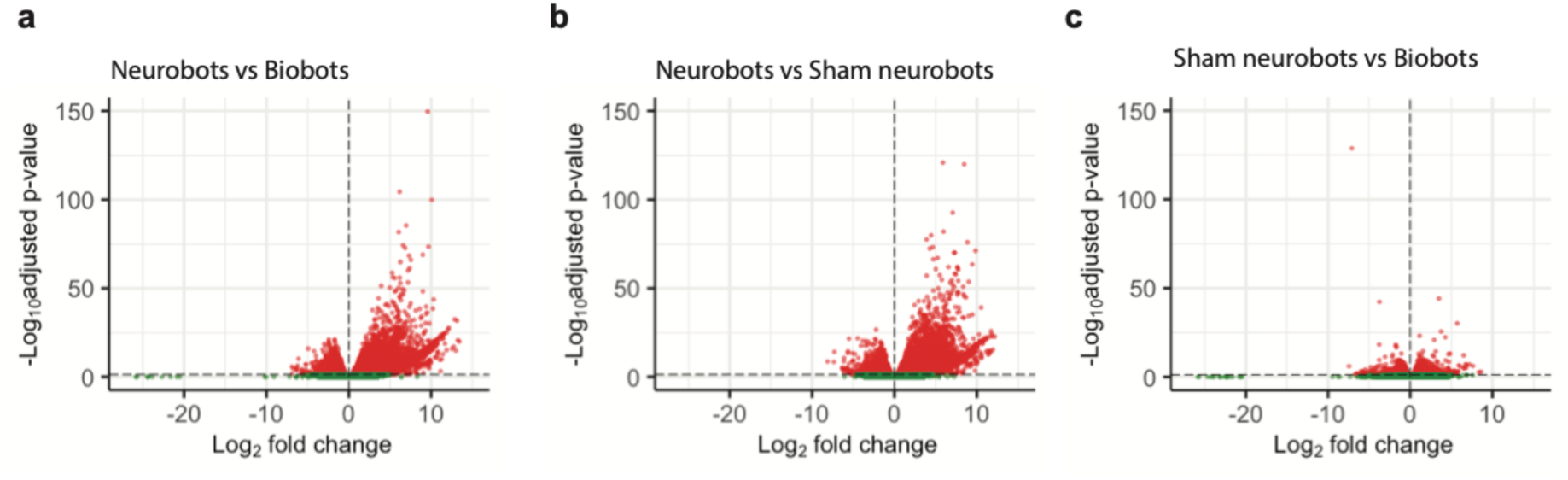
Distribution of differentially expressed genes between different bot groups. (**a-c).** X-axis shows the fold change in gene expression between samples of different groups and Y axis shows the statistical significance of the difference. Red dots represent genes that were significantly up (positive values on the X-axis) or down regulated (negative values on the X-axis); green dots represent genes with no significant change.

We next investigated which biological functions or pathways are significantly associated with the differentially expressed genes. We used Gene Ontology (GO) enrichment analysis to annotate genes to biological processes (bp), molecular function (mf) and cellular components (cc). Due to the large number of upregulated genes in neurobots, we focused on the highly overexpressed genes (4 log-fold or more increase in expression, 2445 genes when comparing neurobots to biobots and 2026 genes when comparing neurobots to shams) for this analysis. The most significantly upregulated pathways in neurobots relative to biobots, as well as in neurobots relative to shams, related to nervous system development, and synapse and neuron projection (Fig. 11 a,b). One of the interesting genes we found up-regulated specifically in neurobots relative to biobots and shams is *Dact-4*, a member of an evolutionarily conserved family of Dishevelled-binding proteins involved in the regulation of Wnt and TGF-beta signaling which is expressed in the Spemann organizer^34^. This suggests that the presence of neurons might exert an organizational influence on the surrounding soma; this molecular signature is consistent with known roles of the nervous system to direct cell behavior in cancer suppression^35–37^, regeneration^38^, and embryonic morphogenesis^32,39^. Trans-synaptic signaling and neurotransmitter receptor activity were also significantly upregulated in neurobots. This included glutamatergic, GABAergic, cholinergic, dopaminergic, serotonergic, and glycinergic receptors. Surprisingly, exclusively in neurobots, we also found significant enrichment in pathways involved in visual perception (Fig. 11a).

**Figure 11.**
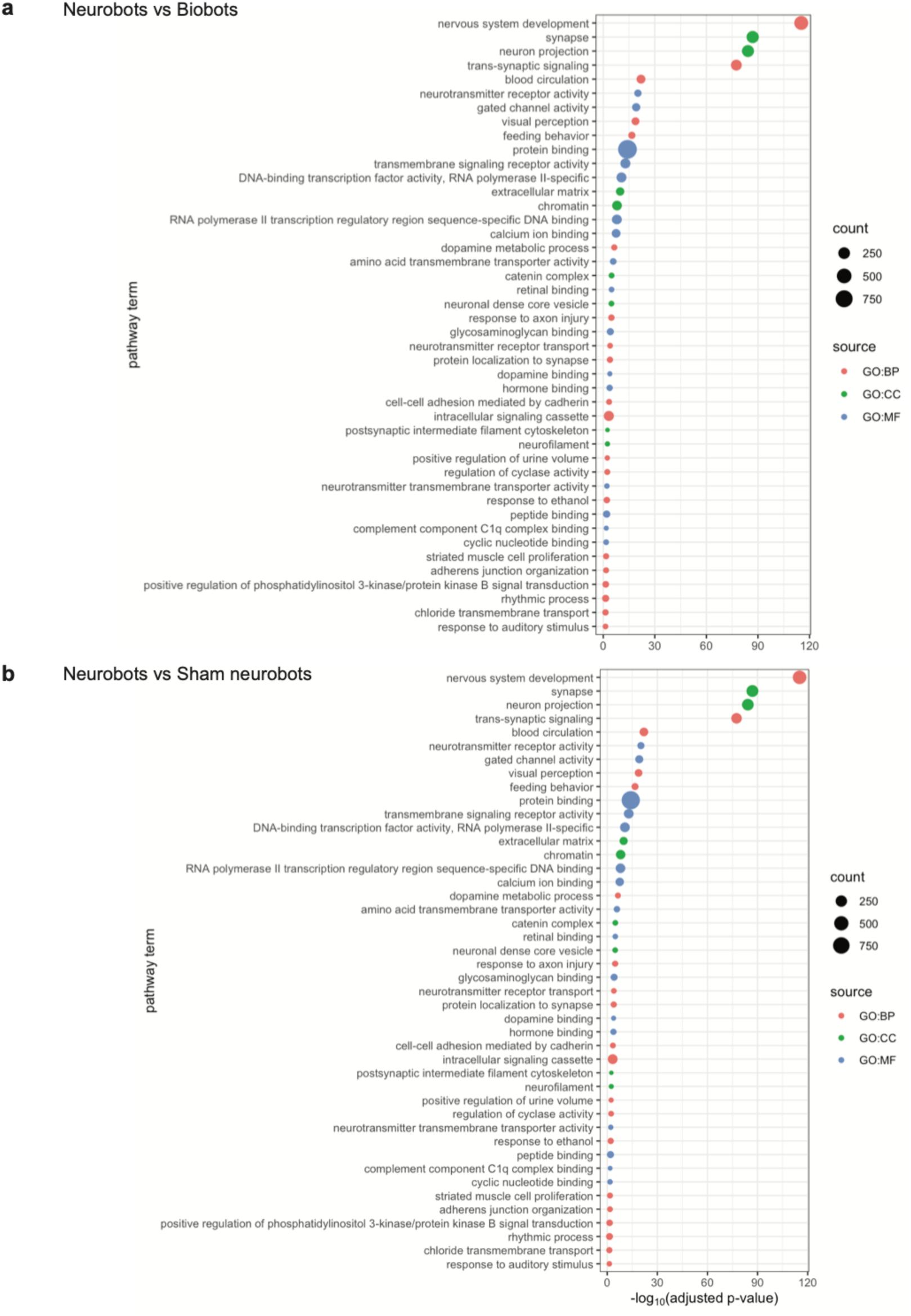
Enrichment analysis performed using Gene Ontology annotations on differentially expressed genes with at least 4-fold upregulation in expression. **a.** neurobots vs. biobots **b.** neurobots vs. shams, BP:Biological processes, CC: Cellular Components, MF: Molecular Function.

Although pathways relating to neuron projection and trans-synaptic signaling were slightly upregulated in sham neurobots compared to biobots, there were far fewer genes in each pathway, and they were less significant in their degree of upregulation compared to those in neurobots (Supp Fig. 6a). Further, there were relatively fewer genes downregulated when comparing neurobots with biobots, and shams (63 and 116 genes respectively) and very few genes were significantly downregulated by 4-log folds when comparing shams with biobots (38 genes). Our enrichment analysis showed that the largest group of downregulated genes in neurobots compared to biobots belonged to the cellular component pathway (extracellular region in Supp. Fig. 6b). Interestingly, this pathway includes some of the genes expressed in the *Xenopus* skin, including glycoprotein 2 (*gp2*), and mucin (*muc17*) suggesting that the properties of the ‘skin’ of neurobots might be different from those of biobots.

In order to identify functional biological modules of differentially expressed genes, we extracted the largest protein-protein-interaction (PPI) sub-network of these genes using the STRING database. We then performed network embedding and clustering using multi-nonnegative matrix factorization (MNMF)^40^ to find specific functional biological modules. The clusters were subsequently enriched using g:Profiler^41^. Based on this analysis, we identified 25 clusters for neurobots vs biobots comparison and 5 clusters in neurobots vs sham neurobots comparison (Supp. Spreadsheet, Supp. Fig 7 and 8). There were not enough upregulated genes between biobots and sham neurobots to allow for this analysis. Similarly, due to the low number of downregulated genes, the network analysis could not be performed at either 4- or 2-fold change threshold level.

Consistent with the findings from the enrichment analysis, we found clusters containing genes critical for the development of the nervous system, cell fate commitment, and *wnt* signaling pathways (Cluster 3, Supp. Fig. 7a, see the NB vs BB tab in Supp. Spreadsheet). A plethora of growth factors (e.g. various FGFs, BDNF and EGF), and their receptors were revealed by Cluster 17, which also showed enrichment in enzyme-linked receptor protein signaling pathways (Supp. Fig. 7b, see the NB vs BB tab in Supp. Spreadsheet).

Neurobots contained genes encoding various neurotransmitter receptors including glutamate (e.g. *gria1-4*), kainate receptors (e.g. *grik1,2,3,5*), GABAergic (e.g. *gabara3,5; gabarb3*) and glycinergic receptors (e.g. *glra3; glrb*), genes encoding voltage gated calcium channels (*cacng3,4,5,7,8*), as well as those involved in the uptake of neurotransmitters (e.g. *slc1a1,2,3*). Genes with important roles in synaptic plasticity were also present in neurobots (*arc, camk2b*, Cluster 15, Figure 12a, see the NB vs BB tab in Supp. Spreadsheet)^31^. Cholinergic and muscarinic (*chrna2,3,4,5,7; chrnb2,3,4, chrm2,4,5*), serotonergic (*htr1a,b,e*; *htr2a*), and dopaminergic (*drd1,2,4*) were among other neurotransmitter receptors (Cluster 23, Figure 12b, see the NB vs BB tab in Supp. Spreadsheet)

**Figure 12.**
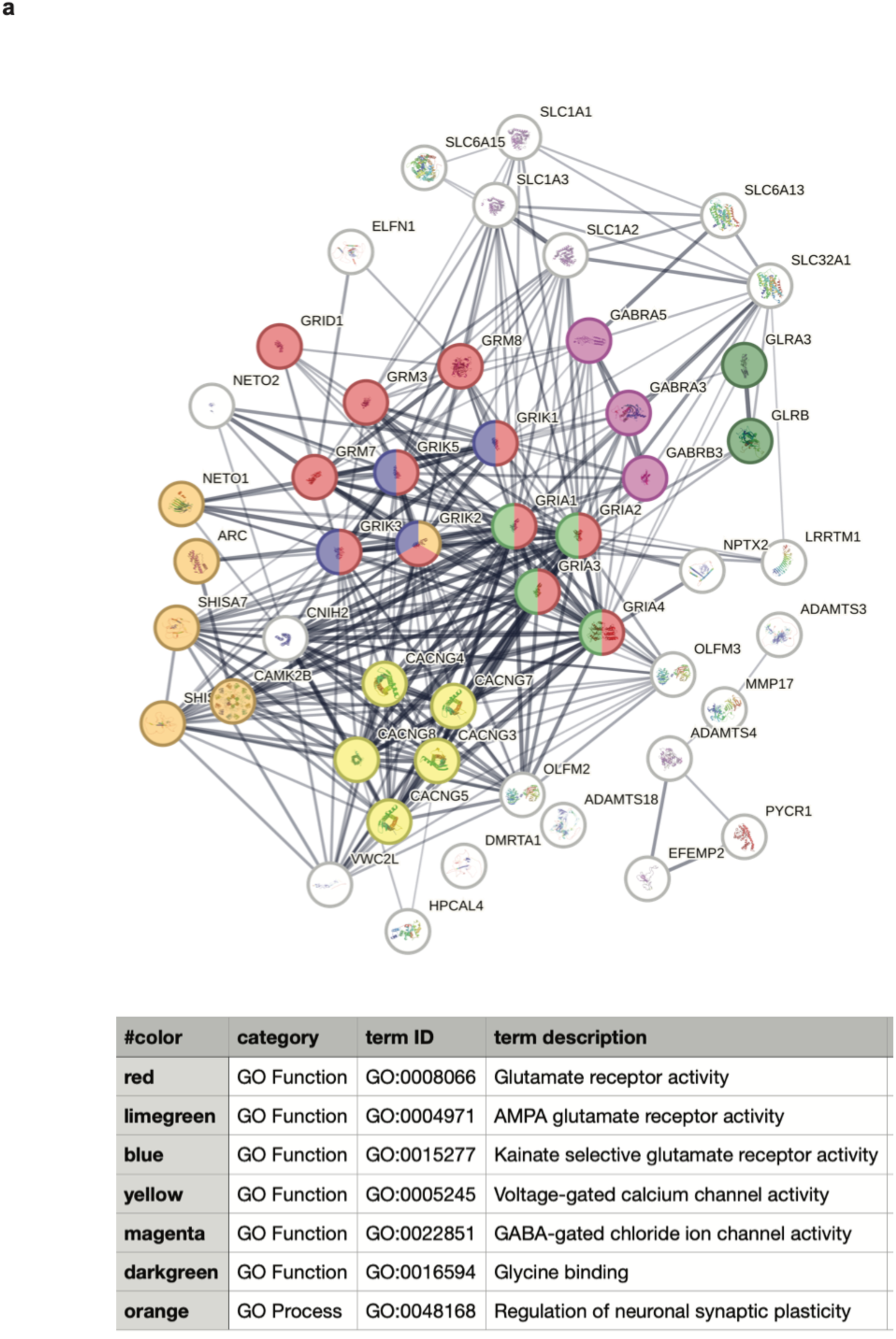
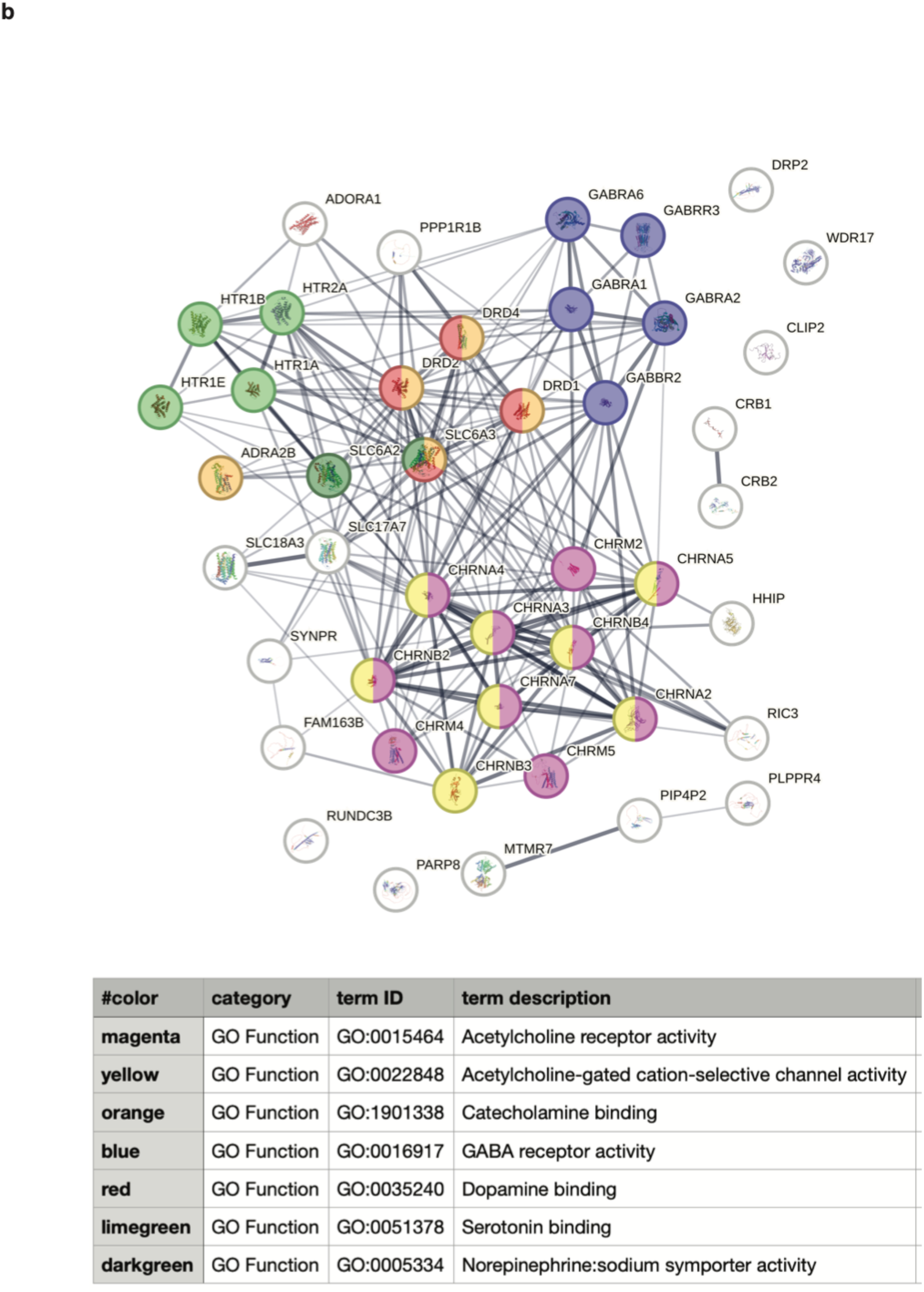
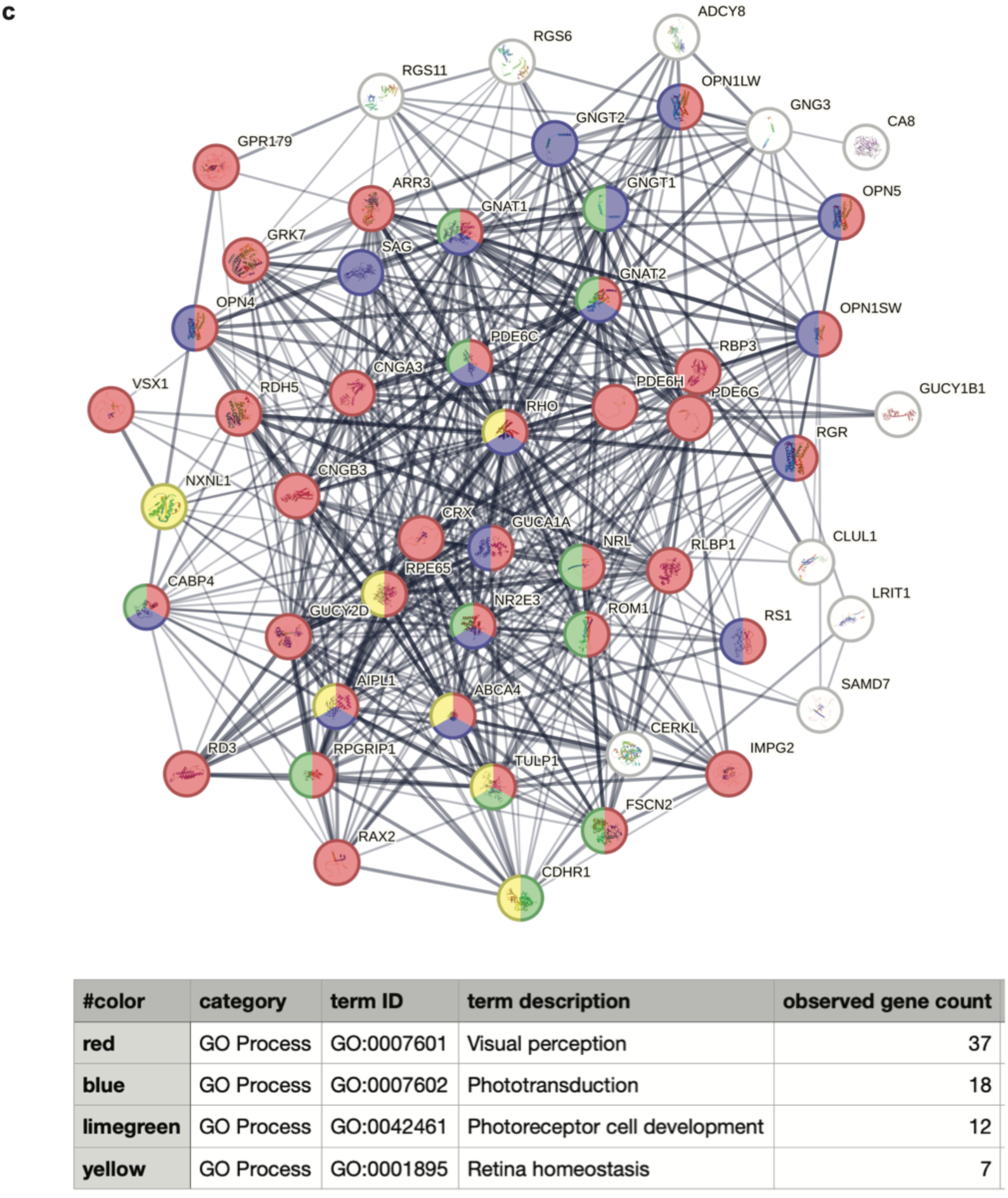
Cluster-based network connectivity patterns among genes that were upregulated in neurobots compared to biobots. **a.** Cluster 15 **b.** Cluster 23 **c.** Cluster 1. Network connectivity was calculated using STRING online tool. The edges indicate both functional and physical protein associations the line thickness indicates strength of data support. Only nodes with interaction scores with confidence higher than 0.4 are shown. Nodes of special interest are highlighted in color.

Notably, one of the largest clusters (Cluster 1) contained genes encoding various aspects of visual perception, phototransduction, and photoreceptor development (Fig 12 c, see the NB vs BB tab in in Supp. Spreadsheet). Specifically, this cluster included red and violet cone opsins (*opn1lw, opn1sw*), retinal G-protein couple receptors (*rgr*), melanopsin (*opn4, opn5*), rhodopsin (*rho*), as well as many other related genes that encode proteins involved in visual processing. In addition to Cluster 1, Cluster 12 (Supp. Fig. 7c) also included genes related to eye, lens, and retina development, including genes found in major retinal cell types^42^, i.e. retinal ganglion cells (*neurod1,2; pou4f1*), and horizontal cells (*onecut1, lhx1*). Genes found in bipolar cells (*unc5d*), and amacrine cells (*prdm13*) were also present in Clusters 21 and 22, respectively (Supp. Fig. 7d,e, see the NB vs BB tab in Supp. Spreadsheet), suggesting that neurobots could potentially sense and process light stimuli.

Many other clusters included significant enrichment in genes encoding various aspects of the nervous system. This included Cluster 5, which contained various synapsins, tubulins, and microtubule associated proteins, which are implicated in biological processes such as synaptic vesicle cycle, neuron development, regulation of neurotransmitter secretion, and synaptic vesicle localization (Supp. Fig. 7f, see the NB vs BB tab in Supp. Spreadsheet). Cluster 9 revealed the presence of voltage gated ion channels including various types of sodium and potassium channels as well as voltage gated calcium channels (Supp. Fig. 7g, see the NB vs BB tab in Supp. Spreadsheet). Cluster 11 contained genes encoding various G-protein coupled receptors, as well as those implicated in the modulation of chemical synaptic transmission (Supp. Fig. 7h, see the NB vs BB tab in Supp. Spreadsheet). Cluster 14 revealed the presence of various hormones and neuropeptides and their receptors (Supp. Fig. 7i, see the NB vs BB tab in Supp. Spreadsheet). Cluster 21 included genes important for axonogenesis and neuron projection development (Supp. Fig. 7d, see the NB vs BB tab in Supp. Spreadsheet).

Moreover, we found that there was significant upregulation in genes encoding extracellular matrix constituents (Fig. 11a, Cluster 20, Supp. Fig. 7j, see the NB vs BB tab in Supp. Spreadsheet) including collagen (*col17a, col4a2*), which is the most abundant fibrous protein and constitutes the main structural element of extracellular matrix^43^, and fibrillins, which are glycoproteins that are secreted in the extracellular matrix and provide mechanical support in connective tissue (*fbln1*)^44^. This finding provides support for the hypothesis that the internal cavity of neurobots (Fig. 2, Supp. Fig. 2) may not be empty, but be comprised of ECM materials, providing structural support for neural processes.

As expected, network embedding and clustering analysis of the upregulated genes in neurobots relative to shams similarly revealed overexpression of genes relating to synapse organization, regulation of neurotransmitter and receptor activity, chemical synaptic transmission (within Clusters 4,5), visual perception (within Cluster 3), neuron projection, and perineuronal nets (within Cluster 2, Supp. Fig. 8). These findings shed light on biological pathways/molecular functions that are innately present in neurobots in the absence of any external manipulations and will inform future work toward building specialized neurobots through selective enhancement of these pathways.

Finally, we tested the hypothesis that neurobots are expressing a more ancient transcriptome as a result of their nascent evolutionary history. We applied a phylostratigraphic analysis for the differentially expressed genes in the different conditions (Fig. 13, NB vs SH and NB vs BB). Interestingly, we found that more than 54% of upregulated genes in neurobots fall into the two categories of most ancient genes (“All living organisms” and “Eukaryota”, Fig. 13a). By comparison, very few ancient genes are downregulated. In total 279 are downregulated in these two strata for the NB vs BB conditions, and 233 for the NB vs SH condition (Fig. 13b), while for the upregulated genes we obtained 941 and 1109, respectively. Therefore, we conclude that the development of neurobots involves a transcriptomic shift towards very ancient genes for neurobots compared to biobots and shams.

**Figure 13.**
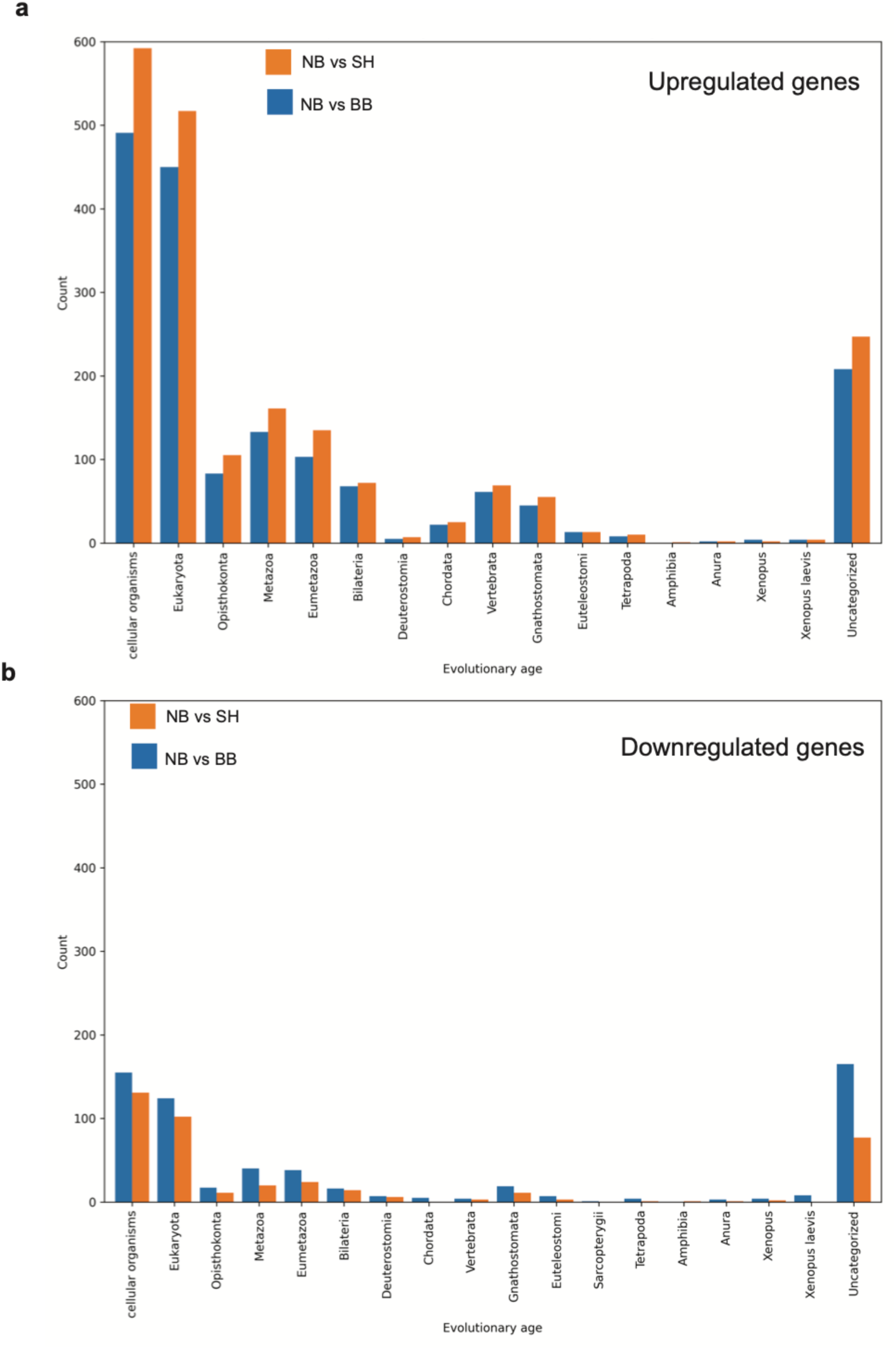
Phylostratigraphic analysis of upregulated or downregulated transcripts in neurobots compared to biobots and shams. **a.** 54% of upregulated genes in neurobots fall into the two categories of most ancient genes (“All living organisms” and “Eukaryota”). **b.** Very few ancient genes are downregulated. In total 279 are downregulated in these two strata for the NB vs BB conditions and 233 for the NB vs SH condition while for the upregulated genes we obtained 941 and 1109, respectively.

## Discussion

In this study we built and investigated behavioral, anatomical, and transcriptional properties of novel living constructs with incorporated neural tissue. Using *Xenopus laevis* embryonic cells, we built two types of living constructs: one using ectodermal cells (biobots) as reported in prior studies^17,45,46^, and another, novel construct, made using ectodermal and neural precursor cells (neurobots, Fig. 1a). We showed that neurobots are viable and self-motile like their non-neuronal counterparts (Supp. Vid. 3,4), and that the implanted neural precursor cells indeed differentiate into neurons and extend their processes throughout the construct (Fig. 1-2, Supp. Fig. 2).

We found that neurobots became significantly larger than biobots and were significantly more elongated (Fig. 1b,c). We saw a large degree of variation in the shape and innervation pattern of neurobots, but in all neurobots we found a central ‘cavity’ seemingly devoid of any cell bodies (Fig. 2, Supp. Fig. 2). We speculate that this region is not truly empty and may be filled with extracellular matrix-like structures (ECM). The presence of neurites extending in extremely straight courses points to the presence of such supporting structure (Fig. 2b, red arrowhead). Consistently, our transcriptomics data showed significant upregulation in the expression of genes related to ECM in neurobots (Fig. 11a,b, Fig. 13). This finding points to tight coupling between neural network formation and ECM gene expression, even in the context of completely novel embodiments.

Neurobots exhibited a diverse range of movement patterns, and these patterns were more complex than those observed in their non-neuronal counterparts, indicating that neural expression may affect movement either directly, i.e. through neural signaling to the motor effectors, or via changes in the expression patterns of motor effectors. We did observe a negative correlation between the degree of neural expression and the density of multiciliated cells (Fig. 8C).

Simultaneous recording of neural activity and behavior could be used to assess the potential neural correlates of the behavior. Our calcium imaging experiments indicated the presence of neural activity. However, measuring this activity in freely moving neurobots was complicated by their movement, which often resulted in losing track of specific neurons, especially in neurobots that showed extensive rotational movement in 3D (Supp. Vid. 1). Such 3D movements could be suppressed by making flattened neurobots, however, these neurobots tended not to move as much (data not shown). The role of neural activity in trajectory complexity may be alternatively investigated in the future through silencing/activating neurons, either pharmacologically or by using optogenetics, and measuring the impact on trajectory complexity. Consistent with a role of neural activity in trajectory complexity, we found that treatment of biobots and neurobots with the GABA_A_ receptor antagonist PTZ resulted in significantly different outcomes. Although most biobots decreased their movement complexity with PTZ treatment, this was not the case for neurobots (Fig. 7). In fact, the majority of neurobots showed an increase in movement complexity, suggesting that neural activity may contribute to the observed differential effect. Further experiments are required to assess neural control of spontaneous and evoked behaviors.

Anatomically, we found that although the majority of neural processes emanating from the implanted neural precursor ‘clumps’ sprouted within the central cavity of the bot, some processes did extend toward the outer epithelium (arrows heads Fig. 2, Supp. Fig. 2). Although we have not demonstrated the presence of synaptic connectivity with the surface epithelial cells, these neural processes are in a position to allow them to modulate activity of the surface epithelial cells, including multiciliated cells, goblet cells, and serotonergic cells among others. It is known that serotonergic signaling impacts the beating frequency of cilia on multiciliated cells, and serotonergic receptors are present on these cells^20^. Serotonergic neurons sending processes to the surface could potentially modulate the movement dynamics of the neurobots. GABAergic receptors are found on the surface of the mucus secreting goblet cells, and treating *Xenopus* embryos with bicuculline, which is also a GABA_A_ antagonist, was shown to inhibits mucus secretion^47^. Our results from the PTZ experiments, which showed a decrease in movement of biobots, could be a result of changes in mucus secretion, which might in turn impact the beating frequency of multiciliated cells.

We raised a group of neurobots in zolmitriptan, which is a selective 5-hydroxytryptamine (5-HT) 1B/1D receptor agonist, known to increase the degree of ectopic neural sprouting in *Xenopus* embryos^30^. Interestingly, three out of four of these neurobots showed an increase in movement complexity when treated with PTZ (filled circles Fig. 7b, Supp. Fig. 5). Moreover, these neurobots, showed a tighter correlation between the amount of implanted tissue and degree of innervation (Supp. Fig. 5). Interestingly, 5-HT_1B_ receptors are shown to modulate GABA release^48^, and serotonergic signaling is thought to affect the migration pattern of cortical interneurons^49^. Altered serotonergic signaling in neurobots treated with zolmitriptan may therefore impact the expression of GABAergic neurons during their development. Further experiments are needed to discover the mechanisms underlying this effect and whether zolmitriptan treatment results in biased expression of specific neural subtypes.

Our transcriptomics analysis revealed the landscape of differentially expressed genes between neurobots, biobots, and sham neurobots. Overall, neurobots showed a significant upregulation in gene expression compared to biobots and sham neurobots, whereas biobots and shams were more similar to one another (Fig. 9a,b, Fig. 10). Functional enrichment analysis revealed that neurobots, compared to both biobots and shams, exhibit a high level of enrichment in genes involved in nervous system development, synapse formation, neuron projection, and trans-synaptic signaling (Fig. 11a,b). Genes encoding major neurotransmitter receptors were present in the transcriptome of neurobots. This included glutamatergic, GABAergic, cholinergic, dopaminergic, serotonergic, and glycinergic receptors.

Our gene network analysis resulted in the identification of multiple functional clusters, allowing us to more deeply examine the genes and pathways that are upregulated in neurobots. Notably, we found a large cluster containing genes with important roles in visual perception (Cluster 1, Fig. 12c, Supp. Fig. 7c). This cluster contained genes normally expressed exclusively in *Xenopus* eyes, including various members of the opsin family such as a retinal G protein coupled receptor, various cone opsins, rhodopsin, as well as genes encoding many other proteins implicated in visual processing. This remarkable finding suggests the possible presence of visually evoked behaviors in neurobots. The next and most exciting step will be to test this hypothesis and discover the ways that light could modulate motor output in neurobots. If present, this will be a completely novel emergent behavior.

We showed that neurobots exhibit a significant increase in the variability of their gene counts compared to their non-neuronal counterparts (BB and SHs, Fig. 9c-f). The excess variability seen in neurobots relative to shams and biobots suggests that neurons may play an important role in guiding the way cells explore the gene expression landscape. The nervous systems of animals are known to influence the behavior of non-neural cells and tissues^39,50^, so it is possible that the information processing activity of the neurons—in response to the unique “life experiences” of individuals, or internally-generated spontaneous signaling—might account for the neurobots exhibiting the largest inter-individual gene expression variability of the groups. Moreover, studies show that when cells are exposed to novel stressors which they do not have existing homeostatic mechanisms to resolve, they resort to making random changes in the expression levels of many genes. It is possible that the bots we report here are undergoing stressors that evolution did not prepare them for, and may be employing this kind of exploration of gene expression space^51–55^.

Finally, based on a phylostratigraphic analysis, we show that the majority of upregulated genes in neurobots consist of the most ancient genes (Fig. 13), significantly different than is the case with biobots and shams. In all cases, the cells were wild-type–no genomic editing, synthetic biology circuits, scaffolds, or drugs were used. These results suggest that novel configurations of cell types can have large-scale systemic effects on the transcriptome of the resulting multicellular construct, and move it toward the gene expression profiles of the evolutionary past.

Building neurobots with predictable nervous system architecture and in large numbers is one of the major remaining challenges of this study. Future automation and standardization efforts will enable higher throughput and consistency, allowing repeated measurements to be performed on neurobots with both identical and distinct nervous system architectures. The impact of pharmacological, optical, and other types of stimulation on the neurobot behavior could be assessed. Additionally, the creation and neuroanatomical characterization of large numbers of identical bots will allow for the discovery of frequently emerging patterns and motifs, thereby shedding light on the potential space for nervous system architectures in novel living constructs whose precise layout has not been shaped by selection for function in this behavioral configuration. This study establishes a model system and experimental roadmap to increase our understanding of the plasticity of evolutionarily determined hardware of living beings to adapt on developmental (not evolutionary) timescales and to provide interfaces to bioengineered living constructs that may provide novel control capabilities for useful synthetic living machines and shed light on the origins of novelty in the evolution of nervous systems.

## Methods

### Construction of biobots, neurobots and shams

Biobots were constructed as described previously^17^, by excising tissue from the animal hemisphere of a Nieuwkoop and Faber stage 9 *Xenopus laevis* embryo (animal cap).

To construct neurobots and sham neurobots we excised 40-50 such animal caps and let them sit with external surface facing up in 60 mm petri dishes filled with a calcium magnesium free solution (50.3 mM NaCl, 0.7 mM KCl, 9.2 mM Na_2_HPO_4_, 0.9 mM KH_2_PO_4_, 2.4 mM NaHCO3, 1.0 mM edetic acid (EDTA), pH 7.3), and coated with 1% agarose made in the same solution. After about 30-40 minutes the cells were fully dissociated. The dissociated cells were transferred to a deep 60 mm petri dish containing 0.75X MMR, using a P200 pipette, taking as little liquid as possible. For constructing sham neurobots, the dissociated cells were immediately reaggregated and formed into clumps (see below). For constructing neurobots, we dispersed the dissociated cells as far as possible by moving the solution in the dish using a P1000 pipette. The cells were left still in the dish for ∼3-4 hours. To reaggregate cells, the dish containing dissociated cells was placed on a shaker and cells were thereby brought together in the middle of the dish. They were then allowed to reaggregate for approximately 1 hour, at which point various clumps of cells formed spontaneously.

Using a P1000 pipette, clumps of ∼ >10 cells were moved into the wells of an agarose coated 6 well plate. Larger clumps were broken into smaller ones that could fit inside the body of the bot, i.e. animal caps excised from a new set of embryos (see Fig. 1a). Depending on the size of the clumps, one or more were implanted. The variability in the size and number of the implanted neural precursor cell clumps (due to the manual nature of their creation), could be one of the factors contributing to the variability in neural expression, as we found a positive (though not significant) correlation between the relative size of the implanted clumps and resulting neurite length and number of neural terminals (see Fig. 8c).

Next, a new set of animal caps were dissociated from a second batch of embryos from a later fertilization (at late blastula, early gastrula stage), and were placed with external surface facing down individually in the wells of the same 6 well plate. The excised animal caps slowly formed a cup and eventually closed up within approximately 10-15 min. Clumps of neural precursor cells (or non-neuronal clumps in the case of shams) were placed inside this cup (Fig. 1aii) before it was closed using fine forceps, and enough time was allowed for the animal cap to fully close before moving the dish to the incubator.

### Immunohistochemistry

Bots were fixed overnight at 4 degrees Celsius in 4% paraformaldehyde (Thermo Fisher Scientific) with 0.25% Gluteraldehyde (Electron Microscopy Sciences) individually in 96 well plates. The next day, they were washed three times at room temperature in PBT (0.1% Triton X-100 (Sigma) in PBS-/- (Gibco)) for at least 15 minutes and then incubated in the 10% Casblock (invitrogen) dissolved in PBT for at least 1 hour. They were then transferred into the solution containing primary antibody (Anti-Acetylated Tubulin antibody, Mouse monoclonal, Sigma T7451) and Hoescht (33342, Thermo Scientific). The plate was sealed using parafilm and covered in foil for light protection and placed on a shaker in the cold room for 3 days at 4C. The bots were next washed 3 times in PBT at room temperature and then transferred and incubated overnight at 4C in the secondary antibody (goat anti-Mouse Alexa 594, Invitrogen A32742), with the dish sealed with parafilm and covered in foil. Finally, the bots were washed again 3 times in PBT and either stored in PBS at 4C or mounted into 15 μ-slide 18 well flat dishes (81821, Ibidi) in an antifade mounting medium (Vectashield) for confocal imaging.

### Calcium imaging

Embryos at the four-cell stage were microinjected in all four blastomeres with mRNA encoding the genetically encoded fluorescent calcium indicator GaCaMP6s. These embryos were used for obtaining clumps of neural precursor cells for implantation. Albino embryos were used as the outer shell of the neurobots in these experiments so that the fluorescent signals from neurons could be visualized more easily as wild type embryos are pigmented. We used a custom-built microscope to measure calcium activity in freely moving neurobots (Supp. Video 1). We used Fiji’s^60^ Descriptor Based Series Registration plugin to correct for the motion of the neurobot (Supp. Video 2), then used Suit2p software^61^ to identify active units (Fig. 3). To avoid movements in Z-direction, which resulted in changes in plane of focus, we created ‘flattened’ neurobots as follows: on the day after their formation neurobots were pressed down using a glass coverslip which was gradually lowered over them as small amounts of MMR were removed from the dish. Neurobots were left under pressure for 3 hours, after which MMR was gradually added to the dish resulting in the release of the coverslip.

### Quantification of the bot shape and neural tracing

We used the brush tool in Fiji^60^ to fill in the shape of the bot and calculated the area and roundness index defined as the major_axis / minor_axis. We used the same tool to estimate the relative amount of implanted neural precursor tissue by dividing the area of the implanted clumps to the outer shell (Fig. 1 a). We used Imaris software (Oxford Instruments) to quantify neural expression using confocal stacks acquired from the bots that were stained with antibodies against acetylated alpha tubulin, which labeled neurons and cilia in multiciliated cells. We manually traced neurites using the filament function and exported values corresponding to the total length of neurites (dendrite length sum parameter in Imaris) and the number of terminal points (number of dendrite terminal points parameter in Imaris). For the analysis of the multiciliated cell distribution, we estimated the total number of MCCs by marking the center of each MCC using the Spots tool in Imaris and calculated the total number of multiciliated cells. We then used this value to calculate the MCC density by dividing this number by the total area of the bot.

### Behavioral analysis

Videos of bot movements were taken over 30 minutes under various conditions and tracked with the DLTdv digitizing tool^28^ in MATLAB and the *x* and *y* coordinates of the center of mass were calculated. A custom Python code was used to extract various kinematic variables using the time series of the coordinates. We calculate total Euclidean distance travelled, speed, and acceleration. Additionally, we calculated the percentage of the well that was traversed by the bots by dividing the space of each well into 0.1 mm bins. We then calculated percent covered area by dividing the number of unique visited bins by the total number of bins. We calculated a complexity index by first calculating the power spectral density (PSD) of the trajectory time series along the *x* and *y* coordinates and identifying peaks in the power. We calculated Welch’s power spectral density estimate with a window size of 400s and overlap of 1s between windows for each of the *x* and *y* time series. We picked a threshold of 10 pixels^2^/Hz (∼0.17 mm^2^/Hz= 0.4 mm/Hz) to detect peaks in the PSD of the *x* and *y* coordinates. This threshold was chosen to remove the baseline noise corresponding to the tracking of the center of mass of bots that had an average radius of 0.4 mm. We then defined the complexity index as the total number of unique peaks in *x* and *y* PSDs.

### RNA-sequencing and bioinformatics

We submitted 12 samples (4 samples per biobot type, 5-15 biobots per sample) submerged in Trizol (Invitrogen) in 2 mL Eppendorf tubes to Novogene for low-input, high-lipid, bulk RNA extraction. Equal quantities of RNA were sequenced from each sample using the NovaSeq6000 sequencer, resulting in consistent library size across samples. Clean reads were extracted from FASTQ files, removing reads with adapter contamination, when uncertain nucleotides constitute more than 10 percent of either read (N > 10%), and when low quality nucleotides (Base Quality less than 5) constitute more than 50 percent of the read. The index to the reference genome (*Xenopus laevis* version 10.1) was built using Hisat2 v2.0.5^62^ and clean reads were aligned to the reference. The mapped reads of each sample were assembled using StringTie (v1.3.3b)^63^ and FeatureCounts v1.5.0-p3^64^ was used to count the reads numbers mapped to each gene. FPKM of each gene was calculated based on the length of the gene and read counts mapped to this gene. Differential expression analysis was performed using the DESeq2 R package (1.20.0)^65^ and the resulting p-values were adjusted using Benjamini and Hochberg’s approach for controlling the false discovery rate.

The webapp g:Profiler^41^ was used to perform functional enrichment analysis of differentially expressed genes. For each comparison, upregulated genes (p-adjusted < 0.05; log2foldchange > 4) and downregulated genes (p-adjusted < 0.05; log2foldchange < 4) were separately mapped from *Xenopus* to human symbols using the HGNC Comparison of Orthology Predictions (HCOP) tool^66^. Genes lacking an established gene symbol were removed from analysis. Each gene list was separately queried using g:Profiler across all data sources. The statistical data scope included only annotated genes and the g:SCS method was used for computing multiple testing corrections for p-values at a threshold of p<0.05. The R package ggplot2 (v3.5.1)^67^ was used to generate dot plots of gene ontology driver terms from g:Profiler. Driver terms were determined by grouping significant terms into sub-ontologies based on their relations, then identifying the leading gene sets that give rise to other significant functions in the ontology neighborhood.

For network analysis and clustering we applied network analysis techniques to discover biological functional modules^68,69^. By integrating gene expression and interaction data, we extracted protein-protein interactions (PPI) for the different biobots in their respective conditions and applied network embedding and clustering techniques similar to those described by Cantini et al. (2015)^70^ and Pio-Lopez et al. (2021)^71^. Specifically, we used the MNMF algorithm developed by Wang et al. (2017)^40^ for network embedding and clustering. To create a network for the biobots, we started by isolating genes of interest and identifying corresponding human orthologs under various conditions using the HCOP database^72^. We then used the STRING database^73^ to extract relevant PPI networks. The clusters identified through our network embedding and clustering method were further analyzed for enrichment using g:Profiler^41^.

### Analysis of gene expression variability

The normalized gene count variability was compared between groups (BBs, NBs, and SHs) using a MATLAB script and the method summarized in Supp. Fig. 9. For each gene in each pair of groups being compared, the mean count value across all pools of both groups was found, and genes were ranked from greatest to least mean. Genes for which any of the counts across all pools was 0 were discarded. Because the count value of a given gene in a given pool represents the mean value of all the individuals in that pool, the standard deviation of the pools gives the standard error of the means (SE) of the group. The SE is related to the number of individuals per pool (n) and the standard deviation of the individuals within the pools (σ) with the equation SE = σ/sqrt(n). By multiplying the SE by sqrt(n), σ can be calculated. Dividing σ by the mean count value of the pools gives the coefficient of variation (CV). The ranked gene CV lists were then split into 100 bins (percentiles) containing equal numbers of genes from highest to lowest counts. Within each bin, the number of genes with greater CV for the first group than the second were counted and divided by bin size to find the fraction of genes in the bin with greater CV in the first group. These fractions were then plotted as bar graphs in Figure 9c-e, with a blue line marking 0.5. The bin values appeared to vary with gene count percentile, so to determine the statistical significance of each bin’s departure from its expected value, a permutation test was used. For each plot (each pair of groups), the order of the gene pairs was randomly shuffled (keeping pairs together), and new bins were generated. This was repeated 1,000 times for random shuffles to produce a distribution of bin fraction values for each bin. The p-value of each bin was defined as the proportion of the bin fractions from the distribution that were further in absolute value from the distribution mean than the true bin fraction. Bins with p-values of p<0.05 were deemed statistically significant and were colored dark blue. All other bins were colored light blue. The mean bin value from the distributions was plotted as a yellow line.

### Phylostratigraphic analysis

We employed the phylostratR package (Arendsi et al., 2019) to conduct a phylostratigraphic analysis on neurobot transcripts, with *Xenopus* laevis (taxon ID ‘8355’) designated as the reference species. This software automates several key steps in evolutionary analysis: (1) it builds a clade tree using species from the UniProt database and aligns it with the latest NCBI taxonomy; (2) the clade tree is trimmed to maintain a phylogenetically diverse selection of representative species for each phylostratum; (3) a comprehensive protein sequence database is constructed from hundreds of species based on this clade tree, with additional data such as human and yeast proteomes manually added, resulting in 329 species for our study; (4) a similarity search is performed by conducting pairwise BLAST comparisons between the proteins encoded by *Xenopus laevis* and those of the target species; (5) the best hits are identified, and gene homology is inferred between *Xenopus laevis* and the target species; (6) each gene is assigned to a phylostratum that corresponds to the oldest clade for which a homolog is identified. Genes specific to *Xenopus laevis* are classified as orphan genes and placed within the *Xenopus laevis* phylostratum. The evolutionary stages we focused on include: All living organisms (bacteria, eubacteria), Eukaryota, Opisthokonta, Metazoa, Eumetazoa, Bilateria, Deuterostomia, Chordata, Vertebrata, Gnathostomata, Euteleostomi, Sarcopterygii, Tetrapoda, Anura, Xenopus, and *Xenopus laevis*. This methodological approach enables a detailed examination of gene emergence and their evolutionary trajectories across various taxa. By implementing Phylostratr, we systematically mapped the age of the neurobot genes in the different conditions with a specific phylostrata to understand the distribution of ages of the neurobots’ overexpressed genes. We used the upregulated and downregulated genes in neurobots (logFC>4 and logFC<-2 respectively).

## Acknowledgements

We would like to thank Drs. Douglas Blackiston, Patrick McMillen and Pai Vaibhav for their technical help and scientific insights, Meghan Short for statistical analysis consultation, Susan Marquez, Thomas Ferrante, Ramses Martinez, and Kostyantyn Shcherbina for their help with engineering, Jeantine Lunshof for philosophical and ethics discussions, and Drs. Donald Ingber and Michael Super for their support throughout the project. Thanks to Gordon Allen for copyediting and Julia Poirier for assistance with the manuscript. This research was supported by HR0011-18-2-0022 and W911NF1920027 both awarded by the Department of Defense, and grants from John Templeton Foundation and Northpond Ventures.

## Notes

### Competing Interest Statement

The authors have declared no competing interest.

