## Supplemental Figures for "Self-Organizing Neural Networks in Novel Moving Bodies: Anatomical, Behavioral, and Transcriptional Characterization of a Living Construct with a Nervous System"

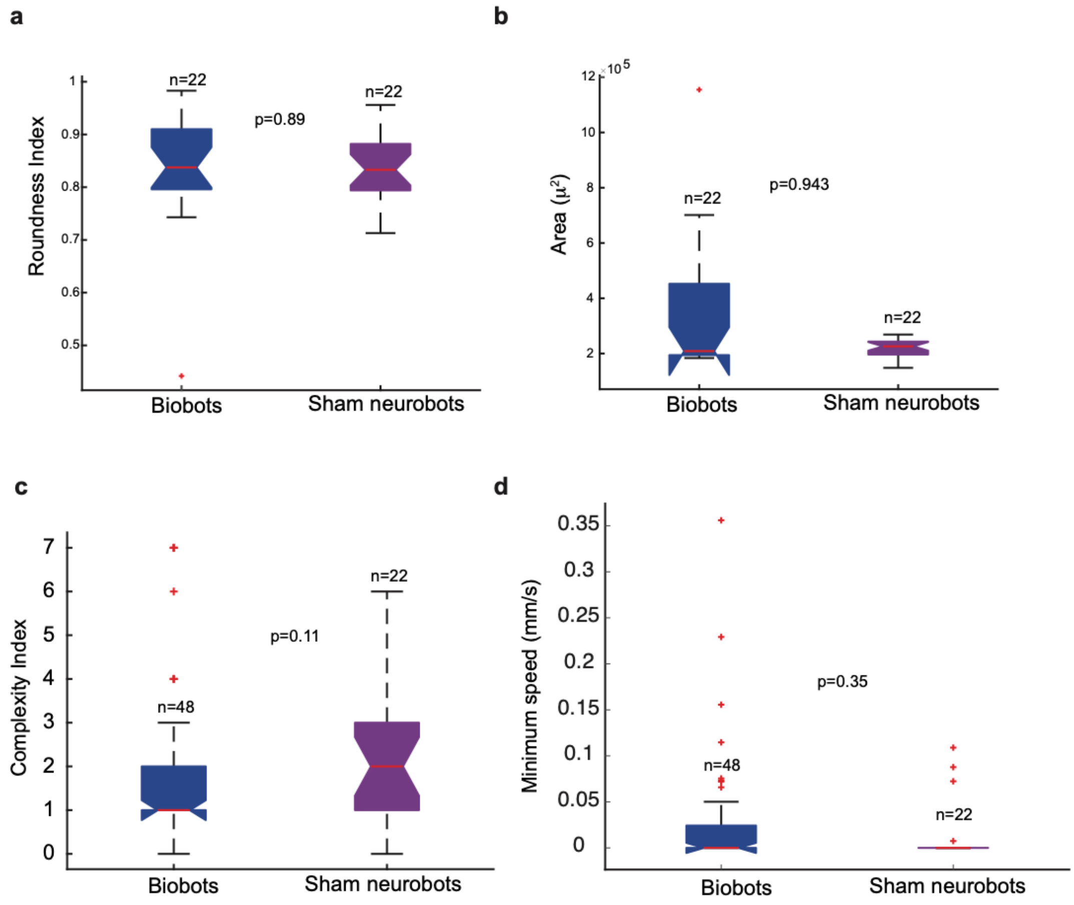
a
Roundness Index
n=22
n=22
p=0.89
Biobots
Sham neurobots
b
Area (μ²)
n=22
n=22
p=0.943
Biobots
Sham neurobots
c
Complexity Index
n=48
n=22
p=0.11
Biobots
Sham neurobots
d
Minimum speed (mm/s)
n=48
n=22
p=0.35
Biobots
Sham neurobots

**Figure S1**

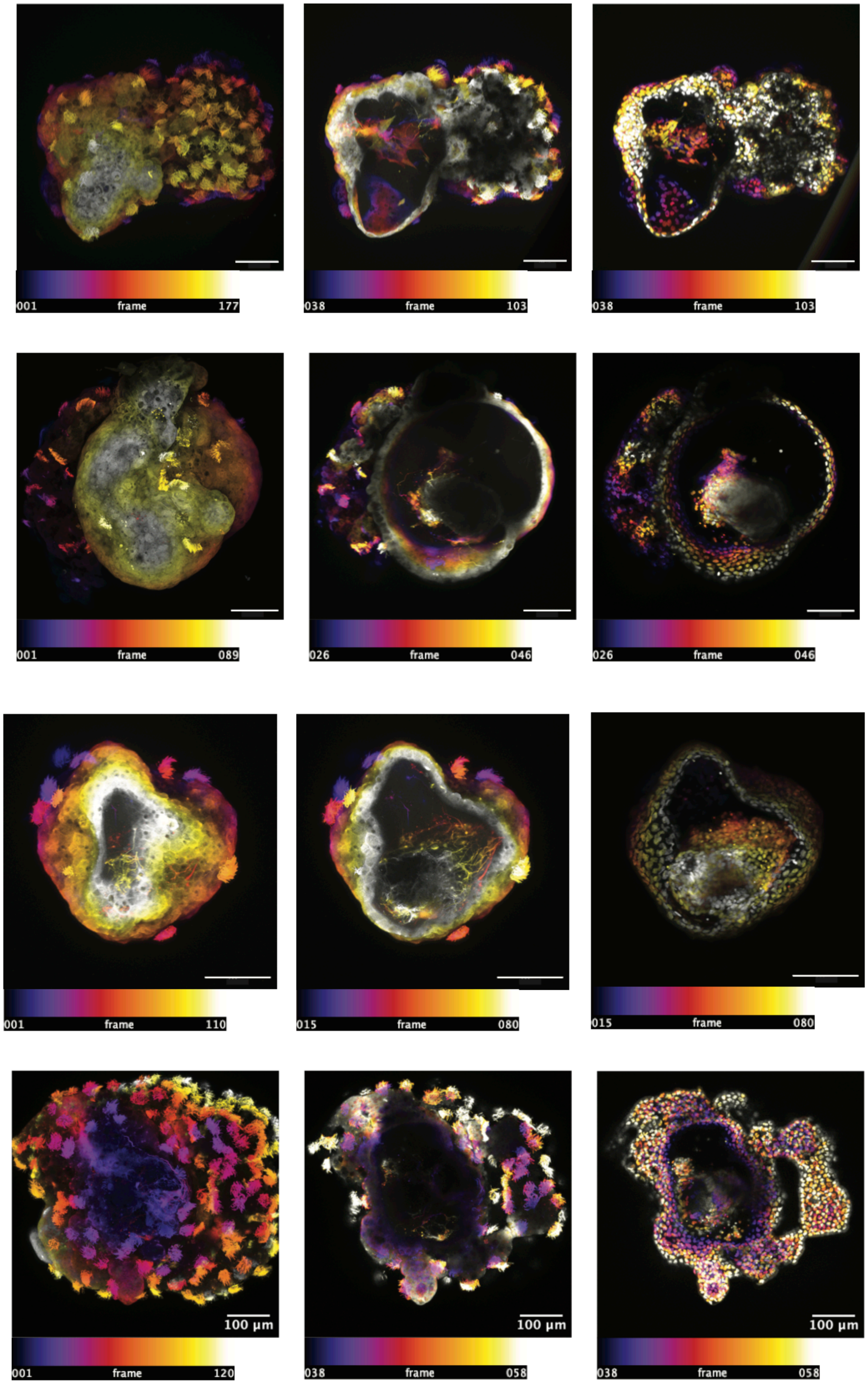
001
frame
177
038
frame
103
038
frame
103
001
frame
089
026
frame
046
026
frame
046
001
frame
110
015
frame
080
015
frame
080
100 µm
001
frame
120
100 µm
038
frame
058
100 µm
038
frame
058

**Figure S2**

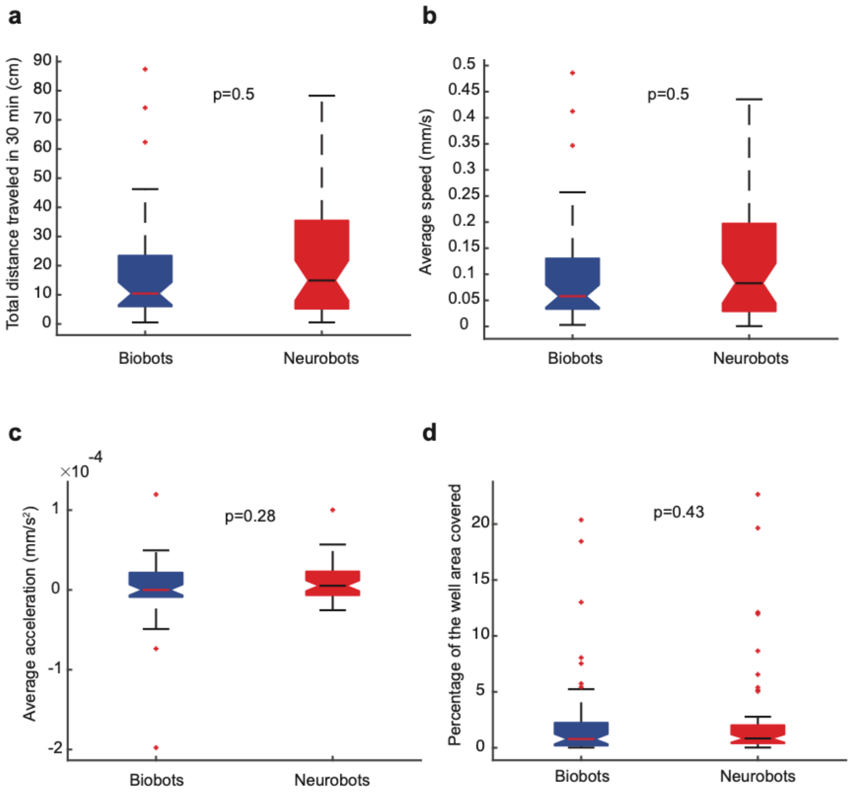

a
b
c
d
Total distance traveled in 30 min (cm)
p=0.5
Average speed (mm/s)
p=0.5
Average acceleration (mm/s²)
×10^-4
p=0.28
Percentage of the well area covered
p=0.43
Biobots
Neurobots


**Figure S3**

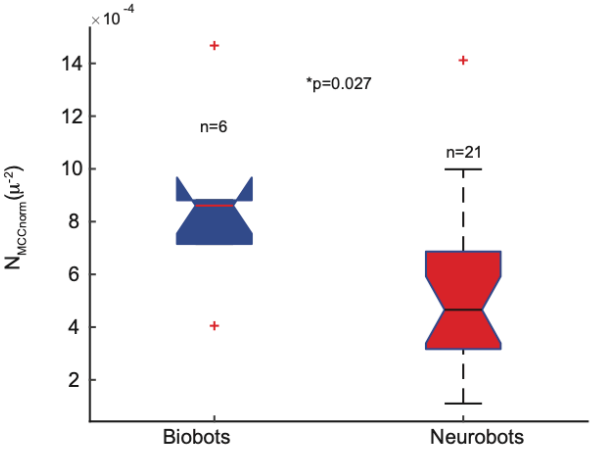
×10 -4
14
12
10
8
6
4
2
$N_{MCCnorm}(\mu^{-2})$
n=6
*p=0.027
n=21
Biobots
Neurobots

**Figure S4**

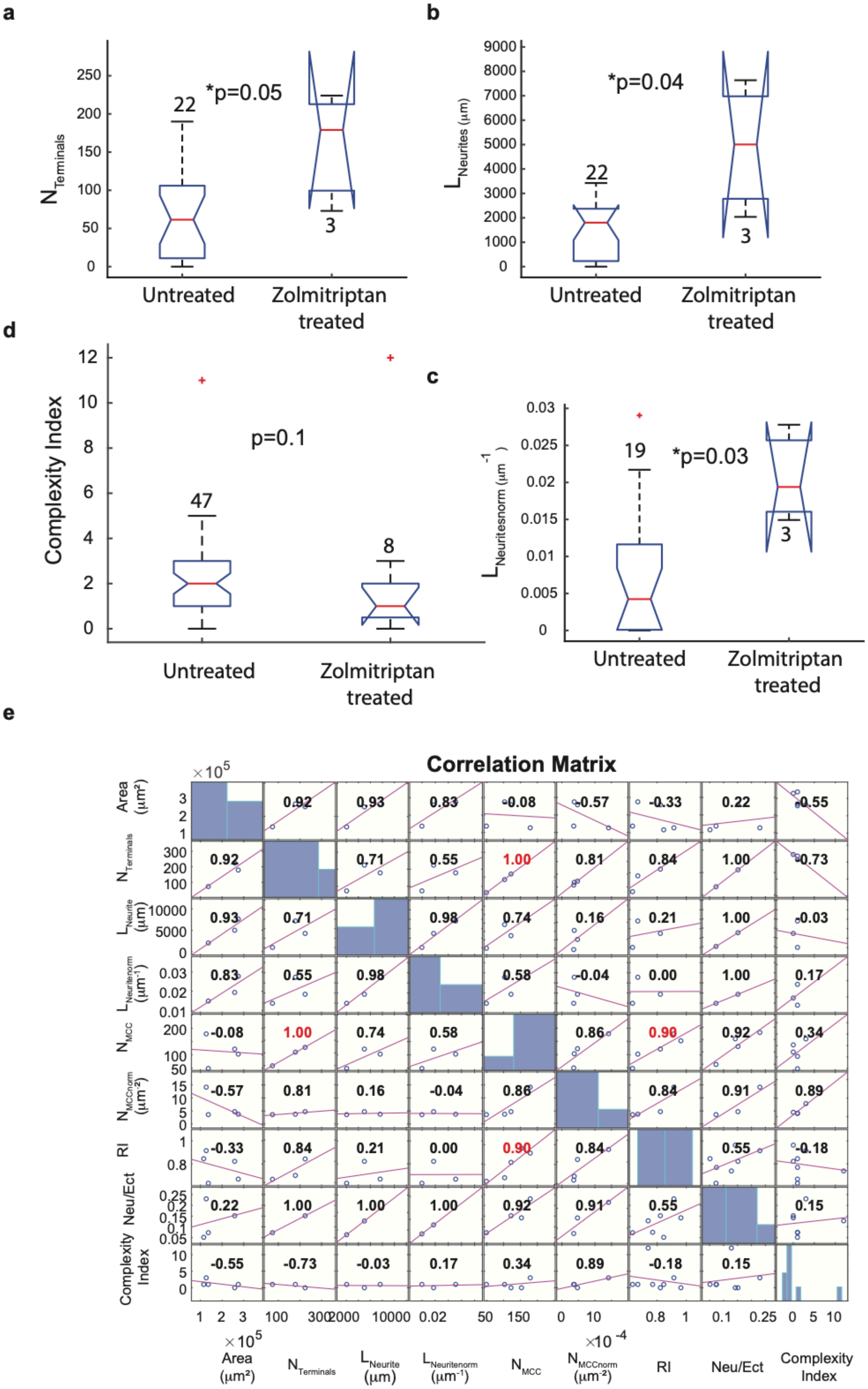
a
b
d
c
e
*p=0.05
*p=0.04
p=0.1
*p=0.03
Untreated
Zolmitriptan treated
Complexity Index
Correlation Matrix
Area (µm²)
RI
Neu/Ect
Complexity Index

**Figure S5**

a

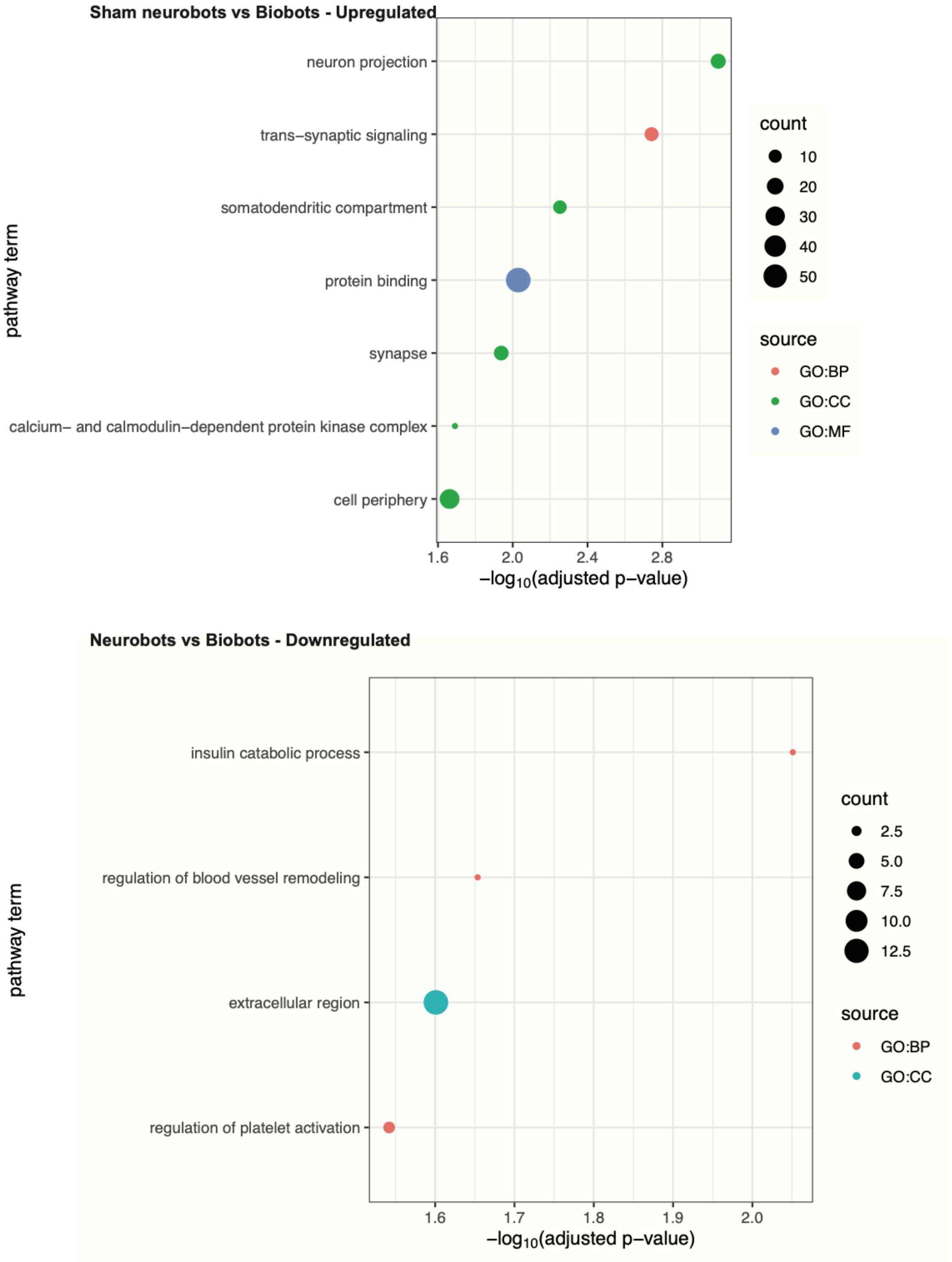

Sham neurobots vs Biobots - Upregulated
neuron projection
trans−synaptic signaling
somatodendritic compartment
protein binding
synapse
calcium− and calmodulin−dependent protein kinase complex
cell periphery
pathway term
1.6
2.0
2.4
2.8
−log10(adjusted p−value)
count
10
20
30
40
50
source
GO:BP
GO:CC
GO:MF
Neurobots vs Biobots - Downregulated
insulin catabolic process
regulation of blood vessel remodeling
extracellular region
regulation of platelet activation
pathway term
1.6
1.7
1.8
1.9
2.0
−log10(adjusted p−value)
count
2.5
5.0
7.5
10.0
12.5
source
GO:BP
GO:CC


**Figure S6**

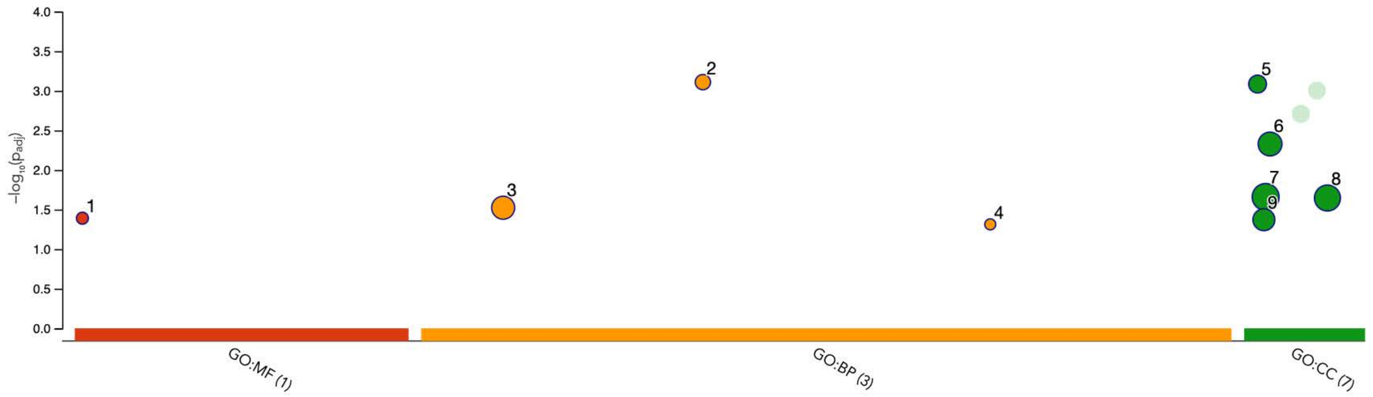


| ID | Source | Term ID | Term Name | $p_{adj}$ (query_1) |
|---|---|---|---|---|
| 1 | GO:MF | GO:0001849 | complement component C1q complex binding | $4.066\times10^{-2}$ |
| 2 | GO:BP | GO:0035633 | maintenance of blood-brain barrier | $7.771\times10^{-4}$ |
| 3 | GO:BP | GO:0007155 | cell adhesion | $2.998\times10^{-2}$ |
| 4 | GO:BP | GO:0097241 | hematopoietic stem cell migration to bone mar... | $4.864\times10^{-2}$ |
| 5 | GO:CC | GO:0005923 | bicellular tight junction | $8.224\times10^{-4}$ |
| 6 | GO:CC | GO:0030054 | cell junction | $4.701\times10^{-3}$ |
| 7 | GO:CC | GO:0016020 | membrane | $2.195\times10^{-2}$ |
| 8 | GO:CC | GO:0071944 | cell periphery | $2.260\times10^{-2}$ |
| 9 | GO:CC | GO:0009986 | cell surface | $4.249\times10^{-2}$ |

**Cluster 2**

| | | |
|---|---|---|
| **version** | e111_eg58_p18_f463989d | g:Profiler |
| **date** | 8/26/2024, 6:27:48 PM | |
| **organism** | hsapiens | |

**Figure S7a**

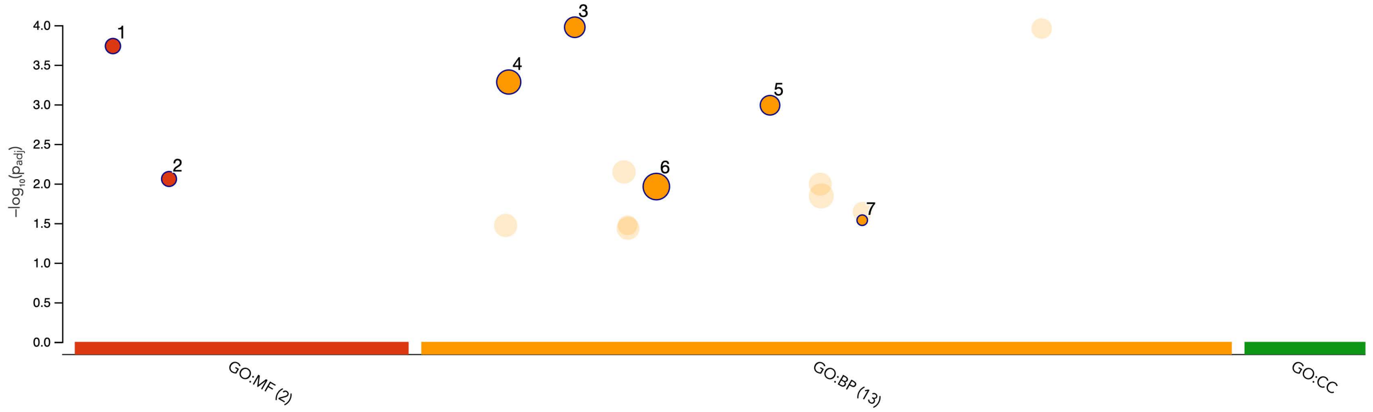


| ID | Source | Term ID | Term Name | $p_{adj}$ (query_1) |
|---|---|---|---|---|
| 1 | GO:MF | GO:0005109 | frizzled binding | $1.837\times10^{-4}$ |
| 2 | GO:MF | GO:0017147 | Wnt-protein binding | $8.767\times10^{-3}$ |
| 3 | GO:BP | GO:0016055 | Wnt signaling pathway | $1.063\times10^{-4}$ |
| 4 | GO:BP | GO:0007399 | nervous system development | $5.230\times10^{-4}$ |
| 5 | GO:BP | GO:0045165 | cell fate commitment | $1.027\times10^{-3}$ |
| 6 | GO:BP | GO:0032501 | multicellular organismal process | $1.093\times10^{-2}$ |
| 7 | GO:BP | GO:0060061 | Spemann organizer formation | $2.908\times10^{-2}$ |

**Cluster 3**

| | |
|---|---|
| **version** | e111_eg58_p18_f463989d |
| **date** | 8/26/2024, 3:56:14 PM |
| **organism** | hsapiens |

g:Profiler

**Figure S7b**

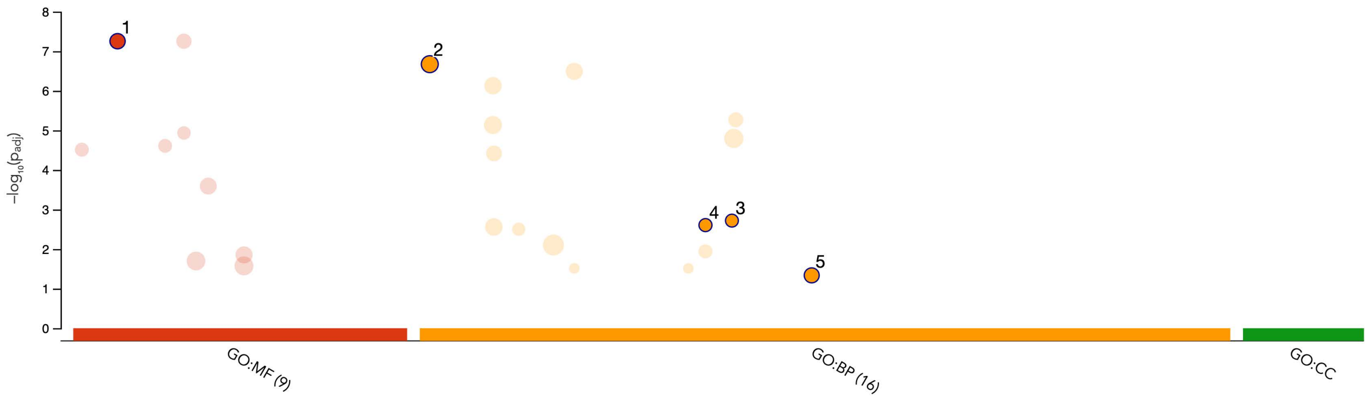


| ID | Source | Term ID | Term Name | $p_{adj}$ (query_... |
|---|---|---|---|---|
| 1 | GO:MF | GO:0005501 | retinoid binding | $5.584\times10^{-8}$ |
| 2 | GO:BP | GO:0001523 | retinoid metabolic process | $2.124\times10^{-7}$ |
| 3 | GO:BP | GO:0042363 | fat-soluble vitamin catabolic process | $1.909\times10^{-3}$ |
| 4 | GO:BP | GO:0035810 | positive regulation of urine volume | $2.479\times10^{-3}$ |
| 5 | GO:BP | GO:0048384 | retinoic acid receptor signaling pathway | $4.613\times10^{-2}$ |

**Cluster 4**

**version** e111_eg58_p18_f463989d
**date** 9/6/2024, 6:36:42 PM
**organism** hsapiens

g:Profiler

**Figure S7c**

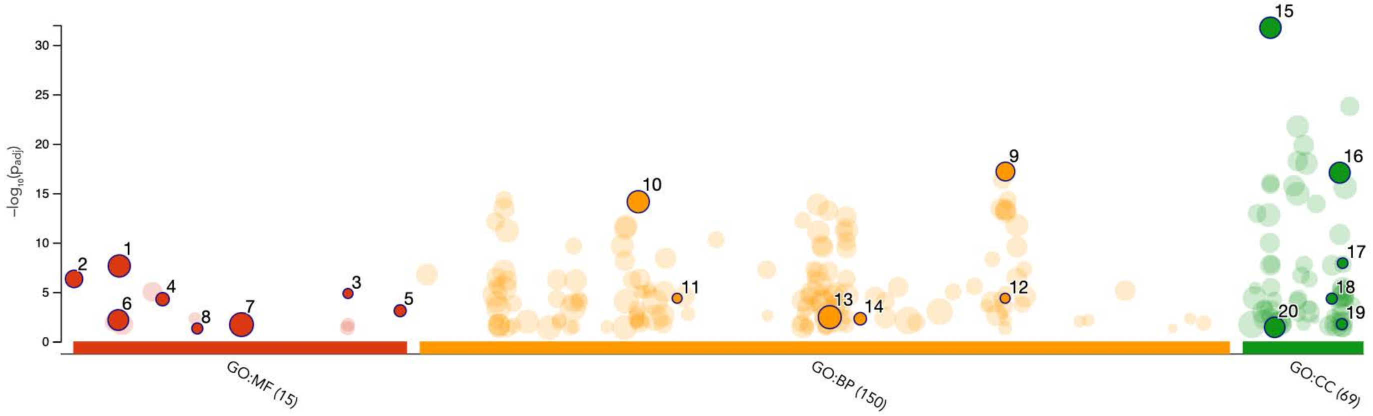


| ID | Source | Term ID | Term Name | $p_{adj}$ (query_1) |
|---|---|---|---|---|
| 1 | GO:MF | GO:0008092 | cytoskeletal protein binding | $2.330\times10^{-8}$ |
| 2 | GO:MF | GO:0000149 | SNARE binding | $4.758\times10^{-7}$ |
| 3 | GO:MF | GO:0099184 | structural constituent of postsynaptic intermediate filament cytoskeleton | $1.429\times10^{-5}$ |
| 4 | GO:MF | GO:0016812 | hydrolase activity, acting on carbon-nitrogen (but not peptide) bonds, in cyclic amides | $5.370\times10^{-5}$ |
| 5 | GO:MF | GO:1903136 | cuprous ion binding | $7.920\times10^{-4}$ |
| 6 | GO:MF | GO:0005543 | phospholipid binding | $6.792\times10^{-3}$ |
| 7 | GO:MF | GO:0042802 | identical protein binding | $2.014\times10^{-2}$ |
| 8 | GO:MF | GO:0031694 | alpha-2A adrenergic receptor binding | $4.952\times10^{-2}$ |
| 9 | GO:BP | GO:0099504 | synaptic vesicle cycle | $6.512\times10^{-18}$ |
| 10 | GO:BP | GO:0031175 | neuron projection development | $7.439\times10^{-15}$ |
| 11 | GO:BP | GO:0033693 | neurofilament bundle assembly | $4.327\times10^{-5}$ |
| 12 | GO:BP | GO:0099185 | postsynaptic intermediate filament cytoskeleton organization | $4.327\times10^{-5}$ |
| 13 | GO:BP | GO:0050877 | nervous system process | $3.548\times10^{-3}$ |
| 14 | GO:BP | GO:0060052 | neurofilament cytoskeleton organization | $5.122\times10^{-3}$ |
| 15 | GO:CC | GO:0030424 | axon | $1.837\times10^{-32}$ |
| 16 | GO:CC | GO:0098793 | presynapse | $8.544\times10^{-18}$ |
| 17 | GO:CC | GO:0099160 | postsynaptic intermediate filament cytoskeleton | $1.221\times10^{-8}$ |
| 18 | GO:CC | GO:0097418 | neurofibrillary tangle | $4.716\times10^{-5}$ |
| 19 | GO:CC | GO:0099012 | neuronal dense core vesicle membrane | $1.794\times10^{-2}$ |
| 20 | GO:CC | GO:0031252 | cell leading edge | $3.806\times10^{-2}$ |

**Cluster 5**

| | |
|---|---|
| **version** | e111_eg58_p18_f463989d |
| **date** | 8/26/2024, 6:29:09 PM |
| **organism** | hsapiens |

g:Profiler

**Figure S7d**

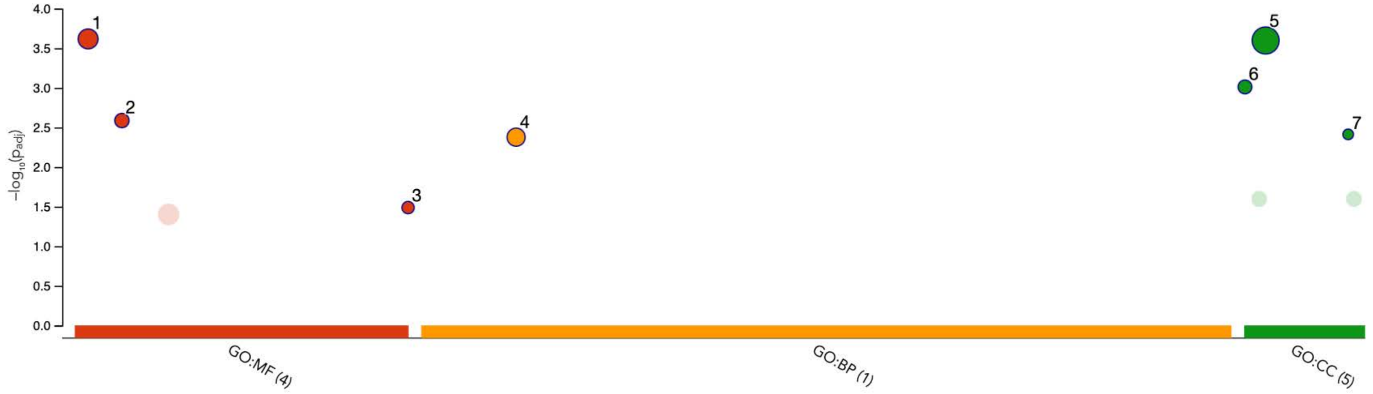


| ID | Source | Term ID | Term Name | $p_{adj}$ (query_1) |
|---|---|---|---|---|
| 1 | GO:MF | GO:0003924 | GTPase activity | $2.411\times10^{-4}$ |
| 2 | GO:MF | GO:0008157 | protein phosphatase 1 binding | $2.588\times10^{-3}$ |
| 3 | GO:MF | GO:2001069 | glycogen binding | $3.256\times10^{-2}$ |
| 4 | GO:BP | GO:0008277 | regulation of G protein-coupled receptor signaling pathway | $4.199\times10^{-3}$ |
| 5 | GO:CC | GO:0016020 | membrane | $2.525\times10^{-4}$ |
| 6 | GO:CC | GO:0000164 | protein phosphatase type 1 complex | $9.741\times10^{-4}$ |
| 7 | GO:CC | GO:0120216 | matrilin complex | $3.871\times10^{-3}$ |

**Cluster 6**

| | |
|---|---|
| **version** | e111_eg58_p18_f463989d |
| **date** | 8/26/2024, 6:29:48 PM |
| **organism** | hsapiens |

g:Profiler

**Figure S7e**

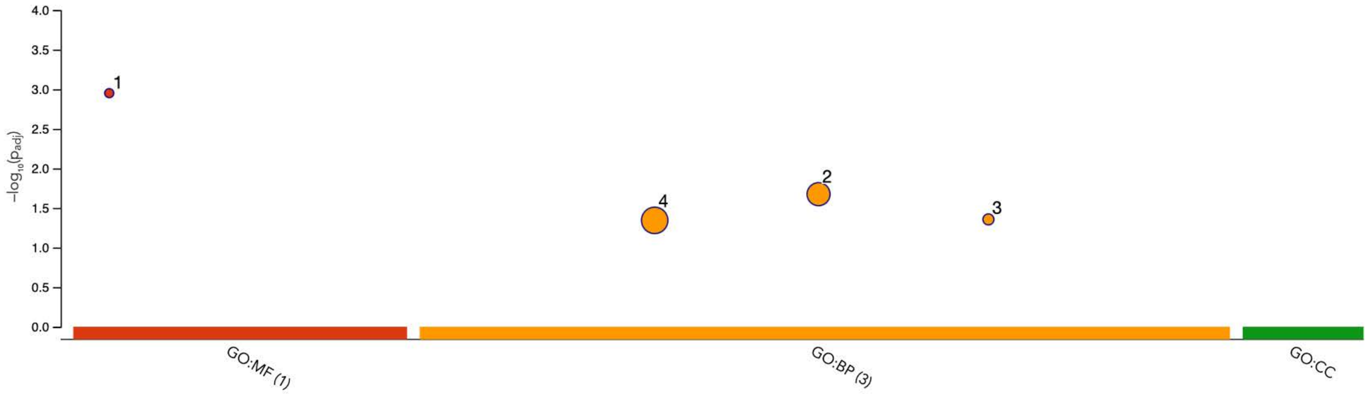


| ID | Source | Term ID | Term Name | $p_{adj}$ (query_1) |
|---|---|---|---|---|
| 1 | GO:MF | GO:0004968 | gonadotropin-releasing hormone receptor activity | $1.120\times10^{-3}$ |
| 2 | GO:BP | GO:0048699 | generation of neurons | $2.115\times10^{-2}$ |
| 3 | GO:BP | GO:0097211 | cellular response to gonadotropin-releasing hormone | $4.408\times10^{-2}$ |
| 4 | GO:BP | GO:0032501 | multicellular organismal process | $4.534\times10^{-2}$ |

**Cluster 8**

| | |
|---|---|
| **version** | e111_eg58_p18_f463989d |
| **date** | 8/26/2024, 6:30:42 PM |
| **organism** | hsapiens |

g:Profiler

**Figure S7f**

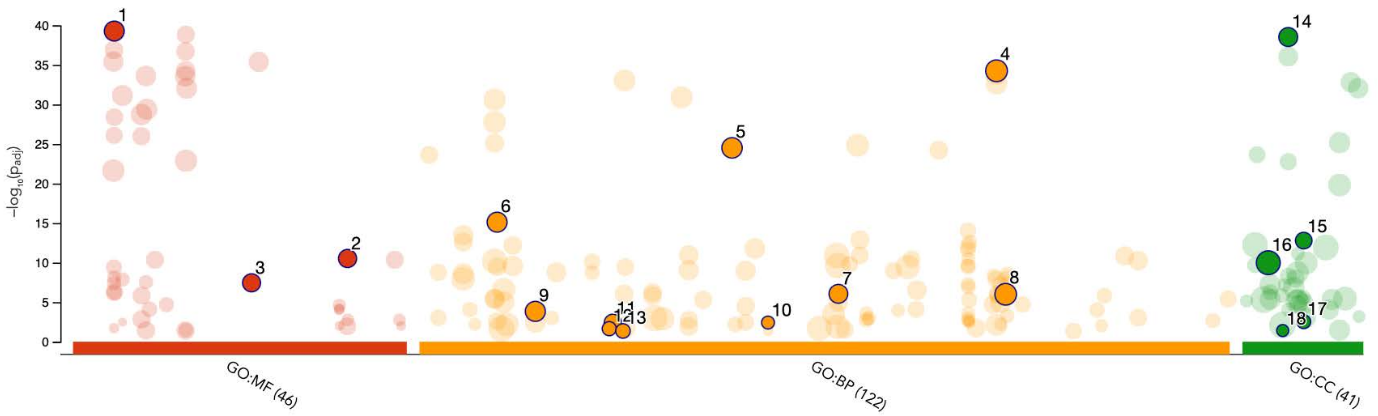


| ID | Source | Term ID | Term Name | $p_{adj}$ (query_1) |
|---|---|---|---|---|
| 1 | GO:MF | GO:0005261 | monoatomic cation channel activity | $5.245\times10^{-40}$ |
| 2 | GO:MF | GO:0099106 | ion channel regulator activity | $2.994\times10^{-11}$ |
| 3 | GO:MF | GO:0044325 | transmembrane transporter binding | $3.579\times10^{-8}$ |
| 4 | GO:BP | GO:0098662 | inorganic cation transmembrane transport | $5.555\times10^{-35}$ |
| 5 | GO:BP | GO:0042391 | regulation of membrane potential | $3.102\times10^{-25}$ |
| 6 | GO:BP | GO:0006936 | muscle contraction | $7.677\times10^{-16}$ |
| 7 | GO:BP | GO:0051260 | protein homooligomerization | $8.827\times10^{-7}$ |
| 8 | GO:BP | GO:0099537 | trans-synaptic signaling | $1.011\times10^{-6}$ |
| 9 | GO:BP | GO:0010038 | response to metal ion | $1.462\times10^{-4}$ |
| 10 | GO:BP | GO:0045161 | neuronal ion channel clustering | $3.782\times10^{-3}$ |
| 11 | GO:BP | GO:0021675 | nerve development | $4.125\times10^{-3}$ |
| 12 | GO:BP | GO:0021554 | optic nerve development | $2.254\times10^{-2}$ |
| 13 | GO:BP | GO:0022038 | corpus callosum development | $4.372\times10^{-2}$ |
| 14 | GO:CC | GO:0034703 | cation channel complex | $2.989\times10^{-39}$ |
| 15 | GO:CC | GO:0044304 | main axon | $1.640\times10^{-13}$ |
| 16 | GO:CC | GO:0030054 | cell junction | $1.030\times10^{-10}$ |
| 17 | GO:CC | GO:0044305 | calyx of Held | $2.989\times10^{-3}$ |
| 18 | GO:CC | GO:0033010 | paranodal junction | $3.884\times10^{-2}$ |

**Cluster 9**

**version** e111_eg58_p18_f463989d
**date** 8/26/2024, 6:31:40 PM
**organism** hsapiens

g:Profiler

**Figure S7g**

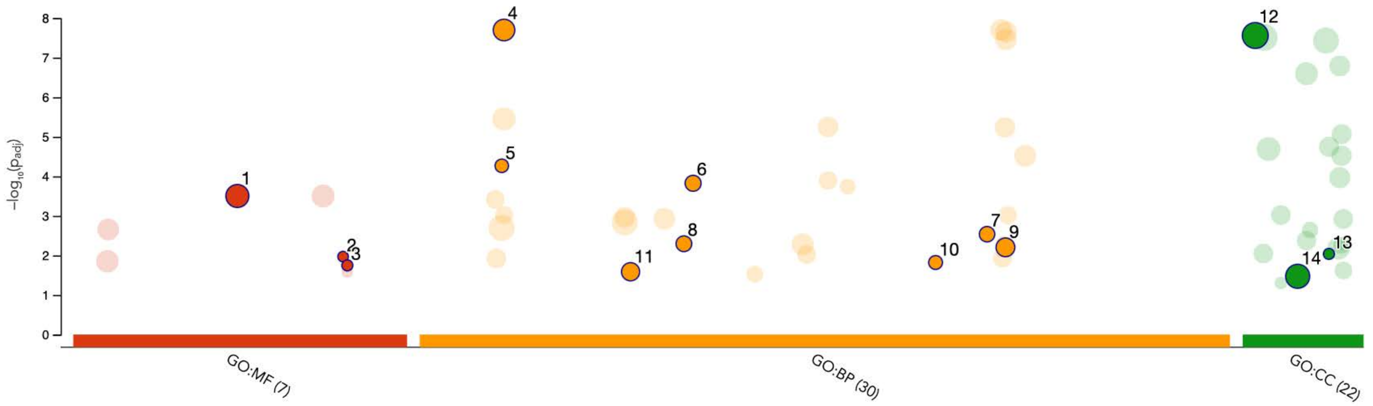


| ID | Source | Term ID | Term Name | $p_{adj}$ (query_1) |
|---|---|---|---|---|
| 1 | GO:MF | GO:0038023 | signaling receptor activity | $3.139\times10^{-4}$ |
| 2 | GO:MF | GO:0097016 | L27 domain binding | $1.073\times10^{-2}$ |
| 3 | GO:MF | GO:0098919 | structural constituent of postsynaptic density | $1.785\times10^{-2}$ |
| 4 | GO:BP | GO:0007268 | chemical synaptic transmission | $1.998\times10^{-8}$ |
| 5 | GO:BP | GO:0007158 | neuron cell-cell adhesion | $5.415\times10^{-5}$ |
| 6 | GO:BP | GO:0035176 | social behavior | $1.500\times10^{-4}$ |
| 7 | GO:BP | GO:0097106 | postsynaptic density organization | $2.903\times10^{-3}$ |
| 8 | GO:BP | GO:0034331 | cell junction maintenance | $5.068\times10^{-3}$ |
| 9 | GO:BP | GO:0099504 | synaptic vesicle cycle | $6.241\times10^{-3}$ |
| 10 | GO:BP | GO:0071625 | vocalization behavior | $1.508\times10^{-2}$ |
| 11 | GO:BP | GO:0030534 | adult behavior | $2.586\times10^{-2}$ |
| 12 | GO:CC | GO:0005886 | plasma membrane | $2.749\times10^{-8}$ |
| 13 | GO:CC | GO:0097025 | MPP7-DLG1-LIN7 complex | $9.148\times10^{-3}$ |
| 14 | GO:CC | GO:0042995 | cell projection | $3.369\times10^{-2}$ |

**Cluster 10**

| | |
|---|---|
| **version** | e111_eg58_p18_f463989d |
| **date** | 8/26/2024, 6:33:19 PM |
| **organism** | hsapiens |

g:Profiler

**Figure S7h**

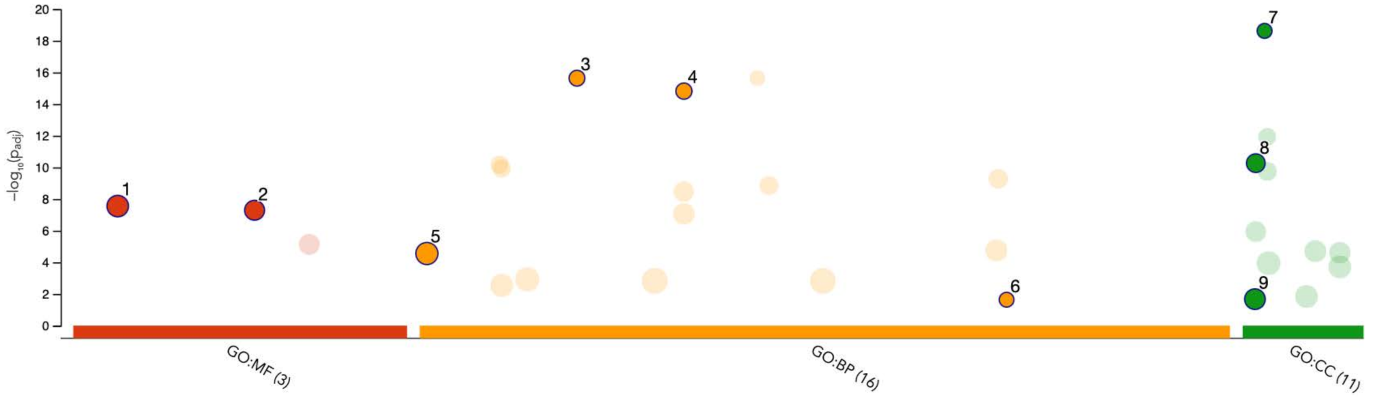


| ID | Source | Term ID | Term Name | $p_{adj}$ (query_1) |
|---|---|---|---|---|
| 1 | GO:MF | GO:0005509 | calcium ion binding | $2.799 \times 10^{-8}$ |
| 2 | GO:MF | GO:0045296 | cadherin binding | $5.126 \times 10^{-8}$ |
| 3 | GO:BP | GO:0016339 | calcium-dependent cell-cell adhesion via plasma membrane cell adhesion molecules | $2.319 \times 10^{-16}$ |
| 4 | GO:BP | GO:0034332 | adherens junction organization | $1.517 \times 10^{-15}$ |
| 5 | GO:BP | GO:0000902 | cell morphogenesis | $2.781 \times 10^{-5}$ |
| 6 | GO:BP | GO:0099560 | synaptic membrane adhesion | $2.316 \times 10^{-2}$ |
| 7 | GO:CC | GO:0016342 | catenin complex | $2.373 \times 10^{-19}$ |
| 8 | GO:CC | GO:0005912 | adherens junction | $5.419 \times 10^{-11}$ |
| 9 | GO:CC | GO:0005874 | microtubule | $2.126 \times 10^{-2}$ |

**Cluster 11**

| | |
|---|---|
| **version** | e111_eg58_p18_f463989d |
| **date** | 8/26/2024, 6:32:43 PM |
| **organism** | hsapiens |

g:Profiler

**Figure S7i**

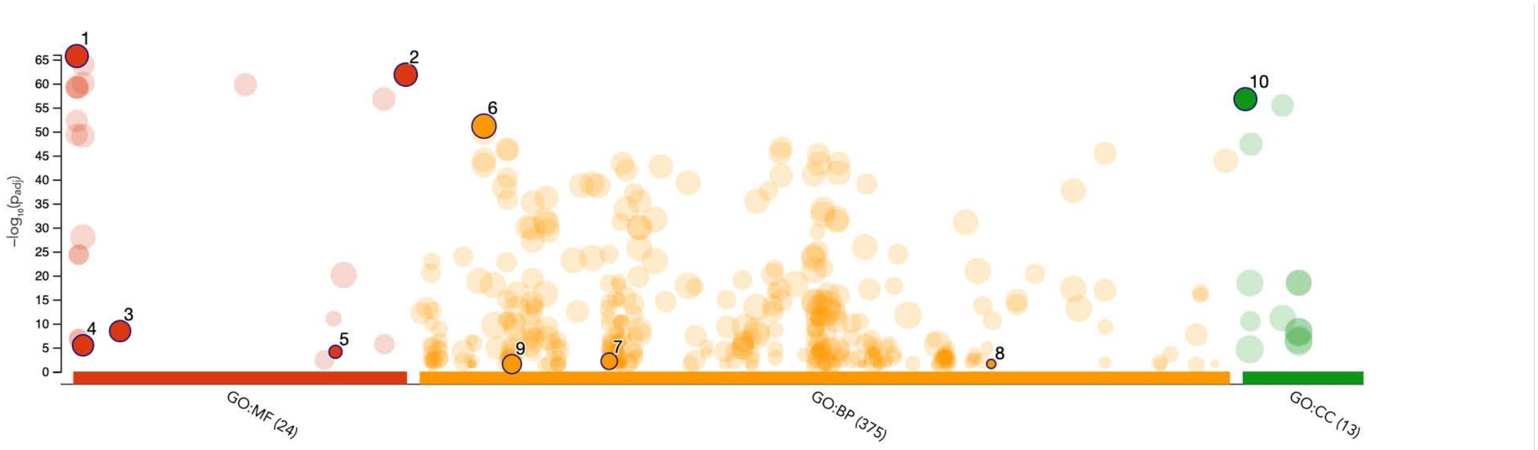


| ID | Source | Term ID | Term Name | $p_{adj}$ (query_1) |
|---|---|---|---|---|
| 1 | GO:MF | GO:0000981 | DNA-binding transcription factor activity, RNA polymerase II-specific | $1.896{\times}10^{-66}$ |
| 2 | GO:MF | GO:1990837 | sequence-specific double-stranded DNA binding | $1.439{\times}10^{-62}$ |
| 3 | GO:MF | GO:0008134 | transcription factor binding | $3.743{\times}10^{-9}$ |
| 4 | GO:MF | GO:0003682 | chromatin binding | $3.416{\times}10^{-6}$ |
| 5 | GO:MF | GO:0071837 | HMG box domain binding | $7.655{\times}10^{-5}$ |
| 6 | GO:BP | GO:0006357 | regulation of transcription by RNA polymerase II | $7.619{\times}10^{-52}$ |
| 7 | GO:BP | GO:0021545 | cranial nerve development | $6.963{\times}10^{-3}$ |
| 8 | GO:BP | GO:0097378 | dorsal spinal cord interneuron axon guidance | $2.578{\times}10^{-2}$ |
| 9 | GO:BP | GO:0007601 | visual perception | $2.868{\times}10^{-2}$ |
| 10 | GO:CC | GO:0000785 | chromatin | $1.833{\times}10^{-57}$ |

**Cluster 12**

| | |
|---|---|
| **version** | e111_eg58_p18_f463989d |
| **date** | 8/26/2024, 6:33:49 PM |
| **organism** | hsapiens |

g:Profiler

**Figure S7j**

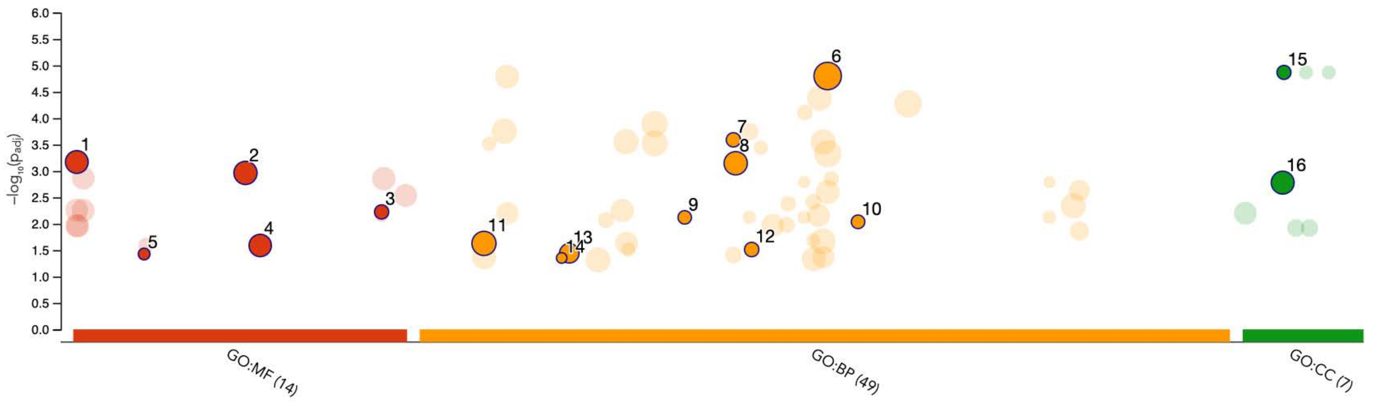


| ID | Source | Term ID | Term Name | $p_{adj}$ (query_1) |
|---|---|---|---|---|
| 1 | GO:MF | GO:0000981 | DNA-binding transcription factor activity, RNA polymerase II-specific | $6.750\times10^{-4}$ |
| 2 | GO:MF | GO:0043565 | sequence-specific DNA binding | $1.086\times10^{-3}$ |
| 3 | GO:MF | GO:0120020 | cholesterol transfer activity | $5.955\times10^{-3}$ |
| 4 | GO:MF | GO:0046983 | protein dimerization activity | $2.574\times10^{-2}$ |
| 5 | GO:MF | GO:0015168 | glycerol transmembrane transporter activity | $3.740\times10^{-2}$ |
| 6 | GO:BP | GO:0050789 | regulation of biological process | $1.585\times10^{-5}$ |
| 7 | GO:BP | GO:0042438 | melanin biosynthetic process | $2.568\times10^{-4}$ |
| 8 | GO:BP | GO:0042592 | homeostatic process | $7.129\times10^{-4}$ |
| 9 | GO:BP | GO:0034375 | high-density lipoprotein particle remodeling | $7.568\times10^{-3}$ |
| 10 | GO:BP | GO:0055091 | phospholipid homeostasis | $9.177\times10^{-3}$ |
| 11 | GO:BP | GO:0006357 | regulation of transcription by RNA polymerase II | $2.354\times10^{-2}$ |
| 12 | GO:BP | GO:0043576 | regulation of respiratory gaseous exchange | $3.070\times10^{-2}$ |
| 13 | GO:BP | GO:0015850 | organic hydroxy compound transport | $3.618\times10^{-2}$ |
| 14 | GO:BP | GO:0014826 | vein smooth muscle contraction | $4.439\times10^{-2}$ |
| 15 | GO:CC | GO:0033162 | melanosome membrane | $1.356\times10^{-5}$ |
| 16 | GO:CC | GO:0032993 | protein-DNA complex | $1.661\times10^{-3}$ |

**Cluster 13**

**version** e111_eg58_p18_f463989d
**date** 8/26/2024, 6:34:23 PM
**organism** hsapiens

g:Profiler

**Figure S7k**

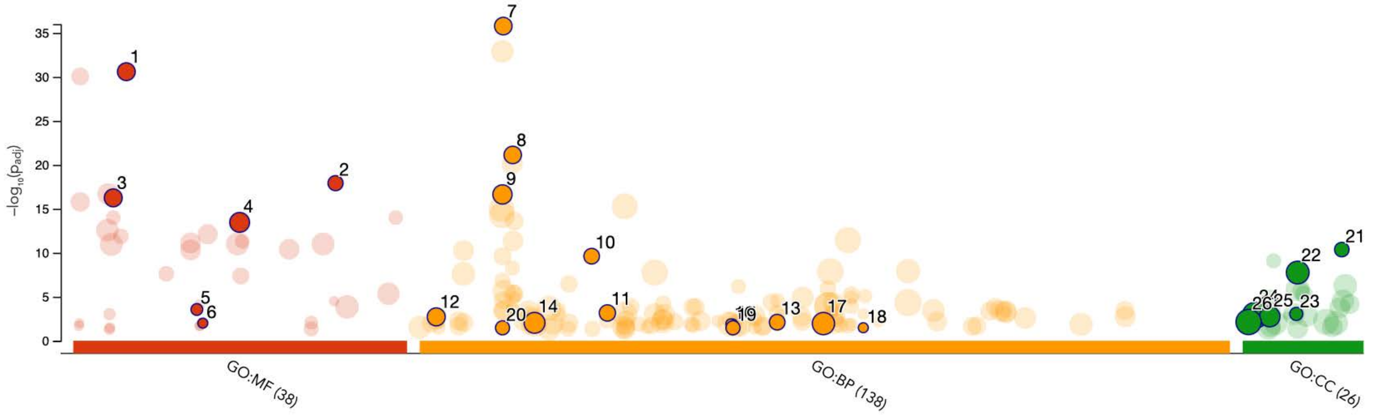


| ID | Source | Term ID | Term Name | $p_{adj}$ (query_1) |
|---|---|---|---|---|
| 1 | GO:MF | GO:0008528 | G protein-coupled peptide receptor activity | $2.569\times10^{-31}$ |
| 2 | GO:MF | GO:0071855 | neuropeptide receptor binding | $1.210\times10^{-18}$ |
| 3 | GO:MF | GO:0005179 | hormone activity | $5.682\times10^{-17}$ |
| 4 | GO:MF | GO:0042277 | peptide binding | $3.494\times10^{-14}$ |
| 5 | GO:MF | GO:0031628 | opioid receptor binding | $2.798\times10^{-4}$ |
| 6 | GO:MF | GO:0031893 | vasopressin receptor binding | $9.945\times10^{-3}$ |
| 7 | GO:BP | GO:0007218 | neuropeptide signaling pathway | $1.666\times10^{-36}$ |
| 8 | GO:BP | GO:0007631 | feeding behavior | $7.475\times10^{-22}$ |
| 9 | GO:BP | GO:0007188 | adenylate cyclase-modulating G protein-coupled receptor signaling pathway | $2.323\times10^{-17}$ |
| 10 | GO:BP | GO:0019098 | reproductive behavior | $2.437\times10^{-10}$ |
| 11 | GO:BP | GO:0019933 | cAMP-mediated signaling | $6.923\times10^{-4}$ |
| 12 | GO:BP | GO:0001936 | regulation of endothelial cell proliferation | $1.933\times10^{-3}$ |
| 13 | GO:BP | GO:0045761 | regulation of adenylate cyclase activity | $7.763\times10^{-3}$ |
| 14 | GO:BP | GO:0009991 | response to extracellular stimulus | $9.213\times10^{-3}$ |
| 15 | GO:BP | GO:0042418 | epinephrine biosynthetic process | $1.110\times10^{-2}$ |
| 16 | GO:BP | GO:0042309 | homoiothermy | $1.110\times10^{-2}$ |
| 17 | GO:BP | GO:0048878 | chemical homeostasis | $1.120\times10^{-2}$ |
| 18 | GO:BP | GO:0060183 | apelin receptor signaling pathway | $3.326\times10^{-2}$ |
| 19 | GO:BP | GO:0042423 | catecholamine biosynthetic process | $3.403\times10^{-2}$ |
| 20 | GO:BP | GO:0007190 | activation of adenylate cyclase activity | $3.403\times10^{-2}$ |
| 21 | GO:CC | GO:0098992 | neuronal dense core vesicle | $4.300\times10^{-11}$ |
| 22 | GO:CC | GO:0043005 | neuron projection | $1.754\times10^{-8}$ |
| 23 | GO:CC | GO:0042583 | chromaffin granule | $8.911\times10^{-4}$ |
| 24 | GO:CC | GO:0005886 | plasma membrane | $1.260\times10^{-3}$ |
| 25 | GO:CC | GO:0030133 | transport vesicle | $1.826\times10^{-3}$ |
| 26 | GO:CC | GO:0005576 | extracellular region | $7.429\times10^{-3}$ |

**Cluster 14**

| | |
|---|---|
| **version** | e111_eg58_p18_f463989d |
| **date** | 8/26/2024, 6:34:55 PM |
| **organism** | hsapiens |

g:Profiler

**Figure S7I**

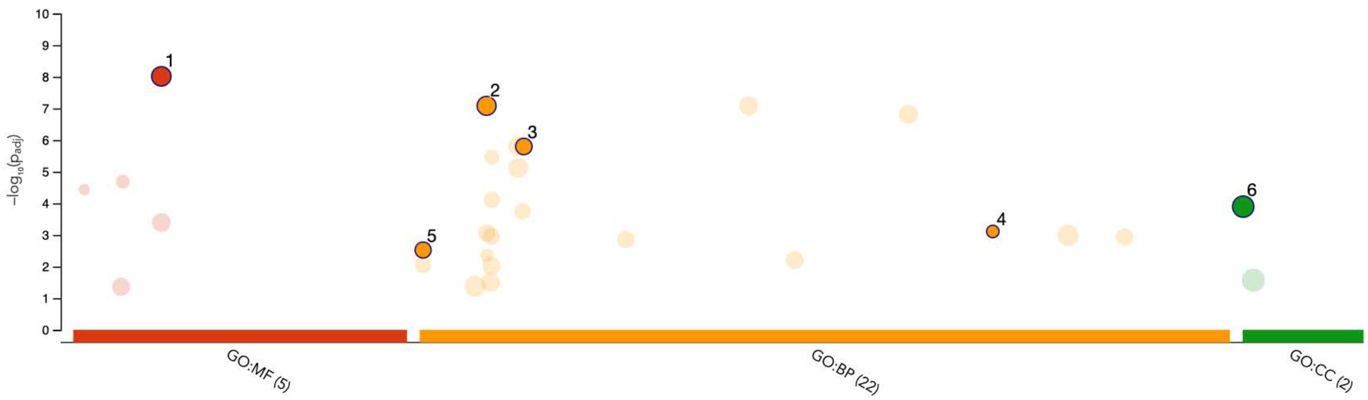


| ID | Source | Term ID | Term Name | $p_{adj}$ (query_1) |
|---|---|---|---|---|
| 1 | GO:MF | GO:0016757 | glycosyltransferase activity | $9.660\times10^{-9}$ |
| 2 | GO:BP | GO:0006486 | protein glycosylation | $8.254\times10^{-8}$ |
| 3 | GO:BP | GO:0009311 | oligosaccharide metabolic process | $1.594\times10^{-6}$ |
| 4 | GO:BP | GO:0097503 | sialylation | $7.763\times10^{-4}$ |
| 5 | GO:BP | GO:0000381 | regulation of alternative mRNA splicing, via spliceosome | $2.998\times10^{-3}$ |
| 6 | GO:CC | GO:0000139 | Golgi membrane | $1.274\times10^{-4}$ |

**Cluster 16**

**version** e111_eg58_p18_f463989d
**date** 8/26/2024, 6:36:06 PM
**organism** hsapiens

g:Profiler

**Figure S7m**

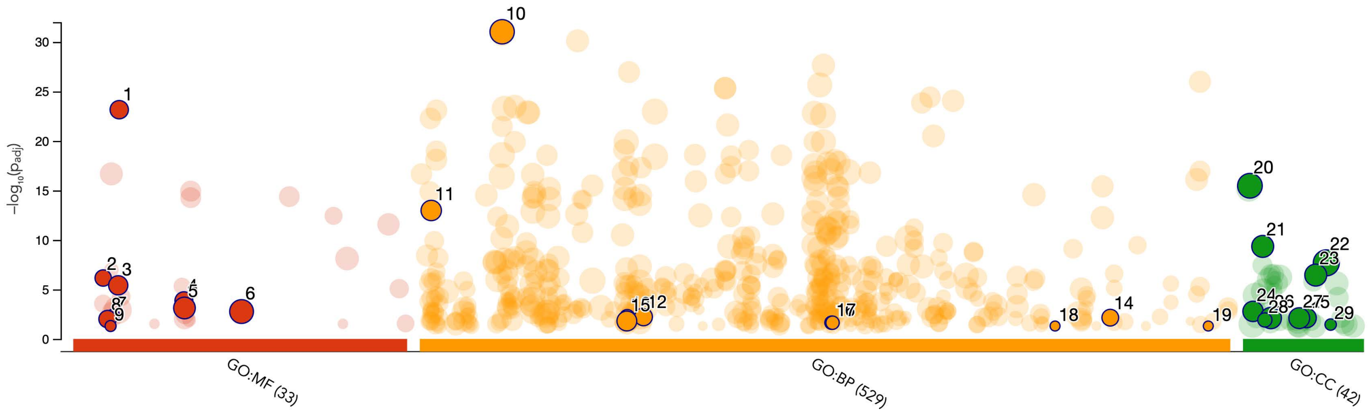


| ID | Source | Term ID | Term Name | $p_{adj}$ (query_... |
|---|---|---|---|---|
| 1 | GO:MF | GO:0008083 | growth factor activity | $6.958\times10^{-24}$ |
| 2 | GO:MF | GO:0004714 | transmembrane receptor protein tyrosine kinase activity | $7.050\times10^{-7}$ |
| 3 | GO:MF | GO:0005539 | glycosaminoglycan binding | $3.860\times10^{-6}$ |
| 4 | GO:MF | GO:0019838 | growth factor binding | $1.390\times10^{-4}$ |
| 5 | GO:MF | GO:0019900 | kinase binding | $7.505\times10^{-4}$ |
| 6 | GO:MF | GO:0042802 | identical protein binding | $1.715\times10^{-3}$ |
| 7 | GO:MF | GO:0005172 | vascular endothelial growth factor receptor binding | $4.502\times10^{-3}$ |
| 8 | GO:MF | GO:0004896 | cytokine receptor activity | $9.862\times10^{-3}$ |
| 9 | GO:MF | GO:0005030 | neurotrophin receptor activity | $4.979\times10^{-2}$ |
| 10 | GO:BP | GO:0007166 | cell surface receptor signaling pathway | $9.377\times10^{-32}$ |
| 11 | GO:BP | GO:0001667 | ameboidal-type cell migration | $1.062\times10^{-13}$ |
| 12 | GO:BP | GO:0031623 | receptor internalization | $5.856\times10^{-3}$ |
| 13 | GO:BP | GO:0030212 | hyaluronan metabolic process | $6.146\times10^{-3}$ |
| 14 | GO:BP | GO:1902893 | regulation of miRNA transcription | $7.110\times10^{-3}$ |
| 15 | GO:BP | GO:0030198 | extracellular matrix organization | $1.554\times10^{-2}$ |
| 16 | GO:BP | GO:0050930 | induction of positive chemotaxis | $2.307\times10^{-2}$ |
| 17 | GO:BP | GO:0050966 | detection of mechanical stimulus involved in sensory perception of pain | $2.307\times10^{-2}$ |
| 18 | GO:BP | GO:1900625 | positive regulation of monocyte aggregation | $4.974\times10^{-2}$ |
| 19 | GO:BP | GO:2000446 | regulation of macrophage migration inhibitory factor signaling pathway | $4.974\times10^{-2}$ |
| 20 | GO:CC | GO:0005615 | extracellular space | $3.354\times10^{-16}$ |
| 21 | GO:CC | GO:0009986 | cell surface | $4.504\times10^{-10}$ |
| 22 | GO:CC | GO:0071944 | cell periphery | $2.080\times10^{-8}$ |
| 23 | GO:CC | GO:0070161 | anchoring junction | $3.567\times10^{-7}$ |
| 24 | GO:CC | GO:0005769 | early endosome | $1.653\times10^{-3}$ |
| 25 | GO:CC | GO:0045121 | membrane raft | $7.652\times10^{-3}$ |
| 26 | GO:CC | GO:0030424 | axon | $7.840\times10^{-3}$ |
| 27 | GO:CC | GO:0043235 | receptor complex | $7.951\times10^{-3}$ |
| 28 | GO:CC | GO:0016327 | apicolateral plasma membrane | $1.211\times10^{-2}$ |
| 29 | GO:CC | GO:0097180 | serine protease inhibitor complex | $3.560\times10^{-2}$ |

**Cluster 17**

| | | |
|---|---|---|
| **version** | e111_eg58_p18_f463989d | g:Profiler |
| **date** | 9/6/2024, 6:37:45 PM | |
| **organism** | hsapiens | |

**Figure S7n**

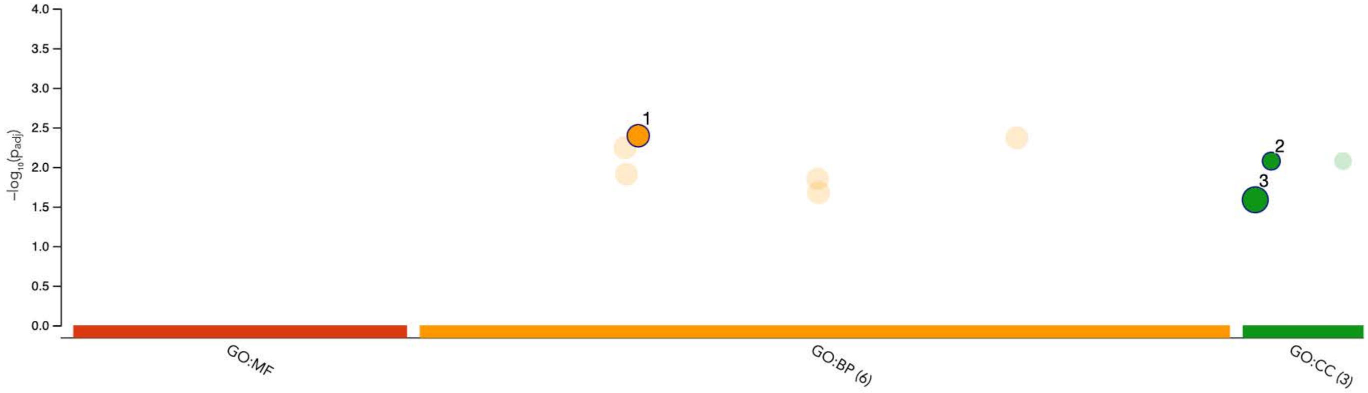


| ID | Source | Term ID | Term Name | $p_{adj}$ (query_1) |
|---|---|---|---|---|
| 1 | GO:BP | GO:0031175 | neuron projection development | $4.034\times10^{-3}$ |
| 2 | GO:CC | GO:0030672 | synaptic vesicle membrane | $8.425\times10^{-3}$ |
| 3 | GO:CC | GO:0005886 | plasma membrane | $2.600\times10^{-2}$ |

**Cluster 18**

| | |
|---|---|
| **version** | e111_eg58_p18_f463989d |
| **date** | 8/26/2024, 6:36:40 PM |
| **organism** | hsapiens |

g:Profiler

**Figure S7o**

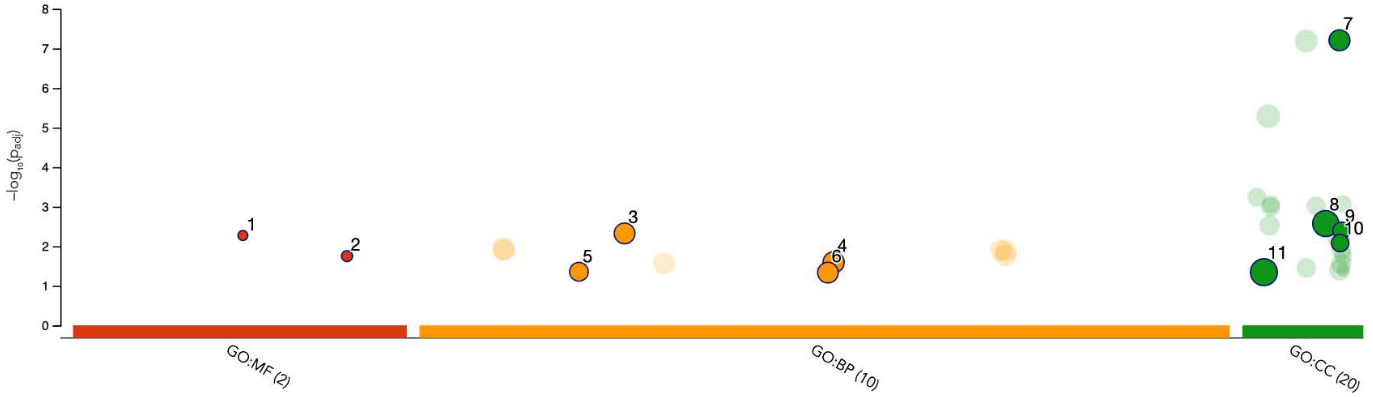


| ID | Source | Term ID | Term Name | $p_{adj}$ (query_1) |
|---|---|---|---|---|
| 1 | GO:MF | GO:0042988 | X11-like protein binding | $5.351\times10^{-3}$ |
| 2 | GO:MF | GO:0098882 | structural constituent of presynaptic active zone | $1.779\times10^{-2}$ |
| 3 | GO:BP | GO:0023061 | signal release | $4.751\times10^{-3}$ |
| 4 | GO:BP | GO:0051046 | regulation of secretion | $2.549\times10^{-2}$ |
| 5 | GO:BP | GO:0017157 | regulation of exocytosis | $4.417\times10^{-2}$ |
| 6 | GO:BP | GO:0050808 | synapse organization | $4.648\times10^{-2}$ |
| 7 | GO:CC | GO:0098793 | presynapse | $6.223\times10^{-8}$ |
| 8 | GO:CC | GO:0071944 | cell periphery | $2.674\times10^{-3}$ |
| 9 | GO:CC | GO:0098982 | GABA-ergic synapse | $4.017\times10^{-3}$ |
| 10 | GO:CC | GO:0098839 | postsynaptic density membrane | $8.309\times10^{-3}$ |
| 11 | GO:CC | GO:0016020 | membrane | $4.537\times10^{-2}$ |

**Cluster 19**

**version** e111_eg58_p18_f463989d
**date** 8/26/2024, 6:37:06 PM
**organism** hsapiens

g:Profiler

**Figure S7p**

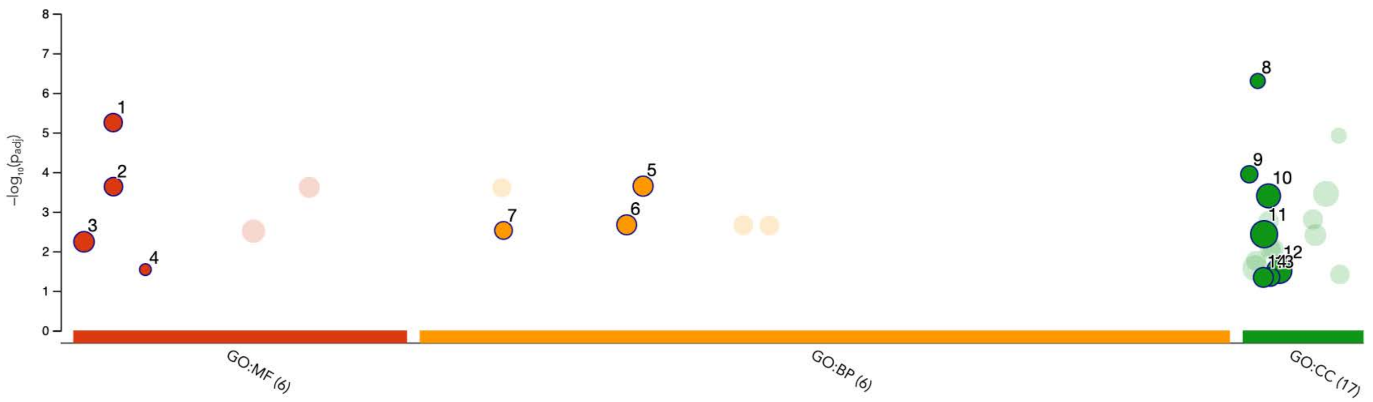


| ID | Source | Term ID | Term Name | $p_{adj}$ (query_1) |
|---|---|---|---|---|
| 1 | GO:MF | GO:0005178 | integrin binding | $5.640\times10^{-6}$ |
| 2 | GO:MF | GO:0005201 | extracellular matrix structural constituent | $2.337\times10^{-4}$ |
| 3 | GO:MF | GO:0003779 | actin binding | $5.773\times10^{-3}$ |
| 4 | GO:MF | GO:0015220 | choline transmembrane transporter activity | $2.909\times10^{-2}$ |
| 5 | GO:BP | GO:0031589 | cell-substrate adhesion | $2.270\times10^{-4}$ |
| 6 | GO:BP | GO:0030198 | extracellular matrix organization | $2.156\times10^{-3}$ |
| 7 | GO:BP | GO:0007229 | integrin-mediated signaling pathway | $2.982\times10^{-3}$ |
| 8 | GO:CC | GO:0008305 | integrin complex | $5.016\times10^{-7}$ |
| 9 | GO:CC | GO:0005604 | basement membrane | $1.137\times10^{-4}$ |
| 10 | GO:CC | GO:0030054 | cell junction | $3.993\times10^{-4}$ |
| 11 | GO:CC | GO:0016020 | membrane | $3.713\times10^{-3}$ |
| 12 | GO:CC | GO:0031982 | vesicle | $3.132\times10^{-2}$ |
| 13 | GO:CC | GO:0030426 | growth cone | $4.481\times10^{-2}$ |
| 14 | GO:CC | GO:0014069 | postsynaptic density | $4.580\times10^{-2}$ |

**Cluster 20**

| | |
|---|---|
| **version** | e111_eg58_p18_f463989d |
| **date** | 8/26/2024, 6:37:35 PM |
| **organism** | hsapiens |

g:Profiler

**Figure S7q**

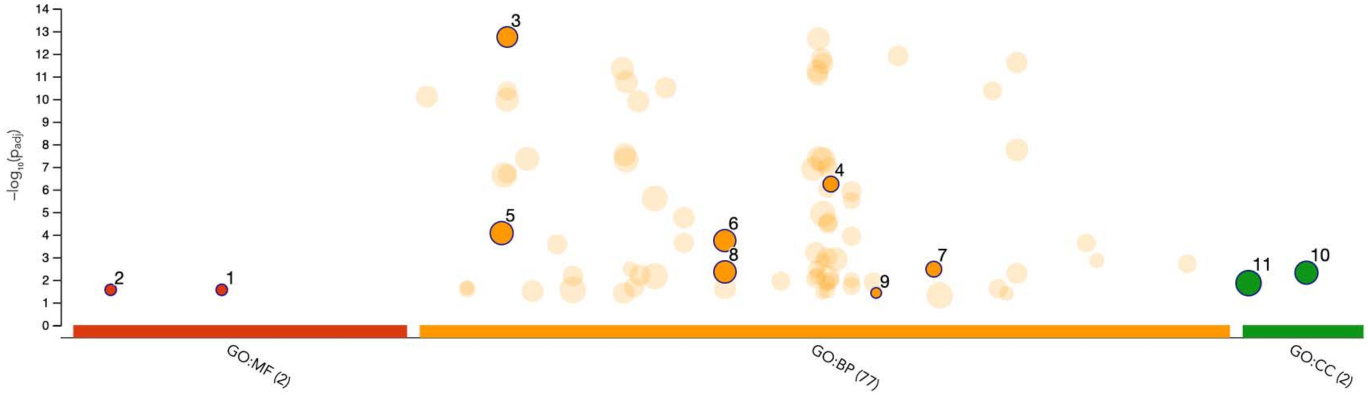


| ID | Source | Term ID | Term Name | $p_{adj}$ (query_1) |
|---|---|---|---|---|
| 1 | GO:MF | GO:0034481 | chondroitin sulfotransferase activity | $2.748 \times 10^{-2}$ |
| 2 | GO:MF | GO:0005042 | netrin receptor activity | $2.748 \times 10^{-2}$ |
| 3 | GO:BP | GO:0007409 | axonogenesis | $1.802 \times 10^{-13}$ |
| 4 | GO:BP | GO:0050919 | negative chemotaxis | $5.790 \times 10^{-7}$ |
| 5 | GO:BP | GO:0007155 | cell adhesion | $8.523 \times 10^{-5}$ |
| 6 | GO:BP | GO:0040007 | growth | $1.834 \times 10^{-4}$ |
| 7 | GO:BP | GO:0071526 | semaphorin-plexin signaling pathway | $3.368 \times 10^{-3}$ |
| 8 | GO:BP | GO:0040012 | regulation of locomotion | $4.425 \times 10^{-3}$ |
| 9 | GO:BP | GO:0060666 | dichotomous subdivision of terminal units involved in salivary gland branching | $3.770 \times 10^{-2}$ |
| 10 | GO:CC | GO:0045202 | synapse | $4.807 \times 10^{-3}$ |
| 11 | GO:CC | GO:0005576 | extracellular region | $1.394 \times 10^{-2}$ |

**Cluster 21**

**version** e111_eg58_p18_f463989d
**date** 8/26/2024, 6:38:02 PM
**organism** hsapiens

g:Profiler

**Figure S7r**

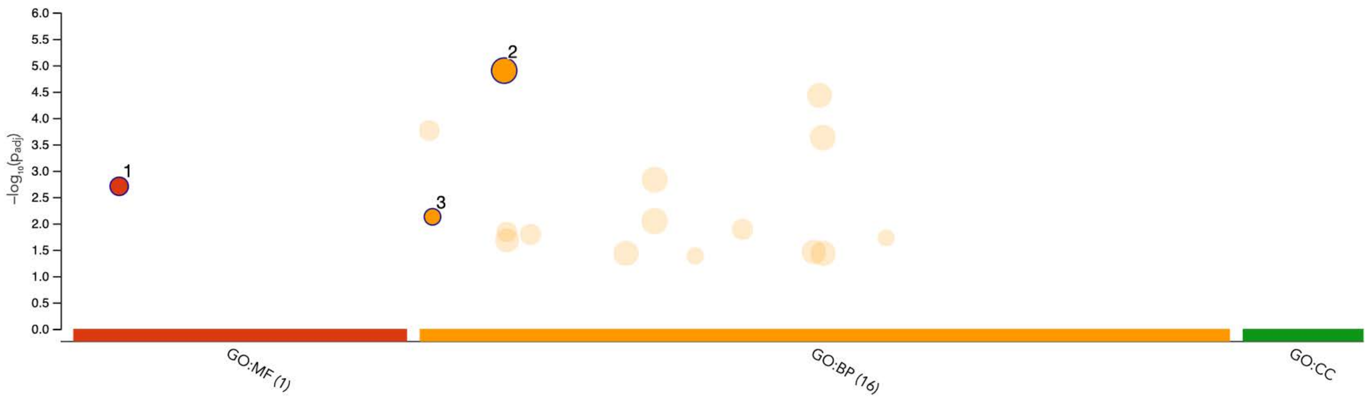


| ID | Source | Term ID | Term Name | $p_{adj}$ (query_1) |
|---|---|---|---|---|
| 1 | GO:MF | GO:0008083 | growth factor activity | $1.976\times10^{-3}$ |
| 2 | GO:BP | GO:0007275 | multicellular organism development | $1.261\times10^{-5}$ |
| 3 | GO:BP | GO:0001756 | somitogenesis | $7.482\times10^{-3}$ |

**Cluster 22**

| | |
|---|---|
| **version** | e111_eg58_p18_f463989d |
| **date** | 8/26/2024, 6:38:35 PM |
| **organism** | hsapiens |

g:Profiler

**Figure S7s**

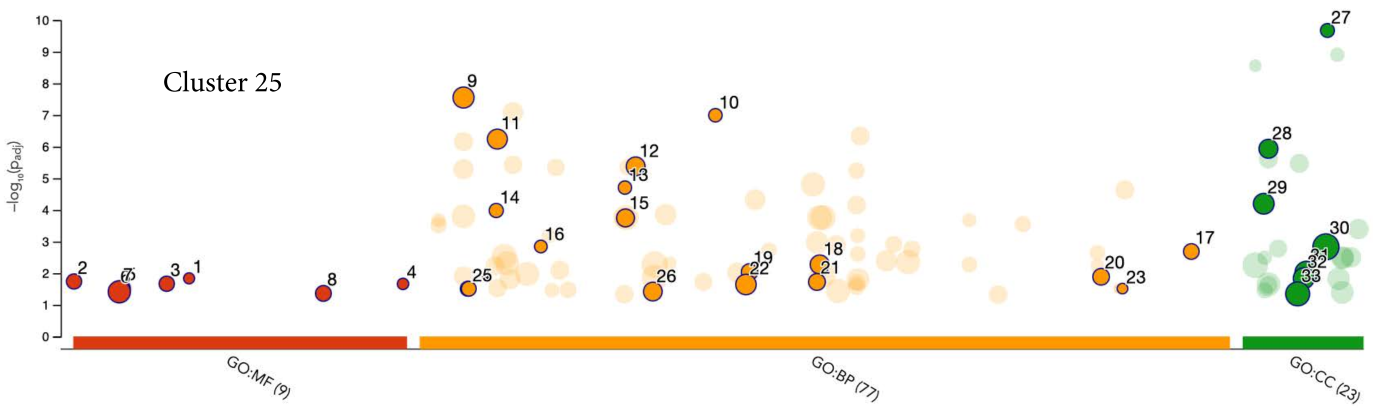


| ID | Source | Term ID | Term Name | $p_{adj}$ (query_1) |
|---|---|---|---|---|
| 1 | GO:MF | GO:0030298 | receptor signaling protein tyrosine kinase activator activity | $1.457\times10^{-2}$ |
| 2 | GO:MF | GO:0000146 | microfilament motor activity | $1.830\times10^{-2}$ |
| 3 | GO:MF | GO:0017080 | sodium channel regulator activity | $2.156\times10^{-2}$ |
| 4 | GO:MF | GO:1990239 | steroid hormone binding | $2.184\times10^{-2}$ |
| 5 | GO:MF | GO:0008307 | structural constituent of muscle | $2.918\times10^{-2}$ |
| 6 | GO:MF | GO:0005391 | P-type sodium:potassium-exchanging transporter activity | $3.053\times10^{-2}$ |
| 7 | GO:MF | GO:0008092 | cytoskeletal protein binding | $3.878\times10^{-2}$ |
| 8 | GO:MF | GO:0060590 | ATPase regulator activity | $4.362\times10^{-2}$ |
| 9 | GO:BP | GO:0003013 | circulatory system process | $2.800\times10^{-8}$ |
| 10 | GO:BP | GO:0036376 | sodium ion export across plasma membrane | $1.017\times10^{-7}$ |
| 11 | GO:BP | GO:0006936 | muscle contraction | $5.774\times10^{-7}$ |
| 12 | GO:BP | GO:0031032 | actomyosin structure organization | $4.198\times10^{-6}$ |
| 13 | GO:BP | GO:0030007 | intracellular potassium ion homeostasis | $1.966\times10^{-5}$ |
| 14 | GO:BP | GO:0006883 | intracellular sodium ion homeostasis | $1.044\times10^{-4}$ |
| 15 | GO:BP | GO:0030048 | actin filament-based movement | $1.796\times10^{-4}$ |
| 16 | GO:BP | GO:0010248 | establishment or maintenance of transmembrane electrochemical gradient | $1.434\times10^{-3}$ |
| 17 | GO:BP | GO:1990573 | potassium ion import across plasma membrane | $2.064\times10^{-3}$ |
| 18 | GO:BP | GO:0048738 | cardiac muscle tissue development | $5.326\times10^{-3}$ |
| 19 | GO:BP | GO:0043462 | regulation of ATP-dependent activity | $9.141\times10^{-3}$ |
| 20 | GO:BP | GO:1902475 | L-alpha-amino acid transmembrane transport | $1.302\times10^{-2}$ |
| 21 | GO:BP | GO:0048644 | muscle organ morphogenesis | $1.894\times10^{-2}$ |
| 22 | GO:BP | GO:0043269 | regulation of monoatomic ion transport | $2.280\times10^{-2}$ |
| 23 | GO:BP | GO:1903416 | response to glycoside | $3.069\times10^{-2}$ |
| 24 | GO:BP | GO:0003171 | atrioventricular valve development | $3.105\times10^{-2}$ |
| 25 | GO:BP | GO:0003209 | cardiac atrium morphogenesis | $3.105\times10^{-2}$ |
| 26 | GO:BP | GO:0032412 | regulation of monoatomic ion transmembrane transporter activity | $3.816\times10^{-2}$ |
| 27 | GO:CC | GO:0090533 | cation-transporting ATPase complex | $2.122\times10^{-10}$ |
| 28 | GO:CC | GO:0030017 | sarcomere | $1.167\times10^{-6}$ |
| 29 | GO:CC | GO:0015629 | actin cytoskeleton | $6.385\times10^{-5}$ |
| 30 | GO:CC | GO:0071944 | cell periphery | $1.472\times10^{-3}$ |
| 31 | GO:CC | GO:0045202 | synapse | $9.662\times10^{-3}$ |
| 32 | GO:CC | GO:0044297 | cell body | $1.448\times10^{-2}$ |
| 33 | GO:CC | GO:0042995 | cell projection | $4.515\times10^{-2}$ |

**Custer 24**

| | |
|---|---|
| **version** | e111_eg58_p18_f463989d |
| **date** | 8/26/2024, 6:39:30 PM |
| **organism** | hsapiens |

g:Profiler

**Figure S7t**

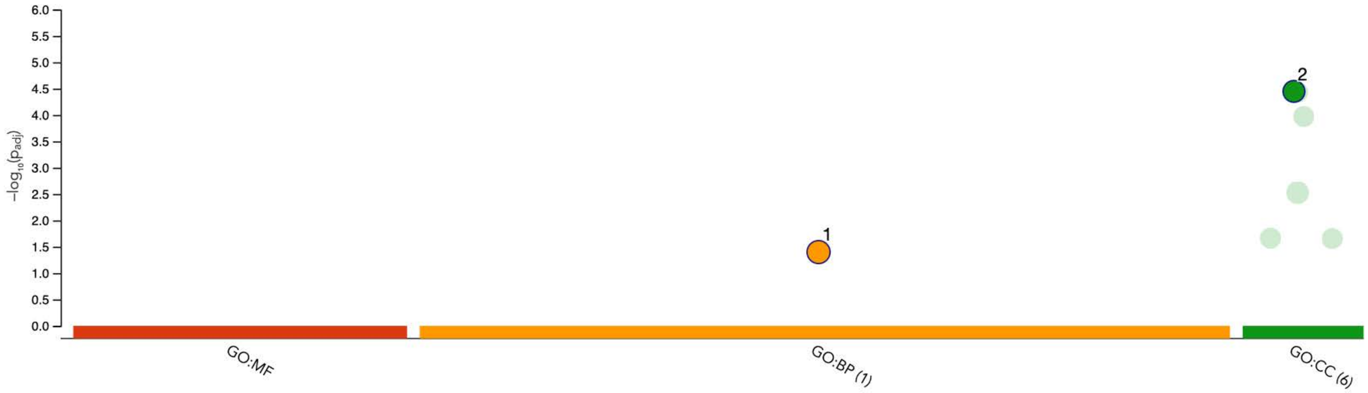


| ID | Source | Term ID | Term Name | $p_{adj}$ (query_1) |
|---|---|---|---|---|
| 1 | GO:BP | GO:0048699 | generation of neurons | $3.987 \times 10^{-2}$ |
| 2 | GO:CC | GO:0036477 | somatodendritic compartment | $3.562 \times 10^{-5}$ |

**Cluster 25**

| | | |
|---|---|---|
| **version** | e111_eg58_p18_f463989d | g:Profiler |
| **date** | 8/26/2024, 6:39:55 PM | |
| **organism** | hsapiens | |

**Figure S7u**

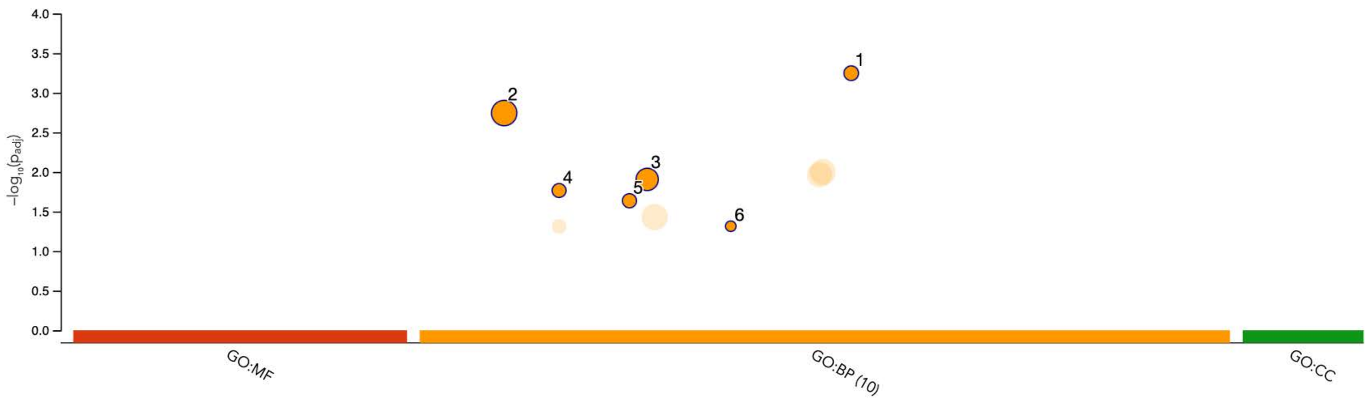


| ID | Source | Term ID | Term Name | $p_{adj}$ (query_1) |
|---|---|---|---|---|
| 1 | GO:BP | GO:0051953 | negative regulation of amine transport | $5.659\times10^{-4}$ |
| 2 | GO:BP | GO:0007275 | multicellular organism development | $1.803\times10^{-3}$ |
| 3 | GO:BP | GO:0032101 | regulation of response to external stimulus | $1.243\times10^{-2}$ |
| 4 | GO:BP | GO:0014048 | regulation of glutamate secretion | $1.718\times10^{-2}$ |
| 5 | GO:BP | GO:0030431 | sleep | $2.314\times10^{-2}$ |
| 6 | GO:BP | GO:0042321 | negative regulation of circadian sleep/wake cycle, sleep | $4.854\times10^{-2}$ |

**Cluster 1**

| | |
|---|---|
| **version** | e111_eg58_p18_f463989d |
| **date** | 8/26/2024, 6:40:31 PM |
| **organism** | hsapiens |

g:Profiler

**Figure S8a**

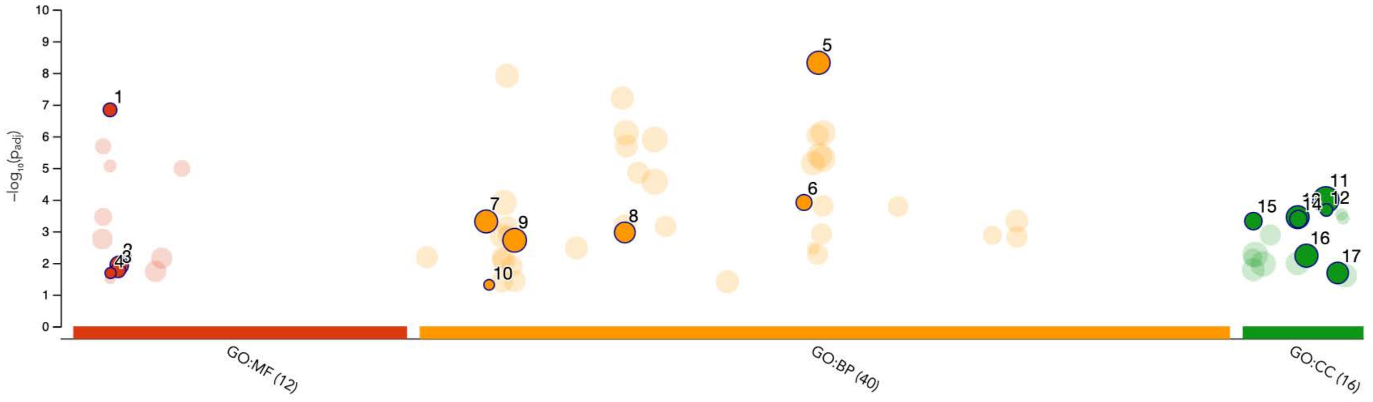


| ID | Source | Term ID | Term Name | $p_{adj}$ (query_1) |
|---|---|---|---|---|
| 1 | GO:MF | GO:0005003 | ephrin receptor activity | $1.461 \times 10^{-7}$ |
| 2 | GO:MF | GO:0008083 | growth factor activity | $1.182 \times 10^{-2}$ |
| 3 | GO:MF | GO:0005540 | hyaluronic acid binding | $1.689 \times 10^{-2}$ |
| 4 | GO:MF | GO:0005030 | neurotrophin receptor activity | $2.078 \times 10^{-2}$ |
| 5 | GO:BP | GO:0048699 | generation of neurons | $4.771 \times 10^{-9}$ |
| 6 | GO:BP | GO:0048013 | ephrin receptor signaling pathway | $1.231 \times 10^{-4}$ |
| 7 | GO:BP | GO:0006468 | protein phosphorylation | $4.888 \times 10^{-4}$ |
| 8 | GO:BP | GO:0023061 | signal release | $1.085 \times 10^{-3}$ |
| 9 | GO:BP | GO:0008283 | cell population proliferation | $1.918 \times 10^{-3}$ |
| 10 | GO:BP | GO:0006583 | melanin biosynthetic process from tyrosine | $4.887 \times 10^{-2}$ |
| 11 | GO:CC | GO:0071944 | cell periphery | $1.004 \times 10^{-4}$ |
| 12 | GO:CC | GO:0072534 | perineuronal net | $2.121 \times 10^{-4}$ |
| 13 | GO:CC | GO:0043005 | neuron projection | $3.582 \times 10^{-4}$ |
| 14 | GO:CC | GO:0043202 | lysosomal lumen | $4.109 \times 10^{-4}$ |
| 15 | GO:CC | GO:0005796 | Golgi lumen | $4.756 \times 10^{-4}$ |
| 16 | GO:CC | GO:0045202 | synapse | $5.883 \times 10^{-3}$ |
| 17 | GO:CC | GO:0098552 | side of membrane | $2.069 \times 10^{-2}$ |

**Cluster 2**

| | |
|---|---|
| **version** | e111_eg58_p18_f463989d |
| **date** | 8/26/2024, 6:40:59 PM |
| **organism** | hsapiens |

g:Profiler

**Figure S8b**

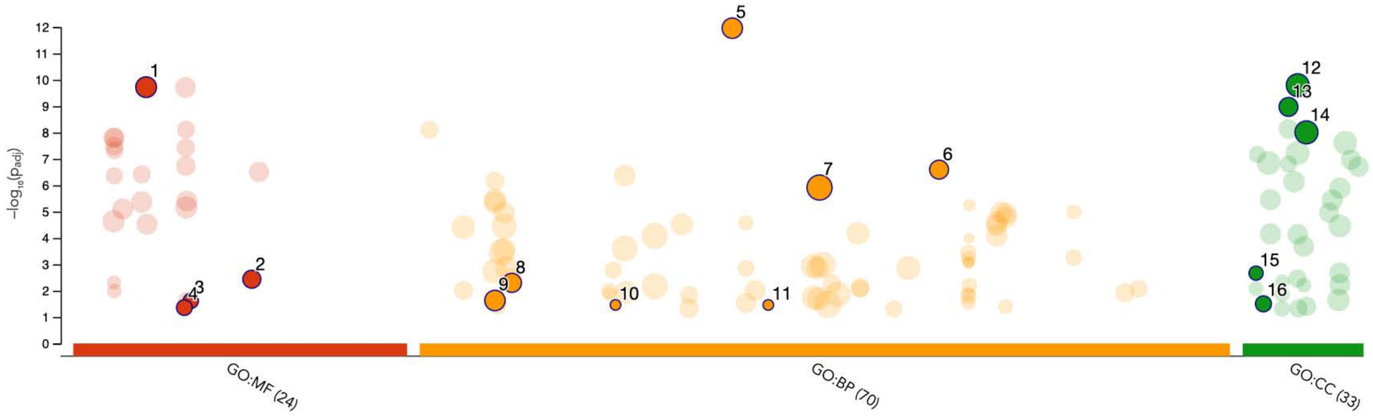


| ID | Source | Term ID | Term Name | $p_{adj}$ (query_1) |
|---|---|---|---|---|
| 1 | GO:MF | GO:0015267 | channel activity | $1.897\times10^{-10}$ |
| 2 | GO:MF | GO:0044325 | transmembrane transporter binding | $3.652\times10^{-3}$ |
| 3 | GO:MF | GO:0030551 | cyclic nucleotide binding | $2.420\times10^{-2}$ |
| 4 | GO:MF | GO:0019894 | kinesin binding | $4.372\times10^{-2}$ |
| 5 | GO:BP | GO:0042391 | regulation of membrane potential | $1.099\times10^{-12}$ |
| 6 | GO:BP | GO:0071805 | potassium ion transmembrane transport | $2.549\times10^{-7}$ |
| 7 | GO:BP | GO:0048731 | system development | $1.218\times10^{-6}$ |
| 8 | GO:BP | GO:0007601 | visual perception | $5.026\times10^{-3}$ |
| 9 | GO:BP | GO:0006816 | calcium ion transport | $2.344\times10^{-2}$ |
| 10 | GO:BP | GO:0021759 | globus pallidus development | $3.458\times10^{-2}$ |
| 11 | GO:BP | GO:0045163 | clustering of voltage-gated potassium channels | $3.458\times10^{-2}$ |
| 12 | GO:CC | GO:0043005 | neuron projection | $1.595\times10^{-10}$ |
| 13 | GO:CC | GO:0034703 | cation channel complex | $1.063\times10^{-9}$ |
| 14 | GO:CC | GO:0045202 | synapse | $9.808\times10^{-9}$ |
| 15 | GO:CC | GO:0005922 | connexin complex | $2.150\times10^{-3}$ |
| 16 | GO:CC | GO:0014704 | intercalated disc | $3.139\times10^{-2}$ |

**Cluster 3**

| | |
|---|---|
| **version** | e111_eg58_p18_f463989d |
| **date** | 8/26/2024, 6:41:29 PM |
| **organism** | hsapiens |

g:Profiler

**Figure S8c**

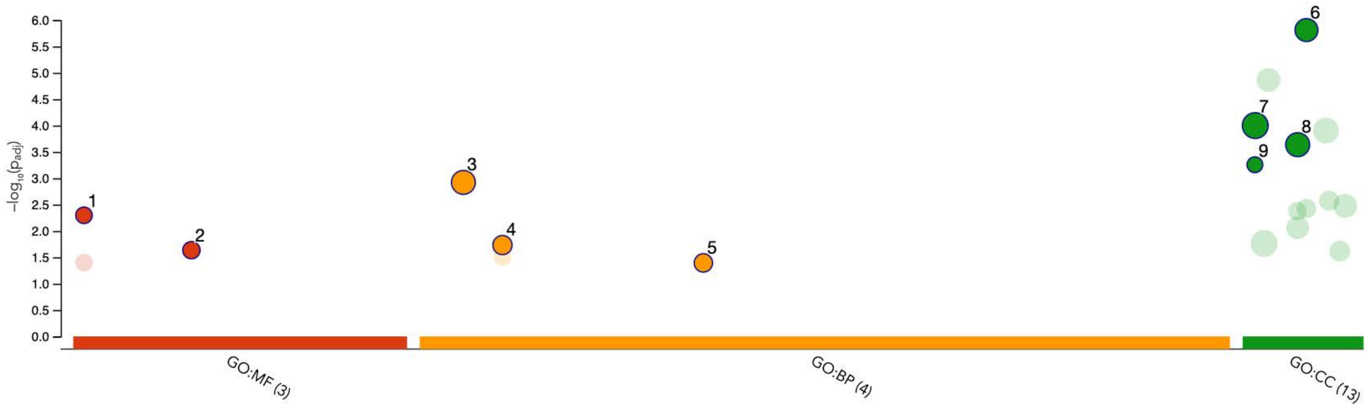


| ID | Source | Term ID | Term Name | $p_{adj}$ (query_1) |
|---|---|---|---|---|
| 1 | GO:MF | GO:0003777 | microtubule motor activity | $5.046 \times 10^{-3}$ |
| 2 | GO:MF | GO:0030594 | neurotransmitter receptor activity | $2.316 \times 10^{-2}$ |
| 3 | GO:BP | GO:0003008 | system process | $1.197 \times 10^{-3}$ |
| 4 | GO:BP | GO:0007188 | adenylate cyclase-modulating G protein-coupled receptor signaling pathway | $1.851 \times 10^{-2}$ |
| 5 | GO:BP | GO:0035725 | sodium ion transmembrane transport | $4.036 \times 10^{-2}$ |
| 6 | GO:CC | GO:0045202 | synapse | $1.544 \times 10^{-6}$ |
| 7 | GO:CC | GO:0005886 | plasma membrane | $9.999 \times 10^{-5}$ |
| 8 | GO:CC | GO:0042995 | cell projection | $2.307 \times 10^{-4}$ |
| 9 | GO:CC | GO:0005871 | kinesin complex | $5.542 \times 10^{-4}$ |

**Cluster 4**

**version** e111_eg58_p18_f463989d

**date** 8/26/2024, 6:42:02 PM

**organism** hsapiens

g:Profiler

**Figure S8d**

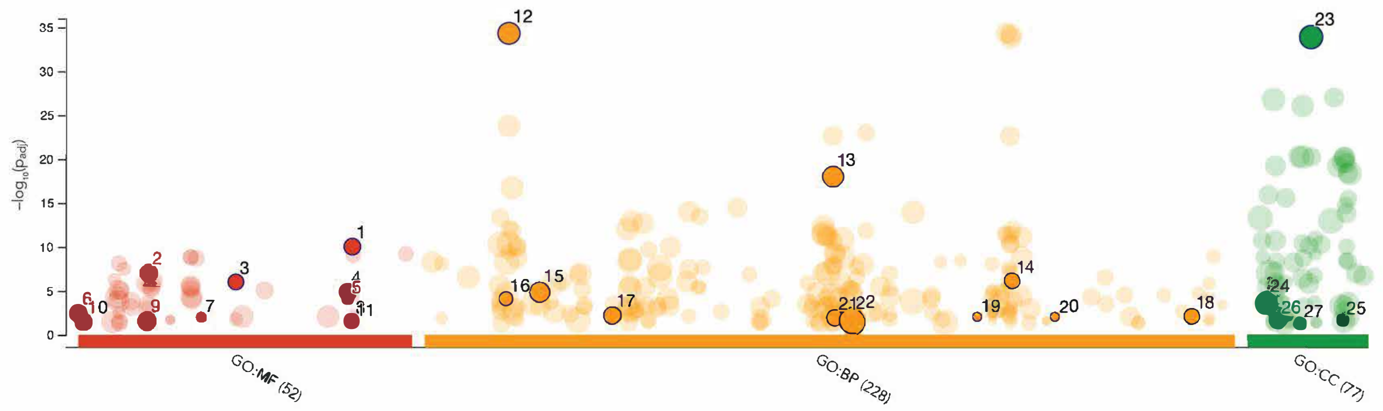


| ID | Source | Term ID | Term Name | $p_{adj}$ (query_1) |
|---|---|---|---|---|
| 1 | GO:MF | GO:0098960 | postsynaptic neurotransmitter receptor activity | $9.236\times10^{-11}$ |
| 2 | GO:MF | GO:0015171 | amino acid transmembrane transporter activity | $9.488\times10^{-8}$ |
| 3 | GO:MF | GO:0035254 | glutamate receptor binding | $9.607\times10^{-7}$ |
| 4 | GO:MF | GO:0097110 | scaffold protein binding | $1.283\times10^{-5}$ |
| 5 | GO:MF | GO:0097109 | neuroligin family protein binding | $8.249\times10^{-5}$ |
| 6 | GO:MF | GO:0000149 | SNARE binding | $3.793\times10^{-3}$ |
| 7 | GO:MF | GO:0031687 | A2A adenosine receptor binding | $1.002\times10^{-2}$ |
| 8 | GO:MF | GO:0098918 | structural constituent of synapse | $2.592\times10^{-2}$ |
| 9 | GO:MF | GO:0015079 | potassium ion transmembrane transporter activity | $2.984\times10^{-2}$ |
| 10 | GO:MF | GO:0001540 | amyloid-beta binding | $3.327\times10^{-2}$ |
| 11 | GO:MF | GO:0098919 | structural constituent of postsynaptic density | $3.330\times10^{-2}$ |
| 12 | GO:BP | GO:0007268 | chemical synaptic transmission | $4.534\times10^{-35}$ |
| 13 | GO:BP | GO:0050808 | synapse organization | $9.738\times10^{-19}$ |
| 14 | GO:BP | GO:0099601 | regulation of neurotransmitter receptor activity | $7.018\times10^{-7}$ |
| 15 | GO:BP | GO:0010038 | response to metal ion | $1.443\times10^{-5}$ |
| 16 | GO:BP | GO:0007158 | neuron cell-cell adhesion | $7.726\times10^{-5}$ |
| 17 | GO:BP | GO:0021510 | spinal cord development | $6.213\times10^{-3}$ |
| 18 | GO:BP | GO:1990090 | cellular response to nerve growth factor stimulus | $7.820\times10^{-3}$ |
| 19 | GO:BP | GO:0090125 | cell-cell adhesion involved in synapse maturation | $8.863\times10^{-3}$ |
| 20 | GO:BP | GO:1900451 | positive regulation of glutamate receptor signaling pathway | $8.863\times10^{-3}$ |
| 21 | GO:BP | GO:0050885 | neuromuscular process controlling balance | $1.354\times10^{-2}$ |
| 22 | GO:BP | GO:0051716 | cellular response to stimulus | $2.732\times10^{-2}$ |
| 23 | GO:CC | GO:0045202 | synapse | $1.159\times10^{-34}$ |
| 24 | GO:CC | GO:0009986 | cell surface | $2.371\times10^{-4}$ |
| 25 | GO:CC | GO:0098635 | protein complex involved in cell-cell adhesion | $2.087\times10^{-2}$ |
| 26 | GO:CC | GO:0031045 | dense core granule | $2.521\times10^{-2}$ |
| 27 | GO:CC | GO:0042584 | chromaffin granule membrane | $4.987\times10^{-2}$ |

**Cluster 5**

| | |
|---|---|
| **version** | e111_eg58_p18_f463989d |
| **date** | 8/26/2024, 6:42:28 PM |
| **organism** | hsapiens |

g:Profiler

**Figure S8e**

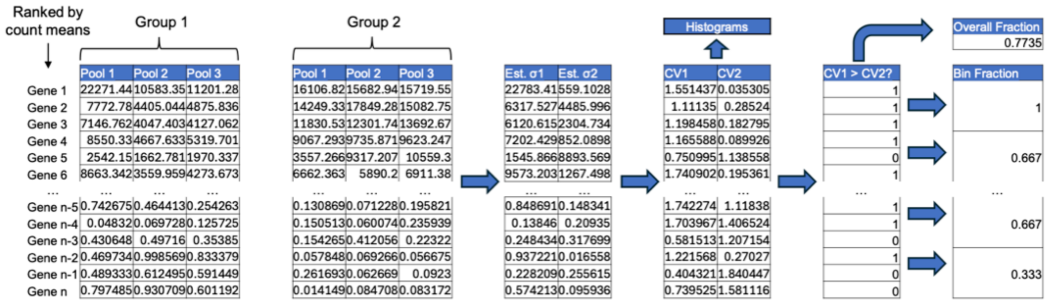

Ranked by count means
Group 1
Group 2
Pool 1 Pool 2 Pool 3
Pool 1 Pool 2 Pool 3
Gene 1 22271.44 10583.35 11201.28
Gene 2 7772.78 4405.044 4875.836
Gene 3 7146.762 4047.403 4127.062
Gene 4 8550.33 4667.633 5319.701
Gene 5 2542.15 1662.781 1970.337
Gene 6 8663.342 3559.959 4273.673
... ... ... ...
Gene n-5 0.742675 0.464413 0.254263
Gene n-4 0.04832 0.069728 0.125725
Gene n-3 0.430648 0.49716 0.35385
Gene n-2 0.469734 0.998569 0.833379
Gene n-1 0.489333 0.612495 0.591449
Gene n 0.797485 0.930709 0.601192
16106.82 15682.94 15719.55
14249.33 17849.28 15082.75
11830.53 12301.74 13692.67
9067.293 9735.871 9623.247
3557.266 9317.207 10559.3
6662.363 5890.2 6911.38
... ... ...
0.130869 0.071228 0.195821
0.150513 0.060074 0.235939
0.154265 0.412056 0.22322
0.057848 0.069266 0.056675
0.261693 0.062669 0.0923
0.014149 0.084708 0.083172
Est. σ1 Est. σ2
22783.41 559.1028
6317.527 4485.996
6120.615 2304.734
7202.429 852.0898
1545.866 8893.569
9573.203 1267.498
... ...
0.848691 0.148341
0.13846 0.20935
0.248434 0.317699
0.937221 0.016558
0.228209 0.255615
0.574213 0.095936
Histograms
CV1 CV2
1.551437 0.035305
1.11135 0.28524
1.198458 0.182795
1.165588 0.089926
0.750995 1.138558
1.740902 0.195361
... ...
1.742274 1.11838
1.703967 1.406524
0.581513 1.207154
1.221568 0.27027
0.404321 1.840447
0.739525 1.581116
CV1 > CV2?
1
1
1
1
0
1
...
1
1
0
1
0
0
Overall Fraction
0.7735
Bin Fraction
1
0.667
...
0.667
0.333


**Figure S9**

**Figure S1.** Comparison of physical and kinematic parameters between biobots and sham neurobots. Kruskal-Wallis test was used to obtain p-values.

**Figure S2.** Examples of Z-projected confocal fluorescent images of neurobots. Each row corresponds to an individual neurobot. The first two columns show staining of acetylated alpha tubulin, which labels multiciliated cells and neurons (color code represents depth in the confocal stack), with the first column depicting the full stack to show the distribution of the multiciliated cells on the surface, and the second column depicting a partial stack to reveal the interior of the neurobot where neurons reside. The last column shows a nuclear stain of the same bot, over the same partial stack as shown in the second column. All neurobots contain processes within the bot and those that extend towards the surface. They also contain a central region with seemingly no nuclei present, which we hypothesize may be filled with extracellular matrix materials.

**Figure S3.** Comparison of various kinematic parameters between biobots and neurobots. We found no significant difference between biobots and neurobots in total distance traveled, average speed, average acceleration, and percentage of the well area visited.

**Figure S4.** Neurobots had a significantly smaller density of multiciliated cells. Kruskal-Wallis test was used to obtain p-values.

**Figure S5.** Impact of treatment with zolmitriptan on neural expression patterns in neurobots.

**a.-c.** Treatment with zolmitriptan increased number of terminals, total length of neurites and neurite density. There was no significant change in complexity index (**d**). **e.** Pairwise correlation between structural parameters of zolmitriptan-treated neurobots and Complexity Index, $N_{terminals}$ = total number of endings, $L_{Neurite}$ = total length of neurites, $L_{Neuritenorm}$ = total length of neurites normalized to area, $N_{MCC}$ = total number of multiciliated cells on the top surface, $N_{MCCnorm}$ = $N_{MCC}$ normalized to area, RI = Roundness Index, Neu/Ect = ratio of the areas of neural implant to ectoderm shell.

**Figure S6.** Enrichment analysis performed using Gene Ontology annotations **a**. sham neurobots vs biobots (4-fold upregulated pathways) and **b.** neurobots vs biobots (4- fold downregulated pathways)

**Figure S7.** Enriched pathways in neurobots when compared to biobots, across clusters identified based on network analysis.

**Figure S8.** Enriched pathways in sham neurobots when compared to biobots, across clusters identified based on network analysis.

**Figure S9.** Method for quantifying variability in gene expression. For a chosen pair of groups, genes were ranked by the mean count value across all pools of both groups, and the CV of each gene's counts across the pools of each group was calculated. The CV list was split into 100 bins (percentiles) containing equal numbers of genes, and the fraction of genes in the bin for which the CV of the first group was greater than that of the second group was found and plotted.
